# Elucidating human gut microbiota interactions that robustly inhibit diverse *Clostridioides difficile* strains across different nutrient landscapes

**DOI:** 10.1101/2024.04.13.589383

**Authors:** Jordy Evan Sulaiman, Jaron Thompson, Yili Qian, Eugenio I. Vivas, Christian Diener, Sean M. Gibbons, Nasia Safdar, Ophelia S. Venturelli

## Abstract

The human gut pathogen *Clostridioides difficile* displays extreme genetic variability and confronts a changeable nutrient landscape in the gut. We mapped gut microbiota inter-species interactions impacting the growth and toxin production of diverse *C. difficile* strains in different nutrient environments. Although negative interactions impacting *C. difficile* are prevalent in environments promoting resource competition, they are sparse in an environment containing *C. difficile*-preferred carbohydrates. *C. difficile* strains display differences in interactions with *Clostridium scindens* and the ability to compete for proline. *C. difficile* toxin production displays substantial community-context dependent variation and does not trend with growth-mediated inter-species interactions. *C. difficile* shows substantial differences in transcriptional profiles in the presence of the closely related species *C. hiranonis* or *C. scindens*. In co-culture with *C. hiranonis*, *C. difficile* exhibits massive alterations in metabolism and other cellular processes, consistent with their high metabolic overlap. Further, *Clostridium hiranonis* inhibits the growth and toxin production of diverse *C. difficile* strains across different nutrient environments and ameliorates the disease severity of a *C. difficile* challenge in a murine model. In sum, strain-level variability and nutrient environments are major variables shaping gut microbiota interactions with *C. difficile*.

## INTRODUCTION

The human gut microbiome exists in a dynamic balance between homeostasis and disruption due to the contrasting evolutionary objectives of the host and the resident gut bacteria. *Clostridioides difficile* is an opportunistic human gut pathogen that can cause life-threatening damage to the colon. Antibiotics are the first-line treatment for *C. difficile* infection (CDI). However, they also damage the commensal gut microbiota that provides *C. difficile* colonization resistance and could cause the recurrence of CDI (rCDI) ^1–3^. Fecal microbiota transplantation (FMT) has proven to be effective for treating rCDI, but the effect of FMT on a patient can vary due to uncharacterized factors and donor microbiota variability ^4^. FMT can also result in the unintentional transfer of antibiotic-resistant bacteria, including other opportunistic pathogens ^5,6^. To overcome these limitations, defined communities of commensal bacteria can be designed to inhibit *C. difficile*. However, low richness communities do not display robustness of anti*-C. difficile* activity to changes in environmental contexts ^7,8^. This in turn could contribute to the variability in efficacy in clinical trials of certain living bacterial therapeutics for treating CDI ^9^. We lack an understanding of how environmental context, such as the genetics of *C. difficile* strains and nutrient environments, impacts the anti-*C. difficile* activity of human gut communities^10,11^.

*C. difficile* has a diverse population structure comprising hundreds of strain types ^12^ that are distributed across at least 8 phylogenetic clades ^13^. This species is defined by a large pangenome ^14^, with an ultralow core genome (as low as 16% based on 73 genomes ^15^) and extreme levels of evolutionary plasticity that have been molded over long periods through frequent exchange with bacterial gene pools in multiple host environments via horizontal gene transfer ^16–19^. This substantial genetic variation among *C. difficile* strains has downstream impacts on the regulation of metabolic pathways and virulence ^16,20–22^. For instance, the emergence of the hypervirulent epidemic strain ribotype 027 has been proposed as the major driver of the increase in the prevalence of CDI ^23,24^. Notably, rCDI is not always due to infection with the same strain, where new strains were observed in 33-56% of recurrent episodes ^25–29^. This suggests that the degree of colonization resistance could vary across different *C. difficile* strains, potentially leading to differences in patient outcomes.

Interactions with gut microbiota are critical determinants of *C. difficile* colonization and toxin production, as evidenced by the colonization resistance variability of different microbiome compositions to *C. difficile*^30^. Previous studies have elucidated principles that influence *C. difficile* growth in human gut communities *in vitro*, such as a strong negative dependence on species richness ^31^, and identified specific mechanisms of *C. difficile* inhibition. For example, certain species compete with *C. difficile* for limiting resources, such as the consumption of specific mucus-derived sugars by *Akkermansia muciniphila*^32^ or the utilization of Stickland metabolism amino acids by *Clostridium* species (e.g. *Clostridium bifermentans* ^33,34^ and *Clostridium scindens* ^33^). In addition, *C. scindens* can produce tryptophan-derived antibiotics that inhibit *C. difficile* growth ^35^. *Clostridium hiranonis* was shown to inhibit *C. difficile in vitro* through more than one mechanism in a single nutrient environment ^31^. However, the contribution of *C. difficile* strain-level variability to these interactions is currently unknown ^31,36,37^.

The bottom-up construction of synthetic microbiomes combined with computational modeling ^38,39^ and principled experimental design techniques ^40^ can be used to efficiently navigate large design landscapes of combinations of species. In addition, these bottom-up approaches can provide a deeper understanding of important molecular and ecological mechanisms. For example, a widely used dynamic ecological model referred to as generalized Lotka–Volterra (gLV) can be used to unravel growth-mediated microbial interactions shaping community assembly ^41–43^. By informing the model with properly collected experimental data, the gLV model can accurately forecast community dynamics as a function of the intrinsic growth of individual species and pairwise interactions with all constituent community members ^38,44^.

To understand how nutrient and strain-level variability shapes interaction networks with *C. difficile*, we used a bottom-up approach to build microbial communities combined with computational modeling. We elucidated strain-level differences in inter-species interactions at the transcriptional level using genome-wide transcriptional profiling. In addition, we discovered that the large variation in toxin production of *C. difficile* in communities was not correlated with growth-mediated inter-species interactions. Our workflow identifies *Clostridium hiranonis* as a “universal” *C. difficile* growth and toxin production inhibitor that is robust to variation in strain backgrounds and nutrient environments. This robust inhibition is consistent with its high metabolic niche overlap with *C. difficile*, which in turn could block the utilization of *C. difficile*-preferred substrates.

Consistent with this notion, genome-wide transcriptional profiling reveals a unique massive alteration of *C. difficile* metabolism in the presence of *C. hiranonis*, which is not observed in co-culture with another closely related species, *C. scindens*. Furthermore, *C. hiranonis* ameliorated the *C. difficile-*induced disease severity of mice due to reduced abundance and toxin production. In sum, we demonstrate that strain-level variability and nutrient environments play an important role in shaping the interactions between *C. difficile* and human gut communities, and highlight *C. hiranonis* as a promising candidate to include in the design of robust anti-*C. difficile* defined consortia.

## RESULTS

### C. difficile strains display substantial phenotypic and genetic variability

To understand how the strain-level genetic variability influences *C. difficile* phenotypes, we characterized 18 *C. difficile* strains (9 from diseased patients that were diagnosed and treated for CDI and 9 from healthy individuals) and *C. difficile* DSM 27147 (R20291 reference strain of the epidemic ribotype 027). We individually profiled their growth in a chemically defined media supplemented with carbohydrate sources shown to promote colonization or virulence activities including succinate ^45,46^, trehalose ^21,22^, mannitol ^46,47^, sorbitol ^46,47^, and various mucus-derived sugars such as sialic acid and *n*-acetyl-D-glucosamine ^36,48^ (**Fig. S1a-d**; **Table S1, 2**). The growth of all *C. difficile* strains was supported in defined media without any carbohydrate source due to their ability to utilize amino acids through Stickland metabolism. In general, supplementation of glucose, mannitol, *n*-acetyl-D-glucosamine (GlcNAc), and sialic acid enhanced the growth of all *C. difficile* strains compared to media without carbohydrate sources.

In most single carbohydrate media, *C. difficile* displayed a unique growth profile that is distinct from the other commensal gut bacteria, where the culture grew rapidly at the beginning followed by a steep decline in OD_600_ during stationary phase at ∼24 h of growth (i.e. non-monotonic growth response). While the variance in monoculture growth biological replicates is low in the first 24 h, this variability increases substantially at the time when OD_600_ declines in stationary phase (**Fig. S1g-h**). This implies that sporulation and cell lysis, in addition to halted cell division as observed by fluorescence microscopy contribute to the observed reduction and variability in OD_600_ (**Fig. S2**). To quantify the variability in growth profiles across *C. difficile* strains, we fit each growth curve to a logistic model to determine the growth rate (*r*) and carrying capacity (*K*) of each strain excluding data points with a >10% reduction in OD_600_ in the late stationary phase (**Fig. 1a-b, S1e**, see **Methods**). Overall, the logistic model displayed a high goodness of fit to the data (Pearson R=0.98, P<10E-05) (**Fig. S1f**).

**Figure 1.**
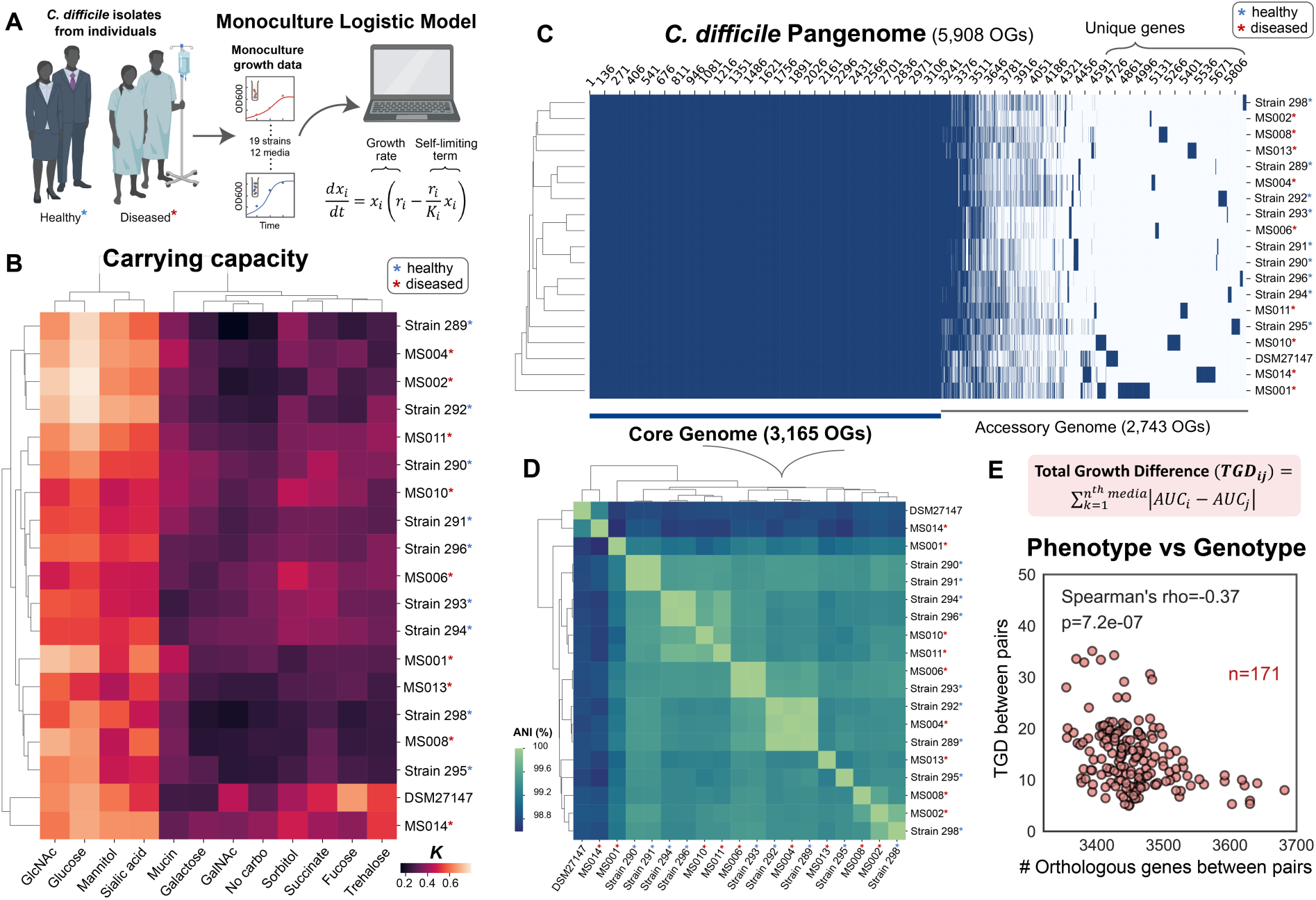
Phenotypic and genotypic characterization of diverse *C. difficile* isolates from diseased and healthy individuals. **a,** Fitting of monoculture growth data from 19 *C. difficile* strains (including 18 *C. difficile* isolates from diseased and healthy individuals, **Table S1**) in 12 media containing different carbohydrate sources (**Fig. S1a-b**) to the logistic model. Mathematical description of the logistic growth model was shown, where *x_i_* is the absolute abundance of species *i*, parameter *r_i_* is its maximum growth rate, and *K_i_* is the carrying capacity. When fitting the experimental data to the model, we cut time points where OD_600_ drops above 10% to exclude the highly variable phase. **b**, Biclustering heatmap of the carrying capacity (*K_i_*) of the *C. difficile* isolates. Strains marked with red asterisks were isolated from diseased patients whereas the ones marked with blue asterisks were isolated from healthy individuals. **c**, Heatmap showing the presence and absence of all genes identified across the 19 *C. difficile* strains (pangenome). The columns indicate the genes, and the rows indicate the *C. difficile* strains. Blue means gene present and white means gene absent. Genes present in all of the 19 strains are the core genome, whereas genes present in a subset of the strains are the accessory genome. **d**, Biclustering heatmap of the Average Nucleotide Identity (ANI) of *C. difficile* isolate pairs based on their whole-genome sequence. The horizontal boxes indicate 100% ANI. **e**, Scatter plot of the Total Growth Difference (TGD) between isolate pairs and the number of orthologous genes between isolate pairs. Mathematical formula to calculate the TGD between isolate pairs is shown on the top, which is the sum of all AUC differences from 24 h of growth in the twelve media. Parts of the figure are generated using Biorender.

We performed whole-genome sequencing on each isolate to provide insights into the genetic variation driving the observed phenotypic variability. The *C. difficile* genome is comprised of a well-conserved core genome (3,165 orthologous genes) and substantial variation in its accessory genome (**Fig. 1c**). Metabolic genes varied substantially across the 19 *C. difficile* strains, where only ∼63% of metabolic genes were shared (**Fig. S3a**). Clustering based on *ANI*, which represents the *A*verage *N*ucleotide *I*dentity of all orthologous genes shared between any two genomes, highlighted strains that are more genetically similar to each other (**Fig. 1d**), such as DSM 27147 and MS014. In addition to other genome similarities, these two strains possessed a mutation in the *treR* gene (L172I) that confers enhanced trehalose metabolism identified in hypervirulent *C. difficile* strains ^22^, consistent with their higher capability to utilize trehalose (**Fig. 1b**). Further, MS001 is clustered separately from the rest of the group based on ANI. MS001 has a much higher number of genes (4110) compared to the other strains (ranging from 3629 to 3892) (**Table S3**), and uniquely lacks the toxins TcdA and TcdB. Indeed, non-toxigenic *C. difficile* strains have distinct phenotypes compared to toxigenic strains, as a consequence of the variability in their genome ^49^. In general, there is no pattern between the *C. difficile* isolates from healthy and sick individuals in terms of their genotype.

To quantify if the genotypic variation displays an informative relationship with phenotypic variation in monoculture, we define the growth difference (GD) as the absolute value of the difference in the AUC of pairs of strains in a specific media. The total growth difference (TGD) is the sum of GD across the 12 media. The TGD and the number of orthologous genes (OGs) or ANI of pairs of *C. difficile* strains displayed a moderate negative correlation (**Fig. 1e, S4a-b**). In addition, growth in glucose, trehalose, galactose, and sorbitol was negatively correlated with ANI and the number of OGs (**Fig. S4c-d**). These results suggest that the genotypic variability quantified by these metrics displays an informative relationship with the utilization of certain carbohydrates.

Although the number of genes responsible for most core processes beyond metabolism is similar across isolates, there was large variability in the number of genes related to DNA recombination and integration, which are markers of mobile genetic elements (MGEs) (**Fig. S3e**). This suggests that MGEs play a major role in driving *C. difficile* genotypic differences, consistent with previous reports ^50,51^. To characterize the contribution of plasmids to the genome of *C. difficile*, we searched for high-coverage contigs within genome assemblies and discovered 11 of such instances in 7 of 19 genomes (**Fig. S5a-c**). These putative plasmids contained direct repeats on their termini indicative of being circular. In addition, the putative plasmids do not contain genes that could provide a selective advantage to these strains such as antibiotic resistance or virulence factors (**Table S5**). Interestingly, 4 of the 11 high-coverage contigs map to the same plasmid that is present in four different genetically distant *C. difficile* isolates from different patients. These isolates also have a highly variable number of conjugative systems and phages, covering 1.4-16.5% of their genomes (**Fig. S3f, Table S6-7**). In sum, the *C. difficile* isolates have highly diverse genomes with substantial variability in metabolic genes and mobile genetic elements.

### Human gut communities containing different C. difficile isolates display differences in interaction networks

Since human gut microbiota interactions are critical determinants of *C. difficile* growth and colonization, we investigated how *C. difficile* genetic variation shapes gut microbiota interspecies interactions. To this end, we built human gut communities from the bottom up with one of 4 diverse *C. difficile* strains (DSM 27147, MS001, MS008, and MS014) and combinations of 7 gut species (*C. scindens* (CS), *C. hiranonis* (CH), *Desulfovibrio piger* (DP), *Bacteroidetes thetaiotaomicron* (BT), *Phocaeicola vulgatus* (PV), *Bacteroidetes uniformis* (BU), and *Collinsella aerofaciens* (CA)) (**Fig. 2a**). Many of these species are prevalent across individuals and span major phyla of the human gut microbiome. These species displayed variation in growth in media with different carbohydrates (**Fig. S1a-b**). The community features CS, previously shown to inhibit the growth of *C. difficile* in gnotobiotic mice ^37^, CH which can inhibit *C. difficile* growth through unknown mechanism ^31^, and *Bacteroides* species, which have the potential for *C. difficile* inhibition in different environments ^36,45,52,53^.

**Figure 2.**
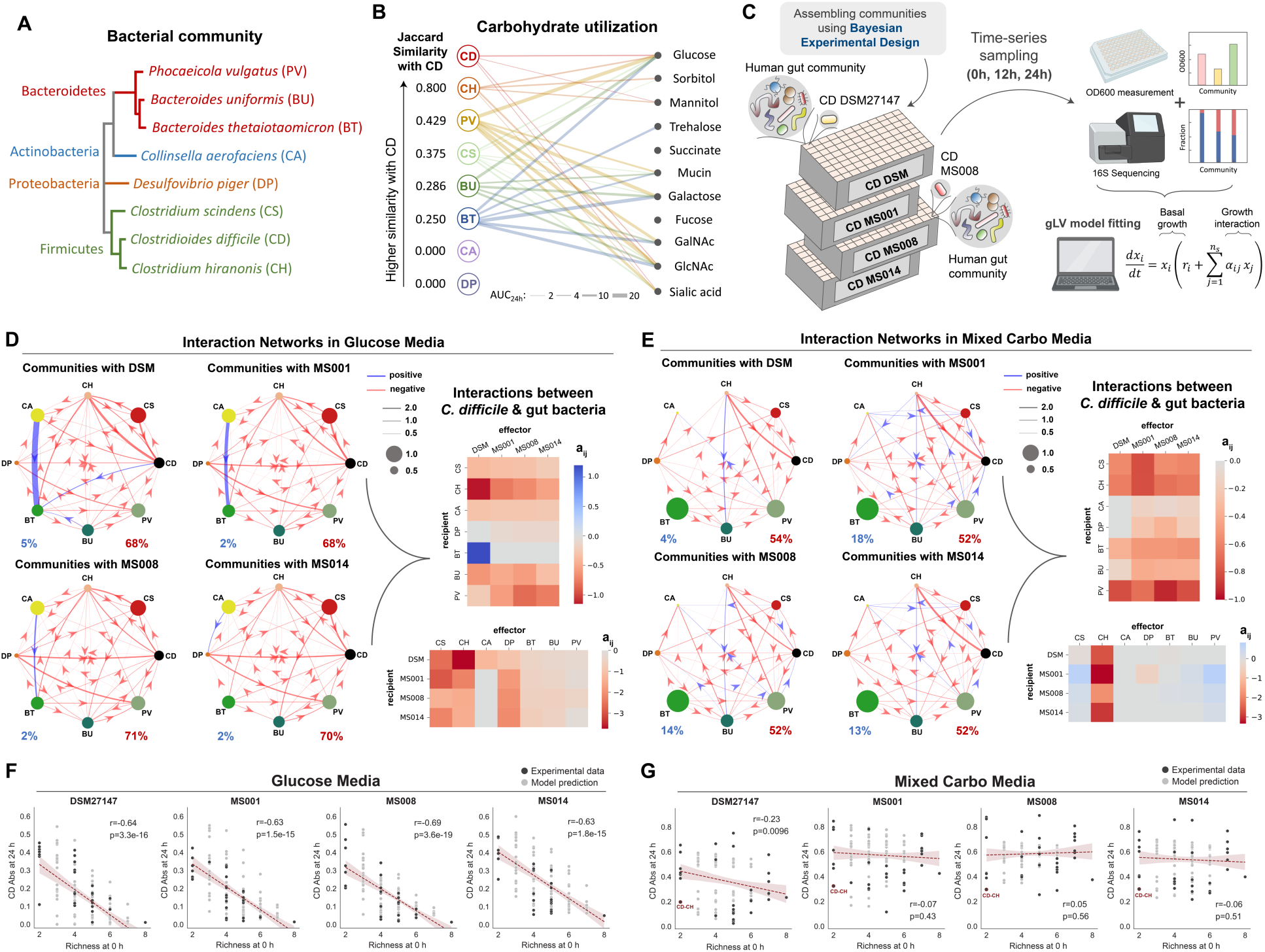
Interspecies interactions between *C. difficile* strains and the human gut bacteria in different nutrient environments. **a**, Phylogenetic tree of the 7-member resident synthetic gut community and *C. difficile*. The phylogenetic tree was generated from the 16S rRNA sequence of each species using the Clustal Omega multiple sequence alignment tool. **b**, Bipartite network of carbohydrate utilization by *C. difficile* and gut bacteria based on their monoculture growth profiles in **Fig. S1a-b**. The edge thickness indicates the AUC24h of the gut species grown in specific carbohydrates subtracted by the AUC24h of the gut species grown in media without any carbohydrates. Only edges with a magnitude larger than 2 are shown. For *C. difficile*, the growth profile of the DSM27147 strain is used as a representative. The Jaccard Similarity values of each gut species with *C. difficile* were computed based on the number of carbohydrates being utilized, where higher Jaccard Similarity values mean larger niche overlap with *C. difficile*. Different colors represent different species. **c**, Schematic of the experimental workflow to assess interactions between different *C. difficile* strains and human gut bacteria in the glucose media. Experimental communities were assembled using the Bayesian experimental design by utilizing monoculture growth data as prior information (See **Methods**). A total of 147 subcommunities (2 to 8 species) containing combinations of gut species and one of the *C. difficile* strains were cultured at an equal absolute abundance ratio in the glucose media. Cultures were grown in microtiter plates in anaerobic conditions and incubated at 37°C. After 12 h and 24 h of growth, aliquots of the culture were taken for multiplexed 16S rRNA sequencing to determine community composition and cell density measurement at 600 nm (OD_600_) to calculate the absolute abundance of each species. Absolute abundance data are used to infer the parameters of a generalized Lotka–Volterra (gLV) model and elucidate the interaction networks of the communities. **d-e**, Inferred interspecies interaction networks between the 7 gut species and each of the representative *C. difficile* strains when grown in the glucose media (**d**) or the mixed carbohydrates media (**e**). Node size represents species carrying capacity in monoculture (mean of all biological replicates) and edge width represents the magnitude of the interspecies interaction coefficient (aij). Edges represent parameters whose absolute values were significantly constrained to be non-zero based on the Wald test (**Fig. S8** for glucose media and **Fig. S10** for mixed carbohydrates media). Percentage of positive (blue) and negative (red) interactions for each community are shown. The right panel shows the heatmap of interspecies interaction coefficients of the gLV model between the different *C. difficile* strains and the 7 gut species in the glucose media (**d**) or the mixed carbohydrates media (**e**). **f-g,** Scatter plots of *C. difficile* absolute abundance at 24 h as a function of initial species richness in all possible subcommunities of 2–8 species simulated by the gLV (gray data points) and in experimentally measured subcommunities (mean value of biological replicates, black data points). **Panel f** are model predictions and experimental data of communities grown in the glucose media, whereas **Panel g** are those grown in the mixed carbohydrates media. Red dashed line indicates the linear regression between the species richness at 0 h and *C. difficile* absolute abundance at 24 h, with the 95% confidence bounds shown as red shading. Pearson’s correlation coefficient (*r*) and *p*-values are shown, which were computed using the pearsonr from the scipy package in Python. Parts of the figure are generated using Biorender.

To infer the inter-species interaction networks, we down selected a set of representative *C. difficile* strains based on their genotypic and phenotypic variations. Strains that have similar genotypes and metabolic genes may display similar interaction networks, whereas interactions may be divergent for strains with large differences in genotype. MS014 shows a similar genotype to DSM 27147 and thus might evolve from the same ancestor, but MS014 was more recently isolated. By contrast, the non-toxigenic strain MS001 has the most different genotype than the other strains, suggesting potentially larger differences in inter-species interactions. Finally, MS008 is genotypically and phenotypically distinct from the other 3 strains (**Fig. 1b-d**). In addition, MS008 clustered differently from MS014, DSM 27147 and MS001 based on metabolic genes, suggesting divergent metabolic capabilities (**Fig. S3a**).

Given the key role of resource competition in the ecology of *C. difficile* ^32–34,54^, the extent of metabolic niche overlap with *C. difficile* may be a major variable influencing interactions with human gut bacteria. To quantify the extent of metabolic niche overlap between each gut species and *C. difficile*, we calculated the Jaccard Similarity of carbohydrate utilization based on the change in growth in the presence and absence of the given carbohydrate (**Fig. 2b**). Notably, CH displayed the largest metabolic niche overlap of carbohydrate utilization with *C. difficile* (Jaccard Index=0.8). In addition to the similarities in carbohydrate utilization, CH has been shown to use amino acids via Stickland metabolism, similar to *C. difficile* ^33^.

To study community inter-species interactions in a gut environment with high resource competition, we used a defined media containing glucose as the sole carbohydrate source. Glucose can support the growth of most species in monoculture including *C. difficile,* and thus promotes inter-species competition (**Fig. S6a**-**c**). To quantify the differences in the inter-species interaction networks, we cultured different combinations of species with one of the four *C. difficile* strains (DSM 27147, MS001, MS008, and MS014) (**Fig. 2c**). Since there are too many community combinations to be comprehensively explored (635 combinations), we used a Bayesian experimental design approach to select combinations of bacteria that would maximize information content as quantified by the expected Kullback-Leibler divergence between the posterior and prior parameter distributions (see **Methods** and **Supplementary text**) ^40^. Briefly, a preliminary gLV model was fit to the monoculture growth in glucose media. We used a Bayesian inference approach to approximate the posterior parameter distribution as a multivariate Gaussian. The parameter distribution inferred for the preliminary model was used as a prior to guide the design of 147 combinations of 2 to 8-member sub-communities containing one of the four *C. difficile* strains (DSM 27147, MS001, MS008, and MS014). Species absolute abundance was determined by multiplying the relative abundance fraction via multiplexed 16S rRNA sequencing by the total biomass obtained by OD_600_ as previously described ^31,38^. The parameters of the gLV model were inferred based on time-series data of species abundances (0, 12, and 24 h) (**Fig. S7a**, DATASET001 in **Table S8**). Based on the parameter posterior distributions, we analyzed parameters with absolute values that were significantly constrained to be non-zero based on the Wald test^55^ (**Fig. S8**, **Supplementary text**). The Wald test compares the parameter mean to its standard deviation to evaluate whether the peak of the posterior parameter distribution is significantly higher or lower than zero compared to the width of the distribution. The percentage of constrained parameters is 76.6%, 73.4%, 75%, and 75% for communities containing DSM, MS001, MS008, or MS014 respectively. To evaluate model prediction performance on held-out data, we performed 10-fold cross-validation where only community samples were subjected to testing (see **Methods**). Using a 10-fold cross-validation, the model prediction exhibited good agreement with the measured species abundance in all communities with different *C. difficile* strains (Pearson’s R=0.93-0.95, P<10E-05), demonstrating that our model can capture and predict the trends in species abundance (**Fig. S7b**).

Consistent with a high competition resource environment, the interaction networks for distinct *C. difficile* strains displayed a high fraction of negative interactions (68-71%) and inhibition of *C. difficile* by all species (**Fig. 2d**). CS and CH display a high magnitude of negative inhibition towards *C. difficile*, consistent with their ability to compete for amino acids via Stickland fermentation. Notably, the *C. difficile* DSM 27147 hypervirulent strain exhibits the largest differences in interaction profile from other *C. difficile* strains (e.g. BT, DP, and CH).

In addition to the observed changes in pairwise interactions with *C. difficile*, other inter-species interactions displayed strain-specific differences. A higher order interaction (HOI) is defined as a substantial change in a pairwise interaction due to the presence of a third community member ^56,57^. Changes in pairwise interactions due to the presence of different *C. difficile* strains may suggest HOI. For instance, the interaction coefficients between CA and BT are substantially impacted by the specific *C. difficile* strain that is present in the community (**Fig. S7c**). To further explore whether *C. difficile* strain variations could impact CA-BT interactions, we cultured the CA-BT pairwise community in the sterilized spent media of *C. difficile* (**Fig. S7d**). The abundances of CA and BT in the community were statistically different when cultured in the sterile conditioned media of the different *C. difficile* strains. This implies that different strains of *C. difficile* differentially altered the chemical environment, which in turn impacted the interactions between CA and BT. In sum, inferred inter-species interaction networks containing distinct *C. difficile* strains displayed infrequent direct and indirect differences.

### Human gut bacteria infrequently inhibit C. difficile in the presence of preferred carbohydrates

Antibiotic treatments lead to massive gut bacterial mortality, alternations in the resource landscape, and changes in community composition. This new environment can be exploited by *C. difficile* ^45,48,58–61^. To explore community interactions in media that mirrors post-antibiotic environments, we designed a media containing multiple carbohydrates that could be utilized by *C. difficile* (mixed carbohydrates media) (**Fig. S9a**). In this media, *C. difficile* strains displayed substantial growth and a diminished decline in OD_600_ in late stationary phase than glucose media (**Fig. S9b**). In pairwise communities, the relative abundance of *C. difficile* was high in all communities (>50% in all cases) except when grown with BT. The absolute abundance of *C. difficile* remained high after three 24 h growth cycles, except for the community containing PV (**Fig. S9c-d**). In the 7-member community, *C. difficile* displayed a relative abundance of ∼20-50% following 24 h of growth (**Fig. S9e**). This contrasts with the low abundance *of C. difficile* in the glucose media (∼1 to 5%) (**Fig. S7a**).

To determine the inter-species interaction network in the presence of multiple preferred carbohydrates, we built a gLV model using a design-test-learn (DTL) cycle (**Fig. S9f**). A DTL cycle was used to account for potentially more complex interactions in the presence of a complex resource environment, which may require additional data to constrain the model parameters. Each cycle consisted of (i) Bayesian experimental design informed by prior experimental observations to select combinations of species that minimize parameter uncertainty (design), (ii) experimental characterization of sub-communities (test), and (iii) updates to the gLV model parameters based on new experimental data (learn) (**Methods** and **Supplementary text**) ^44^. In the initial experiment, we constructed 82 communities consisting of all possible pairwise, leave-one-out, and full communities containing the gut bacteria and individual *C. difficile* strains (**Table S8**, DATASET002). Using 10-fold cross-validation, the model displayed a low to moderate prediction performance of individual species (**Fig. S9g**). To select informative experimental conditions for the second DTL cycle, Bayesian experimental design based on the inferred parameter uncertainties guided the design of 94 new combinations of medium richness communities (3-6 members) (**Table S8**, DATASET003). Using these data, the prediction performance of most individual species was improved (Pearson’s R=0.90 to 0.91, P<10E-05) (**Fig. S9g**). The parameter uncertainty distributions are shown in **Fig. S10**. In comparison to the media with glucose, the constrained non-zero parameters are lower in the mixed carbohydrates media (60.9%, 71.8%, 68.8%, and 67.2% for communities containing DSM, MS001, MS008, and MS014 respectively). To determine whether species predictive performance could be improved with additional data, we performed a sensitivity analysis of the model’s prediction performance by varying how the training and validation data was partitioned (*k* in k-fold) (**Fig. S11**). The model prediction performance increased with *k* and saturated for most species. This implies that additional data for moderately predicted species (e.g. CH and DP) will not substantially improve the model prediction performance. Poor or moderate prediction performance could be due to insufficient variation of the particular species abundance across communities or limited flexibility of the gLV model to capture complex interaction modalities ^39^.

The inferred interaction networks in the mixed carbohydrates media display a higher frequency of positive interactions (4-18%) compared to media containing only glucose (2-5%) (**Fig. 2e**), and *C. difficile* displayed higher absolute abundance across communities (**Fig. S12**). While DSM 27147 exhibited the most different interaction profile in glucose media, this strain displayed similar interaction patterns to MS008 and MS014 in the mixed carbohydrates media. By contrast, MS001 displayed the largest differences in inter-species interactions in the mixed carbohydrates media than the other *C. difficile* strains. Thus, the differential interaction profiles between the *C. difficile* strains and human gut microbiota are nutrient dependent. Of 7 diverse human gut species, only CH displayed negative interactions with each *C. difficile* strain. Several communities used to train the model (3-6 members) containing CH displayed a higher magnitude of *C. difficile* inhibition than the *C. difficile*–CH pairwise community (**Fig. 2g, S13a**). In particular, CS, DP, CA, and PV are enriched in these communities. This suggests that the inhibitory activity of CH can be further enhanced by the presence of specific gut bacteria.

To further investigate inter-species interactions in the mixed carbohydrate media, we cultured different *C. difficile* strains in the sterilized spent media of the gut bacteria and fresh media as a control (**Fig. S14a-b**). Overall, the qualitative effects of the pH-adjusted conditioned media were largely consistent with the signs of the inferred gLV pairwise interaction coefficients (71% agreement compared to 32% in the non-pH-adjusted conditioned media) (**Fig. S14c**). Without pH adjustment, *C. difficile* growth was substantially reduced in *Bacteroides spp.* conditioned media due to the acidification of the environment (pH of 5.0-5.2), and this inhibition was eliminated in the pH-adjusted *Bacteroides spp.* conditions. Since pH changes over time in co-culture, the large variation in the initial pH of the spent media may not be physiologically relevant to microbial community interactions. Notably, *C. difficile* growth was reduced in CS-conditioned media but not in co-culture with CS. This inconsistency suggests that the feedback of metabolite exchange and/or metabolic niche partitioning plays a role in the *C. difficile*-CS pair in the mixed carbohydrates media. Although CS can utilize many of the same carbohydrates as *C. difficile*, CS has a wider range of carbohydrate utilization capabilities than *C. difficile* in the tested media (**Fig. 2b**). This implies that *C. difficile* and CS may prefer utilizing similar resources in monoculture and display distinct metabolic niches in co-culture.

### Model accurately predicts C. difficile inhibition potential in human gut communities

Using the model trained on all data, we forecasted the abundance of *C. difficile* at 24 h in all possible communities (**Fig. 2f-g**). A previous study showed a strong negative dependence between *C. difficile* growth and species richness in a rich media ^31^, consistent with a negative relationship between these variables in glucose media. However, this trend was not present in the presence of mixed carbohydrates. This suggests that high-richness communities may not universally inhibit *C. difficile* in environments with *C. difficile* preferred substrates, and the identity of the species in the community may be more impactful than the number of species.

To determine if our model could design communities to inhibit *C. difficile*, we used our gLV model trained on community data in the mixed carbohydrates media (**Table S8**, DATASET003) to predict *C. difficile* abundance in all possible 2 to 8-member communities (**Fig. S15a**). Based on the model prediction, we selected a 3-member weak inhibitory community (WIC, consisting of BU, CA, and DP) and a strong inhibitory community (SIC, consisting of CH, CS, and DP). The WIC was selected due to its low inhibition potential of *C. difficile*, whereas the SIC was selected for its high inhibition potential against diverse *C. difficile* strains. Although CH was the only species that could strongly inhibit *C. difficile* in the mixed carbohydrates media, CH, CS, and DP were the three most inhibitory species in the glucose media (**Fig. 2d-e**). The interaction networks revealed sparse and almost negligible incoming negative interactions towards *C. difficile* in the WIC. By contrast, the SIC displayed stronger negative interactions towards *C. difficile*, especially from CH (**Fig. S15b**). To validate the model predictions, we cultured WIC and SIC in the absence and presence of different *C. difficile* strains (**Fig. S15c**). We observed that the abundance of all *C. difficile* strains in SIC was significantly lower than those in the WIC (∼2.1 to 4.2-fold lower), corroborating the differential inhibitory potential of the SIC and WIC communities and highlighting that the inhibition of the SIC is robust to strain-level variability. This indicates that the model could predict the *C. difficile* inhibition potential of different communities.

### C. difficile strains have a differential ability to compete with C. scindens over proline

Although CS can inhibit the growth of *C. difficile* via competition for limiting pools of amino acids via Stickland metabolism ^33^, inhibition of most *C. difficile* strains by CS was not observed in the mixed carbohydrates media (**Fig. 2e**). This suggests that these *C. difficile* strains occupied alternative metabolic niches in co-culture with CS. The inferred interaction from CS to MS001 was larger in magnitude than to MS008 or MS014. By contrast, CS moderately inhibited the growth of the DSM strain. Model predictions of co-cultures of CS and individual *C. difficile* strains displayed consistent trends with independent *in vitro* experiments that did not inform the gLV model (**Fig. 3a**).

**Figure 3.**
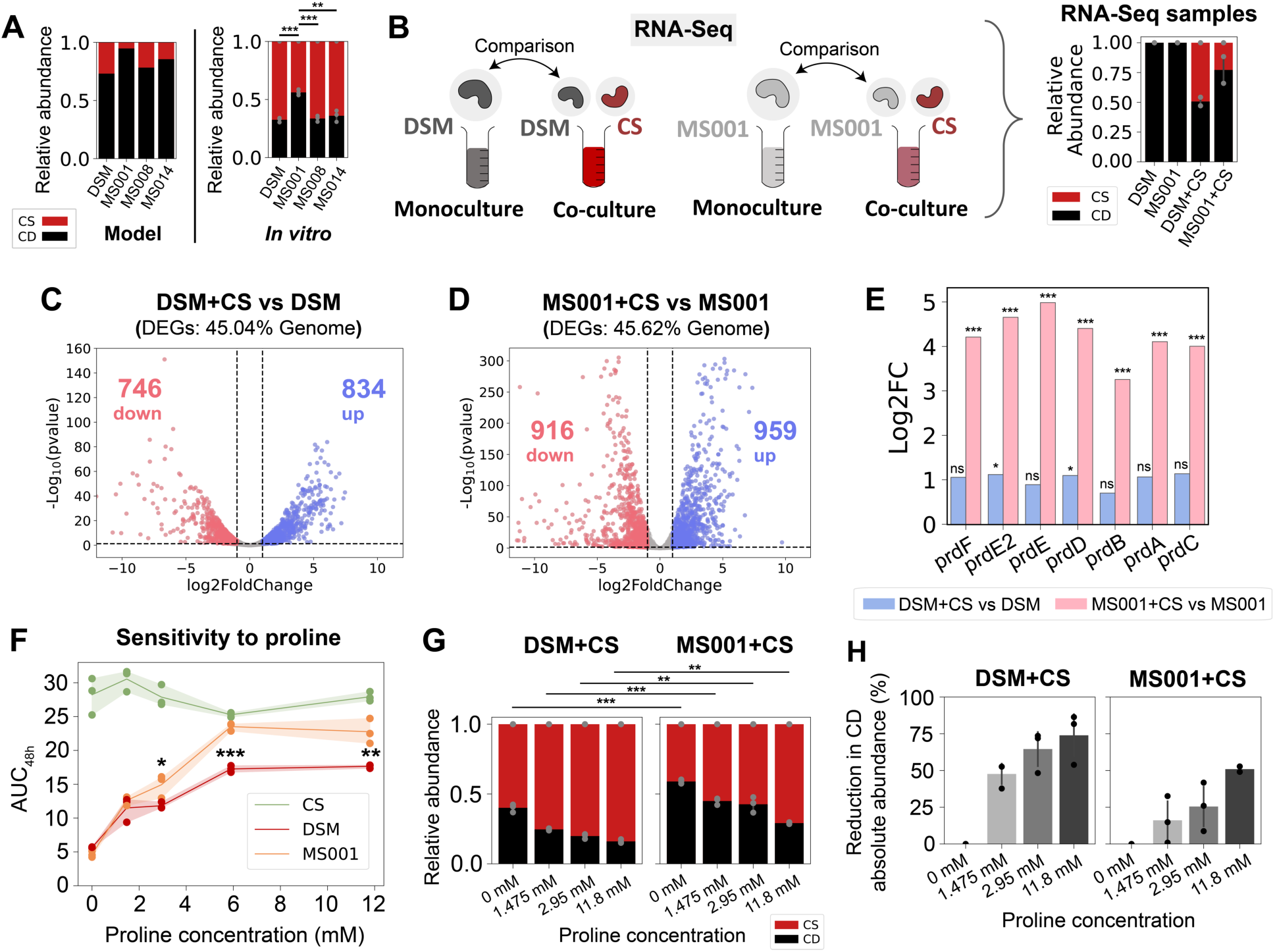
Genome-wide transcriptional profiling of *C. difficile* DSM27147 and *C. difficile* MS001 in the presence of *C. scindens*. **a**, Model prediction and independent experimental validation (not included in model fitting) of the relative abundance of pairwise communities containing CS and one of the four *C. difficile* strains. Each bar represents the average absolute abundance of each species, and the error bars on the *in vitro* data represent s.d. (n=3). Asterisks above the bars indicate the *p*-value from unpaired *t*-test of species relative abundance between co-cultures: ** indicates p<0.01, *** indicates p<0.001. **b**, Schematic of the genome-wide transcriptional profiling experiment of two *C. difficile* strains in the presence of *C. scindens*. Monocultures of *C. difficile* DSM and MS001 strain, and cocultures of DSM+CS and MS001+CS were grown in the mixed carbohydrates media for ∼7 h until they reached exponential phase. Aliquots were taken for DNA extraction for next-generation sequencing to determine the cocultures’ composition, and aliquots were taken for RNA extraction for RNA-Seq. Transcriptomes of *C. difficile* in cocultures (DSM+CS and MS001+CS) were compared to the *C. difficile* monocultures’ transcriptome. The panel on the right shows the stacked bar plot of the composition of the samples subjected to RNA-Seq as determined by 16S sequencing. **c-d**, Volcano plots of log-transformed transcriptional fold changes for *C. difficile* DSM27147 (**c**) and MS001 strain (**d**) in the presence of *C. scindens*. Vertical dashed lines indicate 2-fold change, and the horizontal dashed line indicates the statistical significance threshold (p = 0.05). Blue indicates up-regulated genes and red indicates down-regulated genes. **e**, Bar plot of the log-transformed fold changes of the proline reductase (*prd*) genes of the DSM strain in the presence of CS compared to monoculture (blue) and the MS001 strain in the presence of CS compared to monoculture (pink). Asterisks above the bars indicate the adjusted *p*-value from DESeq2 differential gene expression analysis: * indicates p<0.05, *** indicates p<0.001, ns indicates not significant (p>0.05). **f**, Sensitivity of *C. difficile* DSM27147, MS001, and *C. scindens* monoculture growth towards proline concentration in the mixed carbohydrates media. AUC48h was calculated from the growth curves in **Fig. S17a**. Data were shown as mean and 95% c.i. (shading), n = 3 biological replicates. Asterisks indicate the *p*-value from unpaired *t*-test of the AUC48h between DSM and MS001 strain at specific proline concentration: * indicates p<0.05, ** indicates p<0.01, *** indicates p<0.001. **g,** Stacked bar plot of the relative abundance of *C. difficile* DSM27147 or MS001 grown with CS in media supplemented with different proline concentrations. Each bar represents the average relative abundance of each species, and the error bars represent s.d. (n=3). Asterisks above the bars indicate the *p*-value from unpaired *t*-test of the relative abundance between MS001-CS and DSM-CS coculture at a specific proline concentration: ** indicates p<0.01, *** indicates p<0.001. **h**, Percentage reduction of *C. difficile* abundance in media supplemented with different concentrations of proline compared to media without proline. Percentage reduction was calculated based on data from **Fig. S17d**. Error bars represent s.d. (n=3).

To provide insights into the transcriptional activities that mediate the observed differences in inter-species interactions, we performed genome-wide transcriptional profiling of *C. difficile* strains DSM27147 and MS001 in the presence and absence of CS (**Fig. 3b, S16a**). For both DSM and MS001 strains, ∼45% of transcripts were differentially expressed in the presence of CS than in monoculture, indicating that the presence of CS caused a global shift in the transcriptome of *C. difficile* (**Fig. 3c-d, Table S9-10**).

To identify significant changes in transcriptional activities, we performed gene set enrichment analysis (GSEA) using Kyoto Encyclopedia of Genes and Genomes (KEGG) modules. Many biological pathways such as the amino-acid transport system, pimeloyl-ACP biosynthesis, and iron complex transport system displayed similar patterns in DSM and MS001 (**Fig. S16b-e**). In addition, both *C. difficile* strains up-regulated genes for mannitol utilization, consistent with the inability of CS to utilize mannitol (**Fig. 2b**). This implies that *C. difficile* and CS display niche partitioning in co-culture, thus reducing competition for limiting substrates. In addition, both strains down-regulated the *grd* operon which is involved in glycine utilization via Stickland metabolism. Notably, only the MS001 strain up-regulated the proline reductase (*prd*) genes for Stickland metabolism via the proline pathway (∼10 to 32-fold) (**Fig. 3e**). This implies that these *C. difficile* strains display differential utilization of proline in the presence of CS.

The growth of *C. difficile* increased with supplemented proline (**Fig. 3f, S17a**). The MS001 strain displayed a significantly larger increase in growth than the DSM strain in the presence of intermediate proline concentrations. Although there are some variations in the sequence of the *prd* operon genes among *C. difficile* isolates, their protein-coding sequences are largely similar (**Fig. S17b-c**). By contrast, variation in supplemented proline did not alter the growth of CS. This demonstrates that proline metabolism via the Stickland pathway is crucial for *C. difficile* growth, but not a major resource utilized by CS in monoculture. However, we observed an opposite trend in co-cultures where increasing proline concentrations reduced *C. difficile* growth in the community (**Fig. 3g, S17d**). These results suggest that CS competed more efficiently with *C. difficile* over proline in co-culture, which was distinct from its metabolic niche in monoculture. The absolute abundance of CS increased with supplemented proline only in co-culture with the MS001 strain, but not the DSM strain (**Fig. S17d**). Consistent with the monoculture data, the MS001 strain displayed higher growth than DSM in co-culture with CS (**Fig. S17d**), and its abundance was reduced to a lower degree as a function of proline compared to the DSM strain (**Fig. 3h**). These data suggest that MS001 can compete better with CS over limited proline to perform Stickland metabolism than DSM, consistent with the higher fold change in the expression of the *prd* operon (**Fig. 3e**). These trends are consistent with the stronger inhibition of CS by MS001 compared to DSM in the inferred gLV interaction network (**Fig. 2e**).

### C. difficile toxin production in communities is not explained by growth-mediated inter-species interactions

A myriad of environmental factors including specific nutrients ^62–66^, pH ^67^, and environmental stressors including alteration of the redox potential, antibiotic exposure, and temperature increase ^68^ shape the production of toxins in *C. difficile*. By modifying the environment, certain bacterial species may impact the toxin production of *C. difficile* ^69,70^. However, we lack an understanding of how toxin production is shaped by diverse human gut species. To investigate this question, we characterized *C. difficile* toxin expression in the presence of 25 individual diverse human gut species. Many of these species are prevalent and abundant in the human gut microbiome and are linked to human health and disease ^44^ (**Fig. 4a**, **S18a-b**). Individual species were co-cultured with distinct *C. difficile* strains that we previously used to study community-level interactions (DSM27147, MS008, and MS014), as well as two other *C. difficile* strains isolated from healthy individuals (Strain 292 and Strain 296) which are clustered differently from the previous strains in terms of genotype and phenotype (**Fig. 1b-d**). We measured OD_600_ and performed 16S sequencing to determine species absolute abundances, and end-point toxin quantification using ELISA (**Fig. 4b**). A gLV model was fit to the time-resolved absolute abundance data (0, 12, 24 h) to infer inter-species interactions (**Fig. S18c-d,** DATASET004 in **Table S8**). The inferred interaction parameters using this dataset displayed an informative relationship with the parameters inferred in **Fig. 2e** (DATASET003) (**Fig. S18e**).

**Figure 4.**
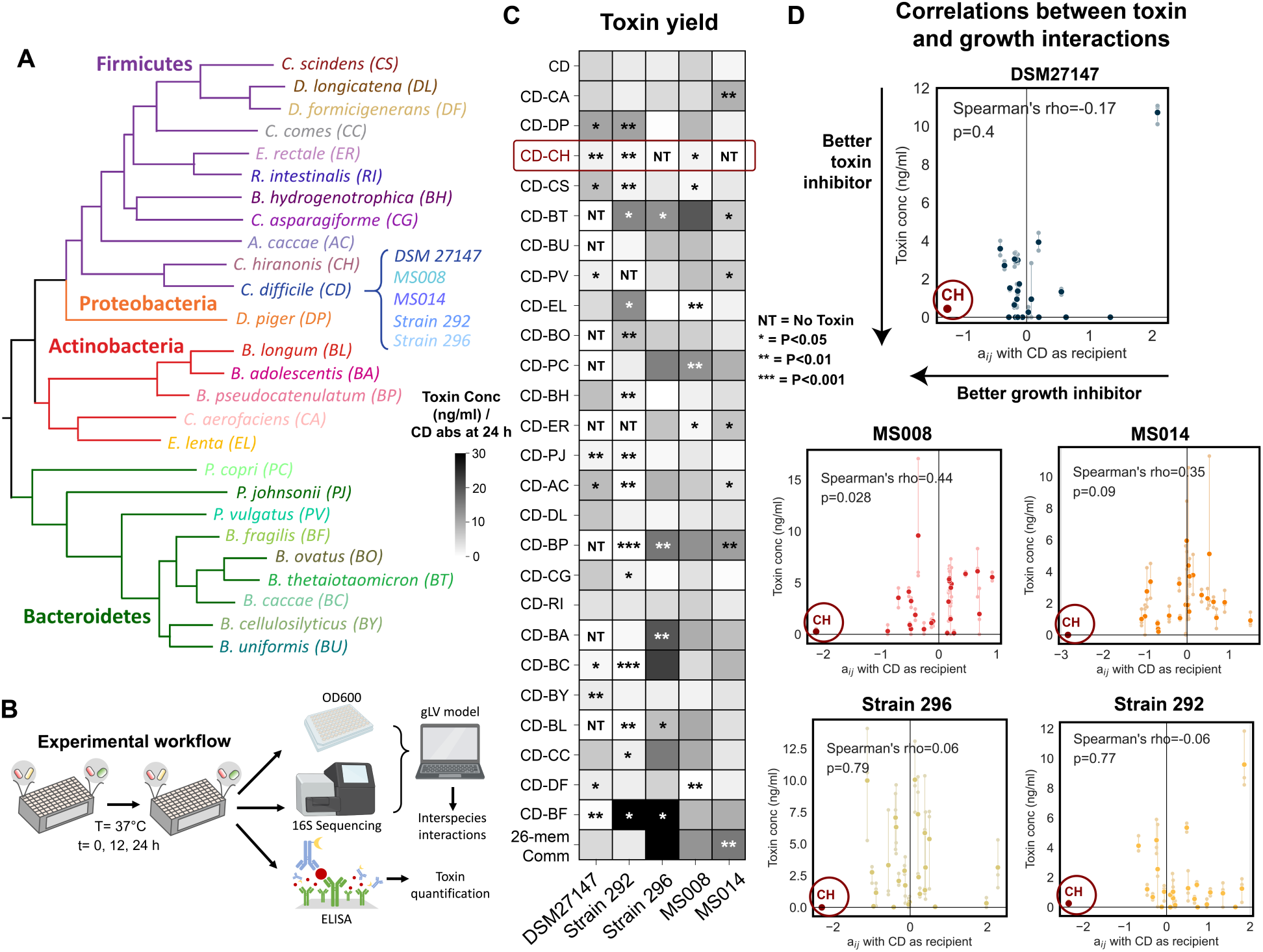
*C. hiranonis* inhibits the growth and toxin production of diverse *C. difficile* strains. **a**, Phylogenetic tree of 25-member resident synthetic gut community and *C. difficile*. **b,** Schematic of the experimental workflow. *C. difficile* was grown with gut communities and samples were taken at 12 and 24 h. Samples were subjected to OD_600_ measurement and 16S sequencing to determine species absolute abundance. Time series abundance measurements were fitted to the gLV model to obtain the interaction parameters of the community members. Samples at 24 h were subjected to toxin quantification via ELISA. **c**, Heatmap of toxin yield (toxin production per *C. difficile* abundance at 24 h) of different *C. difficile* strains when grown in pairwise and 26-member communities with human gut bacteria in the mixed carbohydrates media. Toxin concentrations (TcdA and TcdB) were measured in monocultures or communities after 24 h of growth using ELISA (n=3) (**Fig. S19a**). Asterisks on the heatmap indicate the *p*-value from unpaired *t*-test of the toxin yield in cocultures compared to *C. difficile* monocultures: * indicates p<0.05, ** indicates p<0.01, *** indicates p<0.001, NT indicates No Toxin (toxin concentration per CD absolute abundance = 0 ng/ml). **d**, Scatter plots of the interspecies interaction coefficients (aij where *C. difficile* is the recipient) versus toxin production in cocultures. Solid data points indicate the mean of the biological replicates which are represented by transparent data points connected to the mean with transparent lines. The lower the toxin concentration indicates a better toxin inhibitor and the more negative the aij indicates a better *C. difficile* growth inhibitor. Spearman’s rho and *p*-value are shown, which were computed using the spearmanr from the scipy package in Python. Parts of the figure are generated using Biorender.

Toxin yield (toxin concentration normalized by the *C. difficile* absolute abundance at 24 hr) provides insight into context-dependent changes in toxin production, whereas the toxin concentration may be more physiologically relevant. In 16.2% of conditions, toxin yields were enhanced in communities than in monoculture (36.2% for toxin concentration) (**Fig. 4c**, **S19a**). Meanwhile, in 26.2% of conditions, toxin yields were reduced in communities compared to monoculture (25.4% for toxin concentration). Genotype and toxin production did not display an informative relationship since the similar hypervirulent strains DSM27147 and MS014 displayed very different toxin production profiles in communities. Overall, *C. difficile* strains exhibited substantial variability in toxin production with Strain 296, MS008, and MS014 displaying greater similarity to each other than the other strains (Spearman’s rho=0.53-0.75, P=5.4E-03 to 1.1E-05) (**Fig. 4c, S19b**). These strains displayed higher toxin production in many pairwise communities (e.g. BT, BU, PV, PC, BP, BA, BC, BL, CC, and BF) and the 26-member community. The similarities in toxin production profiles were not explained by toxin protein-coding sequences (**Fig. S19c**). While Strain 296 and MS014 clustered together based on their metabolic genes, MS008 has distinct metabolic genes (**Fig. S3a**). These imply that toxin production in communities is likely impacted by regulatory networks and other cellular processes ^71–73^ that are shaped by gut microbiota inter-species interactions.

Some stresses including nutrient limitations have been reported to induce *C. difficile* toxin production ^72,74^. Strong negative inter-species interactions may activate stress response networks leading to an increase in toxin production. However, our results revealed that toxin production and the inferred pairwise gLV interaction coefficients impacting *C. difficile* growth in communities did not display an informative relationship (**Fig. 4d**). For instance, although the abundance of *C. difficile* Strain 296 was lower than DSM, MS008, and MS014 in the 26-member community (**Fig. S18d**), this strain displayed the highest toxin expression level (**Fig. S19a, d**). In sum, *C. difficile* strain-level variability and human gut microbiota inter-species interactions beyond growth were major variables shaping toxin production.

### C. difficile metabolism, growth, and toxin production are substantially impacted by C. hiranonis

Based on the inferred inter-species interaction network, CH inhibited distinct *C. difficile* strains regardless of whether the nutrient environment favored competition or *C. difficile* growth (**Fig. 2d-e**). Of 25 diverse gut bacteria, CH is the only species that robustly inhibited both *C. difficile* growth and toxin production of diverse *C. difficile* strains (**Fig. 4c-d**), highlighting its potential as a “universal” *C. difficile* inhibitor. This robustness of inhibitory interaction across the two nutrient environments and strain background may be attributed to the substantial metabolic niche overlap for carbohydrate utilization (**Fig. 2b**) and capability for amino acid Stickland metabolism. In addition, introducing *C. hiranonis* into communities with specific human gut species enhanced *C. difficile* growth and toxin inhibition than in co-culture with only *C. hiranonis* (**Fig. S13a-b**).

To provide insights into the mechanisms by which CH inhibits *C. difficile*, we performed genome-wide transcriptional profiling of *C. difficile* DSM27147 in the presence and absence of CH (**Fig. 5a**, **S16a, f**). In the presence of CH, 36% of *C. difficile* genes were differentially expressed compared to monoculture (**Fig. 5b, Table S11**). The transcriptional profile of *C. difficile* in the presence of CH was largely different compared to the co-culture with CS (17% of genes have an opposite sign of fold change) (**Fig. 5c**).

**Figure 5.**
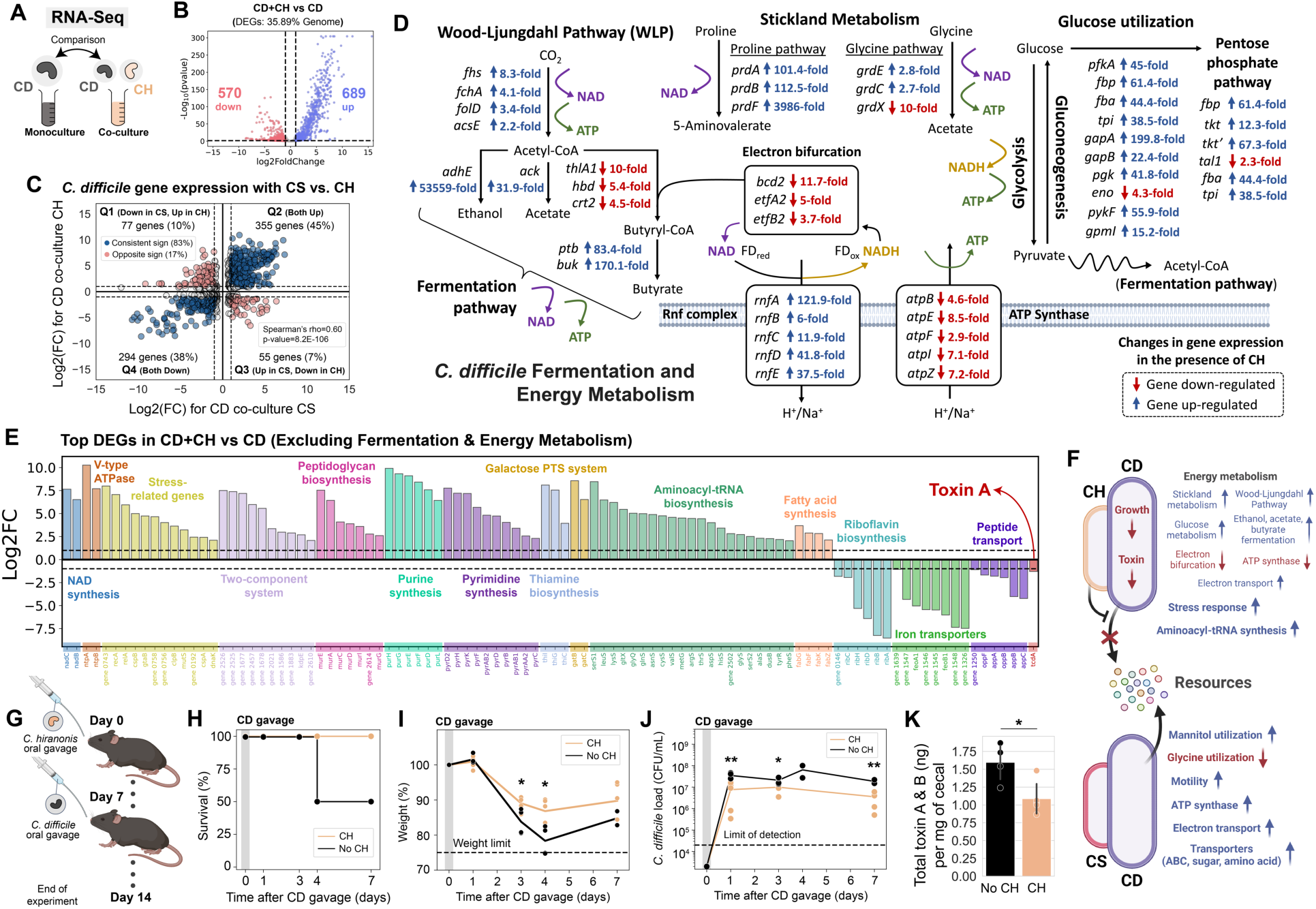
*C. hiranonis* altered *C. difficile* metabolism and other important cellular processes. **a**, Schematic of the genome-wide transcriptional profiling experiment of *C. difficile* DSM 27147 strain in the presence of *C. hiranonis*. Monocultures of *C. difficile* DSM and cocultures of DSM+CH were grown in the mixed carbohydrates media for ∼7 h until they reached the exponential phase. Aliquots were taken for DNA extraction for next-generation sequencing to determine the cocultures’ composition, and aliquots were taken for RNA extraction for RNA-Seq. Transcriptome of *C. difficile* in coculture is compared to the *C. difficile* monocultures’ transcriptome. **b**, Volcano plot of log-transformed transcriptional fold changes for *C. difficile* DSM strain in the presence of *C. hiranonis*. Vertical dashed lines indicate a 2-fold change, and the horizontal dashed line indicates the statistical significance threshold (p = 0.05). Blue indicates up-regulated genes and red indicates down-regulated genes. **c**, Scatter plot of fold changes of *C. difficile* DSM 27147 differentially expressed genes in the presence of CS and CH. Only genes with *p*-value less than 0.05 are shown. Blue indicates a consistent sign of fold changes whereas red indicates an opposite sign of fold changes. Grey indicates genes that are not differentially expressed in the presence of CS and CH (less than 2-fold change marked by the dashed lines). Spearman’s rho and *p*-value are shown, which were computed using the spearmanr from the scipy package in Python. **d**, Differentially expressed genes that are involved in *C. difficile*’s fermentation and energy metabolism. The fold changes were shown next to the gene annotations. Blue indicates up-regulated genes and red indicates down-regulated genes. **e**, Bar plot of the log-transformed fold changes of selected highly differentially expressed genes of *C. difficile* DSM in the presence of *C. hiranonis*. Horizontal dashed lines indicate a 2-fold change. **f**, Schematic highlighting substantial transcriptional changes in *C. difficile* in the presence of CH compared to CS. **g**, Schematic of the mice experiment. Mice were orally gavaged with CH for one week prior to *C. difficile* DSM27147 challenge (n=5). As a control, one group of mice without CH was challenged *C. difficile* (n=4). **h**, Percent survival of the mice after *C. difficile* gavage. **i**, Percent of initial weight after *C. difficile* gavage. Data points indicate individual mice, and the line indicates the average of all mice in the group. The horizontal dashed line indicates the weight limit of 75%. Mice with weights that dropped below the limit were sacrificed. Asterisks indicate the *p*-value from unpaired *t*-test between the weight of mice gavaged with CH and mice without CH: * indicates p<0.05. **j**, *C. difficile* abundance in the fecal and cecal content over time as determined by CFU counting on *C. difficile* selective plates. The horizontal dashed line indicates the limit of detection. Asterisks indicate the *p*-value from unpaired *t*-test between the CFU of *C. difficile* of mice gavaged with CH and mice without CH: ** indicates p<0.01, * indicates p<0.05. **k**, Total amount of *C. difficile* toxin per mg of cecal content. Data were shown as mean ± s.d. (n=5). Asterisks indicate the *p*-value from unpaired *t*-test between the toxin amount from the cecal samples of mice gavaged with CH and mice without CH: * indicates p<0.05. Parts of the figure are generated using Biorender.

Notably, co-culturing with CH yielded a massive alteration in the expression of fermentation and energy metabolism genes in *C. difficile* (**Fig. 5d**). Many genes involved in glycolysis, pentose phosphate pathway, Stickland metabolism, Wood-Ljungdahl Pathway (WLP), and fermentation pathway were highly up-regulated in the presence of CH. Since ATP synthases were down-regulated, it is possible that the cells were forced to generate ATP through the aforementioned pathways to perform essential cellular functions. *C. difficile* couples certain fermentation pathways, such as the butyrate fermentation, to the generation of a sodium/proton gradient using electron bifurcation in combination with the membrane-spanning Rnf complex ^75^. Electron bifurcation couples the NADH-dependent reduction of a substrate to the reduction of ferredoxin. The free energy resulting from the redox potential difference between ferredoxin and NAD^+^ is used to transport ions across the membrane through the Rnf complex, generating NADH in the process. Since electron bifurcating enzymes were down-regulated, Rnf complex genes were up-regulated, and glycolysis genes were highly up-regulated, *C. difficile* likely needed to generate NAD^+^ in the presence of CH. This could be achieved by the reductive Stickland metabolism or the WLP coupled to fermentation pathways. *C. difficile* heavily relies on Stickland reactions for reductive pathways ^76^. When there are abundant preferred electron acceptor substrates such as proline and glycine, the WLP is not used by *C. difficile*. However, *C. difficile* uses WLP as its terminal electron sink to support growth on glucose when *C. difficile* lacks Stickland amino acid acceptors ^76^. Therefore, the concomitant up-regulation of the proline and glycine reductases and genes involved in the WLP suggests that *C. difficile* competed with CH over proline and glycine and thus resorted to the WLP as an alternative electron-accepting pathway.

In addition to altering *C. difficile*‘s metabolism, CH impacted the expression of genes involved in various important cellular pathways such as stress responses (**Fig. 5e, S16g**). For instance, genes related to two-component systems that enable bacteria to adapt to diverse environmental changes, and many stress response genes including *recA* and *relA* were highly up-regulated. Consistent with the inhibition of *C. difficile*’s toxin production in the presence of CH as measured by ELISA, the toxin A (*tcdA*) gene was down-regulated in the presence of CH. Since *C. difficile* toxin expression is tightly linked with metabolic activity ^72^, toxin inhibition by CH could be associated with the massive changes in *C. difficile*’s metabolism. In sum, CH blocked access of *C. difficile* to alternative resource niches and led to a global alteration in the metabolic activities of *C. difficile*, providing insights into mechanisms that could mediate inhibitory inter-species interactions that are robust to strain and nutrient variability (**Fig. 5f**). In contrast, another closely related species, CS, loses its inhibitory activity in the presence of multiple carbohydrates since *C. difficile* can utilize mannitol, which is not utilized by CS.

### C. hiranonis ameliorates the effects of C. difficile in germ-free mice

To examine whether CH could inhibit *C. difficile in vivo*, we used gnotobiotic mice and orally gavaged them with CH for one week to allow time for colonization and immune development ^77^ (**Fig. 5g**). After one week, the mice were orally gavaged with the hypervirulent *C. difficile* DSM27147. As a control, we also gavaged germ-free mice with *C. difficile* (no CH group). Four days after *C. difficile* inoculation, 50% of the mice from the control group (no CH) died (**Fig. 5h**). Although mice orally gavaged with CH also exhibited a decreasing trend in weight during the first few days of *C. difficile* gavage, the relative reduction in weight was lower than the control group (5.3% and 8.5% higher after 3 and 4 days of *C. difficile* challenge respectively) (**Fig. 5i**). While CH has a low relative abundance when co-cultured with *C. difficile* DSM27147 *in vitro* (∼11% in the glucose media and ∼18% in the mixed carbohydrates media after 24 h of growth), CH was highly abundant in mice (∼72% after 7 days of *C. difficile* challenge) (**Fig. S20**). The mice harboring CH also displayed lower *C. difficile* abundance and toxin concentration than in the absence of CH (**Fig. 5j-k**). Thus, CH ameliorates the disease severity of a *C. difficile* challenge in a murine model.

## DISCUSSION

Defined communities that have been optimized to inhibit *C. difficile* hold tremendous promise to overcome the limitations of FMT for treating CDI. For instance, oral consortia from VE303 (Vedanta Biosciences) has passed the phase 2 clinical trial for rCDI ^11^ and is currently undergoing phase 3. Robustness of anti-*C. difficile* activity to environmental variability is not typically considered in the design process. This potential lack of robustness may contribute to the failure of the community to successfully treat a fraction of patients (∼14% after a few months) ^78,79^. The *C. difficile* inhibitory activity of defined communities may be more variable than fecal communities used during FMT due to their reduced functional redundancy, richness, and diversity ^7,8^. Therefore, there is a need to understand how anti-*C. difficile* activity of human gut communities varies in response to diverse *C. difficile* strain backgrounds and environmental contexts (e.g. variations in diet).

Systems biology approaches combining experiments and computational modeling have been used to understand *C. difficile* metabolism and virulence ^80^, study interactions with human gut communities ^31^, and design a bacterial consortium that protects against CDI ^81^. For instance, genome-scale metabolic models were used to guide the design of communities with enriched amino acid metabolism pathways associated with successful FMTs for rCDI treatment ^81^. However, the robustness of the designed communities to environmental context was not evaluated, and thus it is unknown whether they are effective across different strain or nutrient contexts. We used a data-driven approach to dissect interspecies interactions and toxin production of genotypically diverse *C. difficile* strains in human gut communities under different nutrient environments. We combined high-throughput *in vitro* experiments with computational modeling to deduce interaction networks impacting each *C. difficile* strain in different media conditions. We showed that *C. difficile* strain variation could directly or indirectly shape interspecies interactions of human gut microbiota. In addition, strain-level variability has a major impact on toxin production in communities, adding another layer of complexity to the design of robust anti-*C. difficile* consortia. The nutrient environment also plays a key role in shaping the interactions between *C. difficile* and the gut communities. Although it has been reported that *C. difficile* inhibition is prevalent in media that promote resource competition ^31^, we showed that it is sparse when there are multiple preferred carbohydrates for *C. difficile*. Our study showcases our quantitative systems-biology approach to map context-dependent interactions and provides insights into the mechanisms that could enhance the robustness of inhibition across strains and environments. Based on our results, interactions that lead to global shifts in metabolism and other cellular processes may exhibit greater robustness to environmental variability. More broadly, this framework considering robustness as a feature could be applied to the design of anti-pathogen bacterial therapeutics.

Of the 7 gut bacteria used to study community interactions, CS and CH are the only two species that can utilize amino acids to perform Stickland metabolism, similar to *C. difficile*. In the media supplemented with only glucose as a sole carbohydrate, CS and CH have a stronger magnitude of *C. difficile* inhibition compared to the other species (**Fig. 2d**). These inhibitory interactions may stem from competition over Stickland amino acids in addition to glucose, whereas the other gut bacteria only compete for glucose. Previous work has shown that introducing Stickland amino acid competitors can protect mice from CDI ^33^. In sum, competition over Stickland amino acids is an attractive strategy to enhance inhibition against *C. difficile*. However, in the media containing multiple carbohydrates, CH is the only species that can inhibit *C. difficile* whereas CS lost this inhibition capability (**Fig. 2e**). In a different rich media, CH inhibition of *C. difficile* was proposed to arise partially from resource competition and not via external pH change or extracellular protein release ^31^. Our results go beyond this study by demonstrating that CH suppresses the growth and toxin production of diverse *C. difficile* strains in two distinct nutrient environments, yields a massive change in the metabolic activity of *C. difficile*, and improves disease severity in germ-free mice (**Fig. 5**). Although, to our knowledge there is no evidence regarding the role of CH on CDI outcomes in humans, the presence of CH is negatively associated with *C. difficile* colonization in dogs and cats ^82–84^.

A key question is how CH maintains its inhibitory effect on *C. difficile* when provided with multiple *C. difficile*-preferred carbohydrates, whereas the inhibitory capability is abolished for CS. Since CH and CS are closely related, we would expect a similar transcriptional response in *C. difficile* in the presence of these two species. Genome-wide transcriptional profiling revealed that *C. difficile* exhibited a substantial difference in gene expression in the presence of CH and CS (**Fig. 5c**). These data provided insights into the unique transcriptional signature of CH’s inhibition mechanism, which was not observed in the presence of CS. Although our results support the hypothesis that *C. difficile* competes for Stickland amino acids with CS, *C. difficile* could switch to mannitol as an alternative nutrient source, which cannot be utilized by CS (**Fig. 5f**). By contrast, CH and *C. difficile* share highly similar metabolic niches, which may substantially limit the available resources for *C. difficile*. Therefore, *C. difficile* increased expression of enzymes in core energy-generating metabolic pathways in the presence of CH, including glycolysis, pentose phosphate pathway, Stickland metabolism, Wood-Ljungdahl Pathway (WLP), and fermentation (acetate, ethanol, and butyrate production) (**Fig. 5d**), which were not observed when CS was present. Because *C. difficile* normally favors Stickland fermentation over WLP as their main electron-accepting pathway, the activation of WLP suggests that CH successfully competes for reductive Stickland amino acids and forces *C. difficile* to use WLP as their alternative electron sink ^76^. These massive alterations in *C. difficile* core metabolism also impact virulence such as toxin production. Further, *C. difficile* upregulated stressed-related pathways (**Fig. 5e**), which were not observed in the presence of CS (**Fig. S16d-e**). Beyond resource competition, CH may produce an antimicrobial targeting *C. difficile* as previously hypothesized ^31^ that contributes to this unique transcriptional response. Future work could mine the biosynthetic gene clusters in CH for potential antimicrobial compounds and perform targeted and untargeted metabolomics to provide deeper insights into the mechanisms of inter-species interaction.

Certain bacteria in the gut have been reported to increase *C. difficile* toxin production and enhance their fitness and virulence *in vivo*, such as the opportunistic pathogen *Enterococcus faecalis* ^70^. Some metabolites produced by gut microbes such as butyrate could also increase *C. difficile* toxin, albeit moderately ^85^. However, we found that the enhancement of *C. difficile* toxin is sparse among human gut commensals (toxin production per unit biomass is enhanced in only ∼16% of all communities compared to monocultures). In addition, strain-level variability played a larger role in toxin production in communities than inferred gLV growth-mediated inter-species interactions. Since toxin production is tightly linked with metabolism ^73,86^, genotypic variations among *C. difficile* strains would impact their toxin production profiles. The lack of an informative relationship between growth-mediated inter-species interactions and toxin production suggests that inhibiting *C. difficile* growth may not always protect against CDI unless *C. difficile* is excluded from the community. Thus, the identification of *C. difficile* inhibitors should consider both inhibition of growth and toxin production. Further, we discovered that *C. difficile* strains with similar hypervirulent genotypes (DSM 27147 and MS014) have different toxin production profiles in communities. By contrast, an isolate from a healthy individual (Strain 296) has a similar toxin production profile with genetically distinct isolates from patients with CDI (MS008 and MS014) (**Fig. 4c, S19a-b**). This indicates that rather than the genotype of *C. difficile* alone, community context is a major variable shaping *C. difficile* toxin production.

A grand challenge for microbiome engineering is the rational design of microbial communities as living therapeutics for treating multiple human diseases involving alterations in the human gut microbiome. For CDI, a potential driver of the efficacy of FMT is the high richness and diversity of species in the fecal samples, which could repopulate the gut flora and restore colonization resistance. This is further supported by the fact that most of the products with successful outcomes in clinical trials so far are communities derived from stool samples, thus having high species richness ^87^. However, due to heavy reliance on donors, these stool-derived communities suffer from batch-to-batch variations and are designed without any knowledge of molecular mechanisms of *C. difficile* inhibition. This could be overcome by using defined communities that are standardized and optimized to inhibit *C. difficile.* However, the number of strains in a bacterial therapeutic currently scales with manufacturing cost. Our study shows that in the media with multiple carbohydrates preferred by *C. difficile* mimicking a perturbed gut condition, species richness is no longer a strong determinant of *C. difficile* inhibition, but rather the identity of the species in the community (**Fig. 2g**). Therefore, it is conceivable that small bacterial communities with high anti-*C. difficile* activity that is robust to environmental variability could be identified. We identified CH as a “universal” *C. difficile* growth and toxin inhibitor of genotypically diverse *C. difficile* strains and nutrient environments. Therefore, CH may represent a unique class of species that could be used to build a robust anti-*C. difficile* bacterial therapeutics to environmental variability. Future work will elucidate how to expand the number of species communities containing CH to further enhance the anti-*C. difficile* activity and robustness to environmental variability in the mammalian gut.

## METHODS

### Strain, media, and growth conditions

The strains used in this work were obtained from the sources listed in **Table S1**. There are a total of 18 *C. difficile* isolates. Nine of them were obtained from diseased patients who were diagnosed and treated for *C. difficile* infection (CDI) in the UW-Madison Hospital ^88^. These isolates were subjected to *C. difficile* nucleic acid amplification test (NAAT) (GeneXpert) via admission stool sample, and bacterial identification was confirmed via sequencing of the 16S rRNA gene. The other nine isolates were obtained from healthy individuals from the Winning the War on Antibiotic Resistance (WARRIOR) project ^89^. Briefly, the WARRIOR project collects biological specimens, including nasal, oral, and skin swabs and saliva and stool samples, along with extensive data on diet and MDRO risk factors, as an ancillary study of the Survey of the Health of Wisconsin (SHOW)^90^. WARRIOR participants include 600 randomly selected Wisconsin residents aged 18 and over, and *C. difficile* isolates were identified by anaerobic inoculation of stool samples in prereduced *C. difficile* Brucella Broth and then plated on Brucella agar plates. Colonies with correct morphology were identified using Gram staining and catalase testing. The presence of toxin genes is assessed using an in-house PCR assay and bacterial identification is confirmed via sequencing of the 16S rRNA gene.

Single-use glycerol stocks were prepared as described previously ^44^. The media used in this work are anaerobic basal broth (ABB, Oxoid) for growing starter cultures, and in-house defined media (DM) for all of the experiments. DM29 is the defined media without any carbohydrate source (recipe listed in **Table S2**), which was formulated to support the growth of phylogenetically diverse human gut bacteria ^44^ and has been used to study inter-species interactions of human gut communities ^91,92^. For supplementation of single carbohydrate sources to DM29, the carbohydrates were added to a final concentration of 5 g/L. For *mixed carbohydrates media* that mimics a perturbed gut condition, we modified DM29 by adding carbohydrate sources that are preferred by *C. difficile* and could increase in abundance upon antibiotic treatment ^45,48,58–61^, which are glucose, sorbitol, mannitol, trehalose, succinate, galactose, GalNAc, GlcNAc, and sialic acid at a concentration of 2 g/L each.

For all experiments, cells were cultured in an anaerobic chamber (Coy Lab products) with an atmosphere of 2.0 ± 0.5% H2, 15 ± 1% CO2, and balance N2 at 37 °C. Starter cultures were inoculated by adding 200 μL of a single-use 25% glycerol stock to 5 mL of anaerobic basal broth media (ABB) and grown at 37 °C without shaking.

### Growth characterization in media with different carbohydrate sources

Starter cultures of *C. difficile* isolates and gut commensal bacteria were prepared. The cell pellets from starter cultures were collected by centrifugation at 3,000 x g for 10 min, and then washed with DM29 media. The washed cell pellets were resuspended into DM29 media to a final OD_600_ of approximately 0.1. These cultures were inoculated into a 96-well plate (Greiner Bio-One) containing DM29 supplemented with specific carbohydrate sources at a concentration of 5 g/L to an initial OD_600_ of 0.01 (3 biological replicates for each strain). These plates were covered with a gas-permeable seal (Breathe-Easy^®^ sealing membrane) and incubated at 37 °C anaerobically. Cell growth determined by OD_600_ was monitored using Tecan F200 plate reader every 3 h using robotic manipulator arm (RoMa) integrated with our Tecan Freedom Evo 100 instrument.

### Logistic growth model

The logistic growth model was used to describe *C. difficile* population growth dynamics in monoculture experiments. The logistic growth model for species *i* takes the following form:

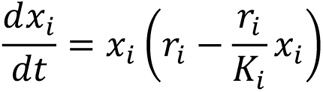

where *x_i_* is the absolute abundance of species *i*, parameter *r_i_* is its maximum growth rate, and *K_i_* is its carrying capacity. Due to the unique growth profile of *C. difficile* isolates, we cut time points where the OD_600_ drops below > 10% to exclude the highly variable phase. Thus, the steady-state solution of the model is the carrying capacity (*K_i_*) (i.e. the value of *x_i_* when 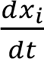 equals 0). We also excluded data points less than 120% of the initial OD_600_ (OD_600_ at t=0) to exclude the lag phase which is not captured in the logistic model. A custom MATLAB script is used to estimate the parameters θ*_i_* = [*r_i_*, *K_i_*] in the logistic growth model. For each species *i*, the model is fitted to experimental data with L2 regularization. Specifically, given a series of *m* experimental OD_600_ measurements, ***x_i_*** = [*x_i_*_1_ ..., *x_i,m_*], and a series of OD_600_ simulated using parameter θ_*i*_ at the same time intervals, 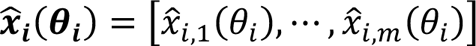, the optimization scheme minimizes the cost function:

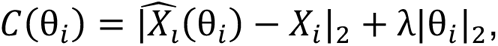

where λ is the L2 regularization parameter, which was set to be 0.02 for all species, and | ⋅ |_2_ indicates vector 2-norm. Solutions to the logistic growth model were obtained using the ode15s solver and the optimization problem was solved using FMINCON in MATLAB (R2022a).

### Fluorescence microscopy of *C. difficile*

Starter cultures of several *C. difficile* strains were prepared. The cell pellets from starter cultures were collected by centrifugation at 3,000 x g for 10 min, and then washed with DM29 media. The washed cell pellets were resuspended into DM29 media to a final OD_600_ of approximately 0.1. These cultures were inoculated into new culture tubes containing either DM29 media or DM29 supplemented with 5 g/L glucose to an initial OD_600_ of 0.01 by adding 500 μl of washed starter cultures to 4.5 mL media. After 16 h and 40 h of growth, 100 μl aliquots were taken, stained with SYBR Green dye, and viewed with a microscope (Nikon Eclipse Ti-E inverted microscope) at 20× dry objective with appropriate filter sets. Images were captured with Photometrics CoolSNAP Dyno CCD camera and associated software (NIS-Elements Ver. 4.51.00).

### Whole genome sequencing of *C. difficile* isolates

*C. difficile* DSM 27147 and the 18 isolates used in this study were subjected to whole-genome sequencing. Strains were grown from a single colony to OD_600_ of 0.3, and then centrifuged to obtain the cell pellets. Genomic DNA was extracted using Qiagen DNeasy Blood and Tissue Kit according to the manufacturer’s protocol. The harvested DNA was detected by the agarose gel electrophoresis and quantified by a Qubit fluorometer. The genomic DNA was sent to SeqCenter (Pittsburgh, PA, USA) for paired-ends Illumina sequencing. Sample libraries were prepared using the Illumina DNA Prep kit and IDT 10 bp UDI indices, and sequenced on an Illumina NextSeq 2000, producing 2 x 151 bp reads. Demultiplexing, quality control, and adapter trimming were performed with bcl-convert (v3.9.3) Illumina software. The clean bases of each sample are ∼1 billion bp. The WGS raw data was submitted and is accessible in BioProject PRJNA902807.

### Whole-genome sequencing data analysis

SPAdes Genome Assembler ^93^ is used to assemble contigs from the whole-genome sequencing data with the following parameters: spades.py *--pe1-1* (forward read fastq file) *--pe1-2* (reverse read fastq file) *--isolate -o* (output name). The *--isolate* option was used due to the high-coverage sequencing data. We then used FastANI ^94^ to compute the whole-genome Average Nucleotide Identity (ANI) values between pairs of isolates, which is defined as the mean nucleotide identity of orthologous gene pairs shared between two microbial genomes. The following parameters are used: fastANI *--ql* (list of contigs.fasta files of all isolates from SPAdes) *--rl* (list of contigs.fasta files of all isolates from SPAdes) –matrix *-o* (output name). The newly sequenced genomes are high-quality drafts with a low number of contigs (median 61 [range 40–458]) and high N50 (median 285,062 [range 146,596–782,135]) (**Table S3**).

Annotation of the contigs was performed using DFAST ^95^. For further comparative genomic analyses, the gene content across 19 C*. difficile* strains was analyzed by clustering all predicted coding sequences into orthologous groups ^96^ (**Fig. S3b**). Clustering of gene orthologs was carried out using ProteinOrtho6 ^96^ across variable coverage and identity settings using BlastP for alignment. Distributions of OGs show a high degree of strain variability with many genes in a limited subset of strains (**Fig. S3c**).

To get Gene Ontology (GO) information, we used BlastP of the NCBI Blast Suite^97^ against the proteins from all *C. difficile* strains that exist in UniProt database (downloaded from UniProtKB on 10^th^ November 2022) at 1E-3 E-value cutoff. Following BlastP, GO information such as biological process, molecular function, and cellular compartment of each protein was extracted from UniProt. To align the sequence of specific genes such as Toxin A (*tcdA*), Toxin B (*tcdB*), RNA polymerase (*rpoB, rpoB’*), we used Clustal Omega multiple sequence alignment tool ^98^.

We evaluated the genetic diversity of our *C. difficile* strains in the context of the other 118 publicly available *C. difficile* genomes (**Table S4**). Phylogenomic analysis was performed using GToTree ^99^ on the *C. difficile* isolates dataset along with 118 public strains. SPAdes FASTA files were used as inputs to GToTree analysis and the resulting tree was visualized using the Interactive Tree of Life web-based tool ^100^. Our isolates span 64% of the *C. difficile* phylogeny of this dataset (9 of 14 major tree branches) (**Fig. S3d**).

To get the relative copy number of the genes in each isolate, the Illumina paired- end reads were aligned to the gene list from DFAST using Bowtie2 ^101^. The detection of conjugative systems was performed using CONJScan ^102^ module of MacSyFinder. The detection of phages was performed using VirSorter ^103^.

### Construction of strain-specific genome-scale metabolic models to assess variations in metabolism

Raw sequencing data was first preprocessed using fastp 0.22.0 ^104^, trimming the first 5bp at the 5’ end and trimming the 3’ end with a sliding window approach, maintaining a minimum quality score of 20. Reads shorter than 60bps were omitted. 85%-95% of reads passed all filters across samples, yielding 2.9M to 7.1M reads per sample. Preprocessed reads were assembled using MEGAHIT 1.2.9 ^105^ using default k-mer sizes and a minimum contig length of 1000bps. Completeness and contamination were assessed using CheckM2 1.0.1 ^106^ yielding completeness of >99.9% for all assemblies while maintaining contamination below 1.5%. Bacterial species identity was verified using the GTDB toolkit 2.1.0 ^107^ using the database version 207. All assemblies were classified as *Clostridioides difficile* by average nucleotide identity and placement in the GTDB reference tree.

*De novo* gene predictions of the assemblies were performed by Prodigal 2.6.3 ^108^. Metabolic draft models were built using CarveMe 1.5.2 ^109^ from the isolate gene predictions using DIAMOND 2.1.6 ^110^ with additional options of “--more-sensitive –top 10 ”. Media composition was translated by manual mapping to the BiGG database ^111^. Salts were decomposed into their aqueous phase ions to mimic the effect of hydrolysis in the translated medium. Draft models were then gapfilled to be able to grow on the mapped media. During gapfilling, no more than 10 new reactions and 6 new metabolites were added to each model. Model quality was assessed using MEMOTE 0.13.0 ^112^. Metabolic reaction content was assessed using the “metabolic_dist” function from MICOM 0.32.5^113^ where metabolic distances were calculated by the Jaccard distance of metabolic reaction absence/presence (1 - shared reactions / total reactions) for each pair of reconstructed models.

### Bacterial genome DNA extraction for 16S amplicon sequencing

All the genomic DNA (gDNA) extraction and next-generation sequencing sample preparation were performed as described previously ^31,44^. Bacterial gDNA extractions were carried out using a modified version of the Qiagen DNeasy Blood and Tissue Kit protocol in 96-well plates. Briefly, cell pellets were resuspended in 180-μl enzymatic lysis buffer containing 20 mg/ml lysozyme (Sigma-Aldrich), 20 mM Tris–HCl pH 8 (Invitrogen), 2 mM EDTA (Sigma-Aldrich), and 1.2% Triton X-100 (Sigma-Aldrich), and then incubated at 37°C at 600 RPM for 30 min. Samples were treated with 25 μL 20 mg/ml Proteinase K (VWR) and 200 μL buffer AL (Qiagen), mixed by pipette, and then incubated at 56°C at 600 RPM for 30 min. Samples were treated with 200 μL 200 proof ethanol (Koptec), mixed by pipette, and transferred to 96-well nucleic acid binding plates (Pall). After washing with 500 μL buffer AW1 and AW2 (Qiagen), a vacuum was applied for 10 min to dry excess ethanol. Genomic DNA was eluted with 110 μL buffer AE (Qiagen) preheated to 56°C and then stored at −20°C.

Genomic DNA concentrations were measured using the Quant-iT™ dsDNA Assay Kit (Invitrogen) with a 6-point DNA standard curve (0, 0.5, 1, 2, 4, 6 ng/μL biotium). 1 μL of samples and 5 μL of standards were diluted into 95 μL of 1× SYBR green (Invitrogen) in TE buffer and mixed by pipette. Fluorescence was measured with an excitation/emission of 485/535 nm (Tecan Spark). Genomic DNA was then normalized to 2 ng/µL by diluting in molecular grade water (VWR International) using a Tecan Evo Liquid Handling Robot.

Dual-indexed primers for multiplexed amplicon sequencing of the V3-V4 region of the 16S rRNA gene were designed as described previously ^38,44^. PCR was performed using the normalized gDNA as template and Phusion High-Fidelity DNA Polymerase (Thermo Fisher) for 25 cycles with 0.05 μM of each primer. Samples were pooled by plate, purified using the DNA Clean & Concentrator™-5 kit (Zymo) and eluted in water, quantified by NanoDrop, and combined in equal proportions into a library. The library was quantified using Qubit 1× HS Assay (Invitrogen), diluted to 4.2 nM, and loaded at 10 pM onto Illumina MiSeq platform for 300-bp paired-end sequencing using MiSeq Reagent Kit v2 (500-cycle), or loaded at 21 pM using MiSeq Reagent Kit v3 (600-cycle) depending on the desired sequencing reads.

### 16S amplicon sequencing data analysis to determine community composition

Sequencing data were analyzed as described previously ^31,38^. Briefly, reads were demultiplexed with Basespace FastQ Generation, and the FastQ files were analyzed using custom Python scripts. Paired reads were merged using PEAR (Paired-End reAd mergeR) v0.9.0 ^114^. A reference database containing 16S V3-V4 region of each species in the study was created by assembling consensus sequence based on sequencing results of each monospecies. Reads were mapped to the reference database using the mothur v1.40.5 command classify.seqs using the Wang method with bootstrap cutoff value of 60% ^115,116^. Relative abundance was calculated by dividing the read counts mapped to each organism by the total reads in the sample. Absolute abundance was calculated by multiplying the relative abundance of an organism by the OD_600_ of the sample. Samples were excluded from further analysis if > 1% of the reads were assigned to a species not expected to be in the community (indicating contamination).

### Parameter estimation of generalized Lotka-Volterra models

The generalized Lotka-Volterra (gLV) model is a set of coupled ordinary differential equations that describe the growth of interacting species over time,

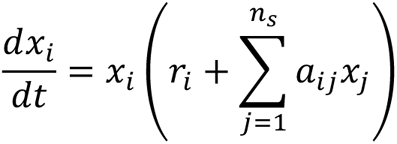

where *x_i_* is the abundance of species *i* and *n_s_* is the total number of species. Model parameters that need to be estimated from data include the species growth rate, denoted as *r_i_*, and coefficients that determine how species *j* affects the growth of species *i*, denoted as *a_ij_*. The data used for parameter estimation is the growth of species over time under different inoculation conditions. For monoculture growth data, we use OD_600_ measurements only, whereas for community data, this was obtained by multiplying the relative abundance obtained from 16S sequencing by the total OD_600_.

A prior over the parameter distribution is set so that growth rates have a mean of 0.3, self-interaction terms have a mean of -1, and inter-species interaction terms have a mean of -0.1. Given a dataset of measured species abundances over time after inoculating different combinations of species, the model parameters are determined by minimizing a cost function given by a weighted squared difference between model-predicted species abundances and measured abundances and a penalty for deviations from the prior mean. Using the fitted parameter estimates, the covariance of the posterior parameter distribution is approximated as the inverse of the Hessian (matrix of second derivatives) of the cost function with respect to the model parameters. The Expectation-Maximization (EM) algorithm is used to optimize the precision of the prior parameter distribution and the precision of the noise distribution, which collectively determine the degree to which estimated parameters are penalized for deviations from the prior mean^117^. In other words, the precision of the prior and noise are hyperparameters that determine the degree of regularization. To evaluate model prediction performance on held-out data, we performed 10-fold cross validation where the degree of regularization was optimized using the EM algorithm and only community samples were subjected to testing (i.e. monoculture data was reserved only for model training). See **Supplementary Text** for a more detailed description of parameter estimation and the EM algorithm.

### Bayesian experimental design to guide community experiments

We define an experimental design as a set of unique inoculation conditions, where in each condition each species may be present or absent, and the total inoculation density sums to OD_600_ of 0.01. We used a Bayesian experimental design approach to select experimental conditions that were expected to collectively minimize parameter uncertainty as quantified by the expected Kullback-Leibler (KL) divergence between the posterior parameter distribution and the prior parameter distribution (See Equation 20 in **Supplementary Text**).

### Growth of synthetic gut communities with *C. difficile* isolates

Starter cultures of all *C. difficile* isolates and commensal gut bacteria were prepared. The cell pellets from starter cultures were collected by centrifugation at 3,000 x g for 10 min, and then washed with DM29 media. The washed cell pellets were resuspended into DM29 media to a final OD_600_ of approximately 0.1.

For the growth experiment of each of the 19 *C. difficile* strains with 8-member gut bacteria at a single timepoint (**Fig. S6b-c**), the monocultures of individual *C. difficile* strains and each gut species were mixed in equal proportions based on OD_600_ and inoculated into 2 mL 96-deep-well plates (Nest Scientific) containing DM29 supplemented with specific carbohydrate sources (glucose, mannitol, galactose, or mucin) to an initial OD_600_ of 0.01. The initial OD_600_ of each species is therefore 0.0011 (0.01 divided by 9, the number of species in the community). As a control, we also inoculated a mixture of gut species without *C. difficile* to the same initial OD_600_ of 0.01. There are a total of 4 plates for media with different carbohydrate sources, each containing 20 communities (18 for different *C. difficile* isolates, 1 for *C. difficile* DSM 27147 strain, and 1 for the gut community without *C. difficile*), with 3 biological replicates for each community. These plates were covered with a gas-permeable seal (Breathe-Easy^®^ sealing membrane) and incubated at 37 °C anaerobically for 24 hours to capture *C. difficile* growth prior to the highly variable late stationary phase response. At the end of the experiment, OD_600_ was measured with a Tecan F200, and cell pellets were collected for DNA extraction, PCR amplification, and NGS sequencing.

For the growth experiment of each of the 19 *C. difficile* strains with 7-member gut bacteria at a single time point in the mixed carbohydrates media (**Fig. S9e**), the monocultures of individual *C. difficile* strains and each gut species were mixed in equal proportions based on OD_600_ and inoculated into 2 mL 96-deep-well plate (Nest Scientific) containing the mixed carbohydrates media to an initial OD_600_ of 0.01. The initial OD_600_ of each species is therefore 0.00125 (0.01 divided by 8, the number of species in the community). As a control, we also inoculated a mixture of gut species without *C. difficile* to the same initial OD_600_ of 0.01. The deep-well plate was covered with a gas-permeable seal (Breathe-Easy^®^ sealing membrane) and incubated at 37 °C anaerobically for 24 hours. At the end of the experiment, OD_600_ was measured with a Tecan F200, and cell pellets were collected for DNA extraction, PCR amplification, and NGS sequencing.

For time-course growth experiment of 4 different *C. difficile* strains with 7 gut bacteria in the glucose media (**Fig. 2d**) or mixed carbohydrates media (**Fig. 2e**), *C. difficile* and gut bacteria were mixed and grown in 2-8 member communities. The community combinations were generated from the Bayesian experimental design (**Table S8**). The monocultures of *C. difficile* strains and each gut species were mixed in equal proportions based on OD_600_ and inoculated into 2 mL 96-deep-well plates (Nest Scientific) containing the glucose media (**Fig. 2d**), or the mixed carbohydrates media (**Fig. 2e**) to an initial OD_600_ of 0.01. For instance, the initial OD_600_ of each species in a 2-member community is therefore 0.005 (0.01 divided by 2, the number of species in the community). These plates were covered with a gas-permeable seal (Breathe-Easy^®^ sealing membrane) and incubated at 37 °C anaerobically. After 12 and 24 hours of growth, OD_600_ was measured with a Tecan F200, and cell pellets were collected for DNA extraction, PCR amplification, and NGS sequencing. For longer-term growth experiments in **Fig. S9c-d**, the communities were grown for 72 hours and passaged using a 1:20 dilution at 24 and 48 h to observe community assembly over three batch culture growth cycles and capture the longer-term behavior of the consortia.

For time-course growth experiment of 5 different *C. difficile* strains with 25 gut bacteria in the mixed carbohydrates media (**Fig. 4**), individual *C. difficile* strain and gut bacteria were mixed and grown in pairwise and full 26-member communities. The monocultures of *C. difficile* strains and each gut species were mixed in equal proportions based on OD_600_ and inoculated into 2 mL 96-deep-well plates (Nest Scientific) containing the mixed carbohydrates media to an initial OD_600_ of 0.01. For pairwise communities, the initial OD_600_ of each species is 0.005 (0.01 divided by 2), and for 26-member communities, the initial OD_600_ of each species is 0.000385 (0.01 divided by 26). These plates were covered with a gas-permeable seal (Breathe-Easy^®^ sealing membrane) and incubated at 37 °C anaerobically. After 12 and 24 hours of growth, OD_600_ was measured with a Tecan F200, and cell pellets were collected for DNA extraction, PCR amplification, and NGS sequencing. Supernatants of communities at 24 hours of growth were collected for toxin quantification using ELISA.

### *C. difficile* toxin measurements using ELISA

Toxin (both TcdA and TcdB) concentrations in *C. difficile* monocultures or co-cultures, and toxin titer in mice cecal samples were determined by comparison to a standard curve using ELISA (tgcBiomics, Germany). The blank media used to grow the cultures were also included in the assay to measure any background noise. Samples subjected to toxin measurements in this study were processed in parallel at the same time using the same batch of ELISA kits to minimize batch-to-batch variations and ensure comparable results.

### Growth of *C. aerofaciens* and *B*. *thetaiotaomicron* in the sterilized spent media of different *C. difficile* strains

Starter cultures of *C. difficile* DSM27147, MS001, MS008, and MS014 were prepared. The cell pellets from starter cultures were collected by centrifugation at 3,000 x g for 10 min, and then washed with DM29 media. The washed cell pellets were resuspended into DM29 media to a final OD_600_ of approximately 0.1. Each of the *C. difficile* strains was inoculated into new culture tubes containing DM29 media supplemented with 5g/L glucose to an initial OD_600_ of 0.01. Culture tubes were incubated at 37°C with no shaking. After an incubation time of 24 h, cultures were spun down aerobically at 3,000 x g for 20 min and sterile filtered using Steriflip 0.2-μM filters (Millipore-Sigma) before returning to the anaerobic chamber.

Then, starter cultures of *C. aerofaciens* and *B. thetaiotaomicron* were prepared. The cell pellets from starter cultures were collected by centrifugation at 3,000 x g for 10 min, and then washed with DM29 media. The washed cell pellets were resuspended into DM29 media to a final OD_600_ of approximately 0.1. CA-BT coculture was inoculated in the sterilized spent media of each *C. difficile* strain mixed with fresh media (DM29 supplemented with 5g/L glucose) at an equal ratio to replenish the nutrients. CA and BT were inoculated at an equal initial abundance to a final OD_600_ of 0.01 in 2 mL 96-deep-well plates (Nest Scientific) that were covered with gas-permeable seals (BreatheEasy), and incubated at 37°C with shaking. After 24 h, OD_600_ of the cultures were measured and the cell pellets were collected for DNA extraction, PCR amplification, and NGS sequencing.

### Growth of *C. difficile* strains in the sterilized spent media of gut bacteria

Starter cultures of commensal gut bacteria were prepared. The cell pellets from starter cultures were collected by centrifugation at 3,000 x g for 10 min, and then washed with DM29 media. The washed cell pellets were resuspended into DM29 media to a final OD_600_ of approximately 0.1. Each of the gut bacteria was inoculated into new culture tubes containing the mixed carbohydrates media to an initial OD_600_ of 0.01. Culture tubes were incubated at 37°C with no shaking. After an incubation time of 24 h, cultures were spun down at 3,000 x g for 20 min and sterile-filtered using Steriflip 0.2-μM filters (Millipore-Sigma). Media control (mixed carbohydrates media) was spun down and filtered in parallel with samples. The pH of the sterilized spent media was adjusted to the same value as the media control.

Then, starter cultures of *C. difficile* strains were prepared. The cell pellets from starter cultures were collected by centrifugation at 3,000 x g for 10 min, and then washed with DM29 media. The washed cell pellets were resuspended into DM29 media to a final OD_600_ of approximately 0.1. The *C. difficile* strains were inoculated in the sterilized spent media of each gut bacteria (and the mixed carbohydrates media as a control) to a final OD_600_ of 0.01 in 96-well microplates that were covered with gas-permeable seals (BreatheEasy). The plates were incubated at 37°C with shaking, and OD_600_ was measured every 3 h (Tecan Infinite Pro F200).

### Transcriptome profiling

*C. difficile* DSM27147 monoculture, *C. difficile* MS001 monoculture, CD DSM-CS coculture, CD MS001-CS coculture, and CD DSM-CH coculture conditions were inoculated from starter cultures into individual culture tubes containing the mixed carbohydrates media. For monoculture conditions, *C. difficile* was inoculated to an OD_600_ of 0.01. For cocultures, *C. difficile* and CS or CH were inoculated to an equal ratio (OD_600_ of 0.005 each). The cultures were incubated anaerobically at 37°C with no shaking for ∼7 h until the culture reached the exponential phase (OD_600_ ∼0.2). 1000 μL of the culture was taken for OD_600_ measurement and total DNA extraction for next-generation sequencing, and 2000 μL of the culture was taken for total RNA extraction for transcriptomics. 4000 μL of RNAprotect (Qiagen) was added to 2000 μL of culture and incubated for 5 min at room temperature. Cultures were then centrifuged at room temperature for 10 min at 3000 g and the supernatant was carefully removed. Cell pellets were immediately subjected to RNA extraction using acidic phenol bead-beating method. Pellets were resuspended in 500 μL 2× Buffer B (200 mM sodium chloride, 20 mM ethylenediaminetetraacetic acid) and transferred to 2 mL microcentrifuge tubes containing 500 μL Phenol:Chloroform:IAA (125:24:1, pH 4.5) and 210 μL 20% sodium dodecyl sulfate and were bead-beated with acid washed beads (Sigma G1277) for 3 min. All solutions used for RNA extraction were RNAse-free. Samples were centrifuged at 4°C for 5 min at 7,200 g, and 600 μL of the upper aqueous phase was added to 60 μL 3 M sodium acetate and 660 μL cold isopropanol and chilled on ice for 5 min before freezing for 5 min at −80°C. Samples were centrifuged at 4°C for 15 min at 18,200 g, the supernatant was decanted, and the pellet was washed with cold 100% ethanol. The pellets were dried in a biosafety cabinet for 15 min and then resuspended in 100 μL RNAse-free water. Samples were purified using RNeasy Mini Kit (Qiagen) and genomic DNA was removed using RNAse-Free DNase Set (Qiagen). Two replicates of each condition were sent to Novogene Corporation Inc (Sacramaneto, CA, United States of America) for rRNA depletion, cDNA library preparation, and sequencing on Illumina NovaSeq. Data was de-multiplexed using Illumina’s bcl2fastq 2.17 software, where one mismatch was allowed for index sequence identification.

The compressed FASTQ files were quality-checked using the FastQC tool v0.12.1^118^. The BBDuk, BBSplit, and BBMap tools from BBTools suite (v38.42) ^119^ were used to trim adapters, deplete rRNA, and map the remaining mRNA reads to the reference genomes. For monoculture or cocultures containing *C. difficile* DSM27147, the reference genome was obtained from GenBank (FN545816.1). For monoculture or cocultures containing *C. difficile* MS001 isolate, the reference genome was obtained from the whole-genome sequencing data that was assembled and annotated using SPAdes Genome Assembler ^93^. The feature-Counts package v1.6.4 ^120^ from the SubRead suite was used to map reads to features on the genome and quantify raw counts for each transcript. Reads per kilobase million (RPKM) values were computed using a custom Python script to see the agreement of gene expression between biological replicates. The gene expression (represented by RPKM values) shows a good correlation between biological replicates (Pearson’s R=0.95-0.98, P<10E-05) (**Fig. S16a**). The DESeq2 Bioconductor library v4.0.3 ^121^ was used in R v4.0.4 to quantify differential gene expression using a negative binomial generalized linear models with apeglm shrinkage estimator ^122^. When calculating RPKM of *C. difficile* genes in the CD-CS and CD-CH coculture, the “reads mapped” in the denominator was the number of reads mapped to the *C. difficile* genome. Similarly, when quantifying differential gene expression for *C. difficile* genes in the CD-CS and CD-CH coculture, only reads mapped to the *C. difficile* genome were provided to DeSeq2. We define differentially expressed genes (DEGs) as those with >2-fold change and a *p*-value less than 0.05. The RNA-seq data was submitted and is accessible in BioProject PRJNA983758.

### Gene Set Enrichment Analysis (GSEA)

GSEA was performed using the GSEA method of the ClusterProfiler R package (v4.2.2)^123^. KEGG modules for *C. difficile* were used as gene sets and were supplied as a user-defined annotation with the TERM2GENE field. The analysis was run with the log2FCs calculated by DeSeq2. The p-value cutoff used was 0.05 and the minimum gene set size used was 3.

### Gnotobiotic mouse experiments

All germ-free mouse experiments were performed following protocols approved by the University of Wisconsin-Madison Animal Care and Use Committee. We used 10-week-old C57BL/6 gnotobiotic male mice (wild-type) and a regular diet (Chow diet, Purina, LabDiet 5021). All strains were grown at 37 °C anaerobically in Anaerobe Basal Broth (ABB, Oxoid) to stationary phase. *C. hiranonis* and *C. difficile* DSM27147 strain for oral gavage was diluted to ∼10,000 CFU/mL, and these cultures were transferred to Hungate tubes (Chemglass) on ice prior to oral gavage. On day 0, 0.2 mL of *C. hiranonis* culture was introduced into the mice by oral gavage inside a Biological Safety Cabinet (BSC) and the mice were housed in biocontainment cages (Allentown Inc.) for the duration of the experiment. After one week, 0.2 mL of *C. difficile* (∼2,000 CFU) was introduced into the mice by oral gavage. Mice were maintained for a total of two weeks after the first colonization with the core community (day 0). Groups of mice (4-5 mice) with the same core community and *C. difficile* were co-housed in a single cage. Mice were weighed and fecal samples were collected at specific time points after oral gavage for NGS sequencing. Cecal contents from mice that were dead or sacrificed in the middle of the experiment were collected for NGS sequencing.

### Genomic DNA extraction from fecal and cecal samples

The DNA extraction for fecal and cecal samples was performed as described previously with some modifications ^124^. Fecal samples (∼50 mg) were transferred into solvent-resistant screw-cap tubes (Sarstedt Inc) with 500 μL 0.1 mm zirconia/silica beads (BioSpec Products) and one 3.2 mm stainless steel bead (BioSpec Products). The samples were resuspended in 500 μL of Buffer A (200 mM NaCl (DOT Scientific), 20 mM EDTA (Sigma) and 200 mM Tris·HCl pH 8.0 (Research Products International)), 210 μL 20% SDS (Alfa Aesar) and 500 μL phenol/chloroform/isoamyl alcohol (Invitrogen). Cells were lysed by mechanical disruption with a bead-beater (BioSpec Products) for 3 min twice, while being placed on ice for 1 min in between to prevent overheating. Next, cells were centrifuged for 7 min at 8,000 x g at 4°C, and the supernatant was transferred to an Eppendorf tube. We added 60 μL 3M sodium acetate (Sigma) and 600 μL isopropanol (LabChem) to the supernatant and incubated on ice for 1 h. Next, samples were centrifuged for 20 min at 18,000 x g at 4°C, and the supernatant was decanted. The harvested DNA pellets were washed once with 500 μL of 100% ethanol (Koptec), and the remaining trace ethanol was removed by air drying the samples. Finally, the DNA pellets were resuspended into 300 μL of AE buffer (Qiagen). The crude DNA extracts were purified by a Zymo DNA Clean & Concentrator™-5 kit (Zymo Research) prior to PCR amplification and NGS sequencing.

### *C. difficile* colony-forming unit counting from fecal and cecal samples

*C. difficile* selective plates were prepared by autoclaving C. difficile agar (Oxoid CM0601) and adding defibrinated horse blood (Lampire 7233401, 70 mL/1L media), norfloxacin (Santa Cruz 215586, 120 μg/mL), moxalactam (Santa Cruz 250419, 320 μg/mL), and erythromycin (Santa Cruz 204742, 100 μg/mL) after the media is cooled to ∼55°C. Right after mice fecal or cecal collection, around 1μL of fresh fecal samples were taken using an inoculating loop and mixed with PBS. The samples were then serially diluted (1:10 dilution) using PBS. Four dilutions of each sample were spotted on *C. difficile* selective agar plates, with 2 technical replicates per sample. Plates were incubated at 37°C for 48 h at which point colonies were counted in the dilution spot containing between 5 and 100 colonies. The CFU/mL for each sample was calculated as the average of the 2 technical replicates times the dilution factor. The lower limit of detection for the assay was 20,000 CFU/mL.

## Data availability

Whole-genome sequence data of the *C. difficile* strains will be deposited in the NCBI database. Mapped growth media and strain-specific genome scale metabolic models in SBML format can be found at https://github.com/gibbons-lab/2023_cdiff_venturelli. Nextflow pipelines for assembly and metabolic model building can be found at https://github.com/gibbons-lab/pipelines. RNA-seq data used in this study will be deposited in the NCBI database. Raw DNA sequencing data and processed sequencing data to determine community composition will be made available via Zenodo prior to publication.

## Code availability

Codes for processing sequencing data, fitting the gLV models, and performing Bayesian experimental design will be available through Github prior to publication. Until then, we have provided the code as a supplementary file.

## Acknowledgments

We would like to thank Jun Feng, Yiyi Liu, Freeman Lan, Tyler D. Ross, Erin O. Loss, Yu-Yu Cheng, and Alex Carr for their helpful advice on this project. This research was supported by the National Institutes of Allergy and Infectious Diseases under grant number R21AI156438 and R21AI159980 for O.S.V, R35GM124774 for O.S.V. The funders had no role in study design, data collection and analysis, decision to publish or preparation of the manuscript.

## Authors contributions

J.E.S. and O.S.V. conceived the study. J.E.S. carried out the experiments. Y.Q. implemented computational modeling for the logistic growth model. J.T. implemented computational modeling for the gLV models and performed Bayesian experimental design. E.I.V. assisted in experimental data collection for the mice experiments. C.D. and S.M.G. constructed the strain-specific metabolic genome scale model. N.S. collected the *C. difficile* isolates used in this study. J.E.S. and O.S.V. analyzed data. J.E.S. and O.S.V. wrote the paper and all authors provided feedback on the manuscript. O.S.V. secured funding.

