## Supplementary Information for "Elucidating human gut microbiota interactions that robustly inhibit diverse *Clostridioides difficile* strains across different nutrient landscapes"

### GLOVE: Generalized LOtka-Volterra parameter Estimation

#### 1 Generalized Lotka-Volterra Model

The generalized Lotka-Volterra (gLV) model is a non-linear ordinary differential equation model that can be used to model the dynamics of interacting microbial species. Given a set of $n_s$ interacting species $\mathbf{x} = (x_1, ..., x_{n_s})^T \in \mathbb{R}_+^{n_s}$, the gLV model is the following set of ordinary differential equations

$$\frac{\mathrm{d}x_i}{\mathrm{d}t} = x_i \left( r_i + \sum_{j=1}^{n_s} a_{ij} x_j \right), \tag{1}$$

where $r_i \in \mathbb{R}_+$ is the growth rate of species $i$, and $a_{ij}$ quantifies the effect of species $j$ on the rate of change of species $i$. The set of model parameters is denoted as $\theta = \{r_i, a_{ij} : i, j \in 1, ..., n_s\}$.

#### 2 Maximum A Posteriori (MAP) parameter estimation

Model predictions for each sample, indexed by $i$, are determined by numerically integrating equation 1 from time zero to a sample specific end-point $t_i$ to give an estimate of species abundances, denoted as $\mathbf{x}_i$. Measured values are denoted as $\hat{\mathbf{x}}_i$, which are assumed to be corrupted by zero-mean Gaussian random noise,

$$\hat{\mathbf{x}}_i = \mathbf{x}_i + \varepsilon, \qquad \varepsilon \sim \mathcal{N}(\mathbf{0}, \mathbf{\Sigma}_x) \tag{2}$$

where $\mathbf{\Sigma}_x$ is the covariance of the noise distribution. Given a set of measured species abundances, $\mathcal{D} = \{\hat{\mathbf{x}}_1, ..., \hat{\mathbf{x}}_n\}$, with $n$ independent measurements, the likelihood of the data given the parameters is

$$p(\mathcal{D}|\theta, \mathbf{\Sigma}_x) = \prod_{i=1}^{n} \mathcal{N}(\hat{\mathbf{x}}_i | \mathbf{x}_i, \mathbf{\Sigma}_x). \tag{3}$$

Given a prior on parameters, $p(\theta) = \mathcal{N}(\theta|\theta_0, \mathbf{\Sigma}_0)$, the posterior parameter density is proportional to the product of the likelihood and the prior,

$$p(\theta|\mathcal{D}, \mathbf{\Sigma}_x, \mathbf{\Sigma}_0) \propto \prod_{i=1}^{n} \mathcal{N}(\hat{\mathbf{x}}_i | \mathbf{x}_i, \mathbf{\Sigma}_x) \mathcal{N}(\theta|\theta_0, \mathbf{\Sigma}_0). \tag{4}$$

Ignoring constants independent of $\theta$, we define the cost function $\mathcal{L}(\theta)$ as the negative log of equation 4,

$$\mathcal{L}(\theta) \overset{def}{=} \sum_{i=1}^{n} (\hat{\mathbf{x}}_i - \mathbf{x}_i)^T \mathbf{\Sigma}_x^{-1} (\hat{\mathbf{x}}_i - \mathbf{x}_i) + (\theta - \theta_0)^T \mathbf{\Sigma}_0^{-1} (\theta - \theta_0). \tag{5}$$

Because equation 5 is the proportional to the negative log posterior parameter distribution, minimizing this function with respect to $\theta$ gives the Maximum A Posteriori (MAP) estimate of model parameters,

$$\theta_{MAP} = \underset{\theta}{\operatorname{argmin}} \, \mathcal{L}(\theta). \tag{6}$$

#### 3 Gradient computation using forward sensitivity equations

To determine $\theta_{MAP}$, we use the MINIMIZE function provided by the Python package SCIPY using the NEWTON-CG method. This approach benefits significantly from providing the gradient of Eq. 5 computed using the forward sensitivity equations. The gradient of Eq. 5 is given by

$$\nabla_\theta \mathcal{L}(\theta) = \sum_{i=1}^{n} (\mathbf{x}_i - \hat{\mathbf{x}}_i)^T \mathbf{\Sigma}_x^{-1} \mathbf{Z}_i + (\theta - \theta_0)^T \mathbf{\Sigma}_0^{-1}, \tag{7}$$

where $\mathbf{Z}_i = \nabla_\theta \mathbf{x}_i$ is the model sensitivity matrix. Evaluating the elements of the sensitivity matrix can be done analytically using the forward sensitivity equations [2],

$$\frac{\mathrm{d}\mathbf{Z}}{\mathrm{d}t} = \left(\nabla_\mathbf{x} \frac{\mathrm{d}\mathbf{x}}{\mathrm{d}t}\right) \mathbf{Z} + \nabla_\theta \frac{\mathrm{d}\mathbf{x}}{\mathrm{d}t}. \tag{8}$$

Integration of equation 8 must be done simultaneously with $\frac{\mathrm{d}\mathbf{x}}{\mathrm{d}t}$ using a numerical differential equation solver. For the gLV model given by equation 1,

$$\left[\nabla_\mathbf{x} \frac{\mathrm{d}\mathbf{x}}{\mathrm{d}t}\right]_{ij} = x_i A_{ij} + \delta_{ij}\left(r_i + \sum_{l=1}^{n_s} A_{il} x_l\right). \tag{9}$$

Breaking up the gradient $\nabla_\theta \frac{\mathrm{d}\mathbf{x}}{\mathrm{d}t}$ into separate components for $r_j$ and $A_{jk}$ gives

$$\left[\nabla_\mathbf{r} \frac{\mathrm{d}\mathbf{x}}{\mathrm{d}t}\right]_{ij} = x_i \delta_{ij} \tag{10}$$

and

$$\left[\nabla_A \frac{\mathrm{d}\mathbf{x}}{\mathrm{d}t}\right]_{ijk} = x_i \delta_{ij} x_k \tag{11}$$

where $\delta$ is the Kronecker delta ($\delta_{ij} = 1$ if $i = j$ and 0 otherwise).

#### 4 Approximation of posterior parameter distribution

The Laplace approximation is a widely used approach used to approximate a probability density function as a Gaussian centered at a mode of the true distribution [1]. With $p(\theta|\mathcal{D})$ as the true posterior parameter distribution and $\theta_{MAP}$ a mode of $p(\theta|\mathcal{D})$, we seek an approximate Gaussian distribution, $q(\theta)$, centered at $\theta_{MAP}$. Let $f(\theta)$ denote the non-normalized true posterior parameter distribution,

$$p(\theta|\mathcal{D}) = \frac{f(\theta)}{C},$$

where $C$ is a normalizing constant. Defining the log of the approximating distribution, $\log q(\theta)$, as a second order Taylor series expansion of $\log f(\theta)$ about $\theta_{MAP}$ gives

$$\log f(\theta) \approx \log f(\theta_{MAP}) + \frac{1}{2}(\theta - \theta_{MAP})^T \nabla_\theta \nabla_\theta \log f(\theta)|_{\theta=\theta_{MAP}} (\theta - \theta_{MAP}) \stackrel{def}{=} \log q(\theta).$$

Taking the exponential gives

$$q(\theta) = f(\theta_{MAP}) \exp\left(-\frac{1}{2}(\theta - \theta_{MAP})^T \mathbf{\Sigma}_\theta^{-1} (\theta - \theta_{MAP})\right)$$

where

$$\mathbf{\Sigma}_{\theta}^{-1} = -\nabla_{\theta}\nabla_{\theta}\log f(\theta)|_{\theta=\theta_{MAP}}.$$

Because the cost function, $\mathcal{L}(\theta)$, was defined as the negative log of the posterior parameter distribution, the covariance matrix of the approximate distribution, $q(\theta)$, is given by

$$\begin{aligned}\mathbf{\Sigma}_{\theta}^{-1} &= \nabla_{\theta}\nabla_{\theta}\mathcal{L}(\theta) \\ &= \mathbf{\Sigma}_{0}^{-1} + \sum_{i=1}^{n}(\mathbf{x}_i - \hat{\mathbf{x}}_i)^T\mathbf{\Sigma}_{x}^{-1}\nabla_{\theta}\nabla_{\theta}\mathbf{x}_i + \sum_{i=1}^{n}\mathbf{Z}_i^T\mathbf{\Sigma}_{x}^{-1}\mathbf{Z}_i \\ &\approx \mathbf{\Sigma}_{0}^{-1} + \sum_{i=1}^{n}\mathbf{Z}_i^T\mathbf{\Sigma}_{x}^{-1}\mathbf{Z}_i,\end{aligned} \tag{12}$$

where the sum over the residuals, $(\mathbf{x}_i - \hat{\mathbf{x}}_i)$, is approximated as equal to zero. The Laplace approximated posterior parameter distribution is then

$$p(\theta|\mathcal{D}, \mathbf{\Sigma}_x, \mathbf{\Sigma}_0) \approx q(\theta) = \mathcal{N}(\theta|\theta_{MAP}, \mathbf{\Sigma}_{\theta}). \tag{13}$$

#### 5 Wald test for parameter significance

For a parameter $\theta_i$, we use a two-tailed Wald test to test whether the parameter is significantly constrained to be non-zero by the experimental data. Intuitively, the Wald test compares the parameter mean to its standard deviation in order to evaluate whether the peak of the posterior parameter distribution is significantly higher or lower than zero compared to the width of the distribution. We denote the null hypothesis as, $H_0 : \theta_i = 0$. The Wald test of size $\alpha = .05$ for parameter $i$ rejects $H_0$ when $|W_i| > z_{\alpha/2}$, where

$$W_i = \frac{[\theta_{MAP}]_i}{\sqrt{[\mathbf{\Sigma}_{\theta}]_{ii}}}. \tag{14}$$

The p-value for an observed Wald statistic, $W_i$, is given by $p = 2\Phi(-|W_i|)$, which is the probability of the Wald statistic being at least as large as the observed value under the null hypothesis [3].

#### 6 Estimation of posterior predictive distribution

Given a posterior parameter distribution, the posterior predictive distribution is found by marginalizing model predictions with respect to parameters,

$$p(\hat{\mathbf{x}}|\mathcal{D}, \mathbf{\Sigma}_x, \mathbf{\Sigma}_0) = \int p(\hat{\mathbf{x}}|\theta, \mathbf{\Sigma}_x)p(\theta|\mathcal{D}, \mathbf{\Sigma}_x, \mathbf{\Sigma}_0)\mathrm{d}\theta. \tag{15}$$

Linearizing the model prediction with respect to the model parameters around $\theta_{\mathrm{MAP}}$ gives

$$\mathbf{x}(\theta) \approx \mathbf{x}(\theta_{\mathrm{MAP}}) + \mathbf{Z}(\theta - \theta_{\mathrm{MAP}}), \tag{16}$$

which allows for the following analytical solution to equation 15

$$p(\hat{\mathbf{x}}|\mathcal{D}, \mathbf{\Sigma}_x, \mathbf{\Sigma}_0) = \mathcal{N}(\hat{\mathbf{x}}|\mathbf{x}(\theta_{\mathrm{MAP}}),\ \mathbf{\Sigma}_x + \mathbf{Z}\mathbf{\Sigma}_{\theta}\mathbf{Z}^T). \tag{17}$$

This approximation is referred to as the linear-Gaussian approximation [1]. When reporting predicted uncertainty, species abundances $i$ are reported as $x_i(\theta_{\mathrm{MAP}}) \pm \sigma_i$, where

$$\sigma_i = \sqrt{[\mathbf{\Sigma}_x + \mathbf{Z}\mathbf{\Sigma}_{\theta}\mathbf{Z}^T]_{ii}} \tag{18}$$

#### 7 Bayesian experimental design

The goal of Bayesian experimental design is to select an experimental design, denoted as $\mathbf{q} = \{q_1, ..., q_n\}$ that results in data, $\mathcal{D}(\mathbf{q}) = \{\hat{\mathbf{x}}(q_1), ..., \hat{\mathbf{x}}(q_n)\}$, that is expected to minimize the spread of the parameter distribution according to Bayes' formula,

$$p(\theta|\mathcal{D}(\mathbf{q})) = \frac{p(\mathcal{D}(\mathbf{q})|\theta)p(\theta)}{p(\mathcal{D}(\mathbf{q}))}. \tag{19}$$

The variable $q_i$ is introduced to denote a particular experimental condition, which could for example be the specification of species to inoculate and time points to measure species abundances after inoculation. It is important to note that the prior $p(\theta)$ could be the posterior determined from performing inference on previous data. In order to select for an optimal experimental design, we define a utility function, $U(\mathbf{q})$, as the expected information gain (EIG) that would result from updating the current model with a new dataset $\mathcal{D}(\mathbf{q})$. The EIG is the expected Kullback-Leibler divergence between the parameter posterior and the parameter prior distributions, which has an analytical expression in the case of the linear-Gaussian model

$$\begin{aligned}
U_{\text{EIG}}(\mathbf{q}) &= -\int\int \ln p(\mathcal{D}(\mathbf{q}))p(\mathcal{D}(\mathbf{q})|\theta)\mathrm{d}\mathbf{y}p(\theta)\mathrm{d}\theta \\
&= -\int \ln p(\mathcal{D}(\mathbf{q}))\int p(\mathcal{D}(\mathbf{q}),\theta)\mathrm{d}\theta\mathrm{d}\mathbf{y} \\
&= -\int \ln p(\mathcal{D}(\mathbf{q}))p(\mathcal{D}(\mathbf{q}))\mathrm{d}\mathbf{y} \\
&= \ln|\mathbf{\Sigma}_x + \sum_{i=1}^{n}\mathbf{Z}(q_i)\mathbf{\Sigma}_0\mathbf{Z}(q_i)^T|
\end{aligned} \tag{20}$$

where the probability distribution of the data is determined by marginalizing over the parameter distribution

$$p(\mathcal{D}(\mathbf{q})) = \prod_{i=1}^{n} p(\hat{\mathbf{x}}(q_i)) = \prod_{i=1}^{n}\mathcal{N}(\hat{\mathbf{x}}_i|\mathbf{x}(\theta_0), \Sigma_x + \mathbf{Z}(q_i)\mathbf{\Sigma}_0\mathbf{Z}(q_i)^T) \tag{21}$$

and $\mathbf{Z}(q_i) = \nabla_\theta \mathbf{x}(q_i)$ is the gradient of the model prediction of the outcome corresponding to experimental condition $q_i$ with respect to model parameters. Given a set of all possible experimental conditions, denoted as $\mathcal{Q}$, we use a Greedy search algorithm to determine the subset of $n$ experimental conditions that maximize Eq. 20, where the first selected experimental condition is given by,

$$q_1 = \underset{q\in\mathcal{Q}}{\text{argmax}} \quad \ln|\mathbf{\Sigma}_x + \mathbf{Z}(q)\mathbf{\Sigma}_0\mathbf{Z}(q)^T| \tag{22}$$

and subsequent experimental conditions are selected according to

$$q_{i+1} = \underset{q\in\mathcal{Q}}{\text{argmax}} \quad \ln|\mathbf{\Sigma}_x + \mathbf{Z}(q)\mathbf{\Sigma}_i\mathbf{Z}(q)^T| \tag{23}$$

where

$$\mathbf{\Sigma}_i = \mathbf{\Sigma}_{i-1} - \mathbf{\Sigma}_{i-1}\mathbf{Z}(q_i)^T(\Sigma_x + \mathbf{Z}(q_i)\mathbf{\Sigma}_{i-1}\mathbf{Z}(q_i)^T)^{-1}\mathbf{Z}(q_i)\mathbf{\Sigma}_{i-1} \tag{24}$$

until $i = n$, where $n$ is the total number of conditions that are experimentally practical to collect.

#### 8 Hyper-parameter optimization

Optimization of hyper-parameters using only the training data is often called *empirical Bayes* [1]. Model hyper-parameters include the covariance of the noise distribution $\mathbf{\Sigma}_x$ and the covariance of the prior parameter distribution $\mathbf{\Sigma}_0$, which is a diagonal matrix with entries given by $\alpha \in \mathbb{R}_+^{n_\theta}$. To determine a set of optimal hyper-parameters, $\xi^* = \{\mathbf{\Sigma}_x^*, \mathbf{\Sigma}_0^*\}$, we seek $\xi$ that maximizes the marginal likelihood function given by

$$p(\mathcal{D}|\xi) = \int_\theta p(\mathcal{D}, \theta|\xi) d\theta. \tag{25}$$

We can use the *expectation maximization* (EM) algorithm to update $\xi^{(l+1)}$, which involves maximizing the expected log likelihood,

$$\xi^{(l+1)} = \underset{\xi}{\text{argmax}}\ \mathbb{E}_{\theta|\mathcal{D},\xi^{(l)}}[\ln p(\mathcal{D}, \theta|\xi)], \tag{26}$$

followed by re-evaluation of the posterior parameter distribution $p(\theta|\mathcal{D}, \xi^{(l+1)})$. The process of maximizing Eq 26 and updating the posterior parameter distribution is repeated until convergence of the marginal likelihood function given by Eq 30. With $\ln p(\mathcal{D}, \theta) = \ln p(\mathcal{D}|\theta) + \ln p(\theta)$, Eq 26 becomes

$$\mathbb{E}_{\theta|\mathcal{D},\xi^{(l)}}[\ln p(\mathcal{D}, \theta|\xi)] = \mathbb{E}_{\theta|\mathcal{D},\xi^{(l)}}[\ln p(\mathcal{D}|\theta, \xi)] + \mathbb{E}_{\theta|\mathcal{D},\xi^{(l)}}[\ln p(\theta|\xi)]. \tag{27}$$

The first term is the expectation of the log likelihood,

$$\begin{aligned}\mathbb{E}_{\theta|\mathcal{D},\xi^{(l)}}[\ln p(\mathcal{D}|\theta, \xi)] = &-\frac{1}{2}\sum_{i=1}^{n} \mathbb{E}_{\theta|\mathcal{D},\xi^{(l)}}\left[(\hat{\mathbf{x}}_i - \mathbf{x}_i)^T \mathbf{\Sigma}_x^{-1}(\hat{\mathbf{x}}_i - \mathbf{x}_i)\right] \\ &- \frac{n}{2}\ln\left(2\pi \det \mathbf{\Sigma}_x\right).\end{aligned}$$

linearizing the model with respect to $\theta$ about $\theta_{\text{MAP}}$ and evaluating the expectation gives

$$\begin{aligned}\mathbb{E}_{\theta|\mathcal{D},\xi^{(l)}}[\ln p(\mathcal{D}|\theta, \xi)] = &-\frac{1}{2}\sum_{i=1}^{n} (\hat{\mathbf{x}}_i - \mathbf{x}_i)^T \mathbf{\Sigma}_x^{-1} (\hat{\mathbf{x}}_i - \mathbf{x}_i) \\ &+ \text{Tr}\left(\mathbf{\Sigma}_x^{-1}\mathbf{Z}_i\mathbf{\Sigma}_\theta\mathbf{Z}_i^T\right) - \frac{n}{2}\ln\left(2\pi \det \mathbf{\Sigma}_x\right).\end{aligned}$$

Evaluating the second term in Eq 27 gives

$$\mathbb{E}_{\theta|\mathcal{D},\xi^{(l)}}[\ln p(\theta|\xi)] = -\frac{1}{2}\sum_{k=1}^{n_\theta}\left(-\ln\,[\alpha]_k - ([(\theta_{\text{MAP}} - \theta_0)^2]_k + [\mathbf{\Sigma}_\theta]_{kk})/[\alpha]_k - \ln 2\pi\right)$$

Update equations for $\xi$ are found by taking the derivative of Eq 27 with respect to $\mathbf{\Sigma}_x$ and $[\alpha]_k$ and solving for $\mathbf{\Sigma}_x^{(l+1)}$ and $[\alpha^{(l+1)}]_k$,

$$\mathbf{\Sigma}_x^{(l+1)} = \frac{1}{n}\sum_{i=1}^{n}(\hat{\mathbf{x}}_i - \mathbf{x}_i)(\hat{\mathbf{x}}_i - \mathbf{x}_i)^T + \mathbf{Z}_i\mathbf{\Sigma}_\theta\mathbf{Z}_i^T, \tag{28}$$

$$[\alpha^{(l+1)}]_k = [(\theta_{\text{MAP}} - \theta_0)^2]_k + [\mathbf{\Sigma}_\theta]_{kk}. \tag{29}$$

Model hyper-parameters are updated until convergence of the log of the model evidence (i.e. marginal likelihood), which is approximated using the Laplace approximation as

$$\begin{aligned}\ln p(\mathcal{D}|\xi) \approx & -\frac{1}{2}\ln \det \boldsymbol{\Sigma}_0 - \frac{n}{2}\ln \det \boldsymbol{\Sigma}_x + \frac{1}{2}\ln \det \boldsymbol{\Sigma}_\theta \\ & -\frac{1}{2}\theta_{\mathrm{MAP}}^T \boldsymbol{\Sigma}_0^{-1}\theta_{\mathrm{MAP}} - \frac{1}{2}\sum_{i=1}^{n}(\hat{\mathbf{x}}_i - \mathbf{x}_i)^T \boldsymbol{\Sigma}_x^{-1}(\hat{\mathbf{x}}_i - \mathbf{x}_i). \end{aligned} \tag{30}$$

[2] Brian Munsky, William S Hlavacek, and Lev S Tsimring. *Quantitative biology: theory, computational methods, and models.* MIT Press, 2018.

[3] Larry Wasserman. *All of statistics: a concise course in statistical inference*, volume 26. Springer, 2004.

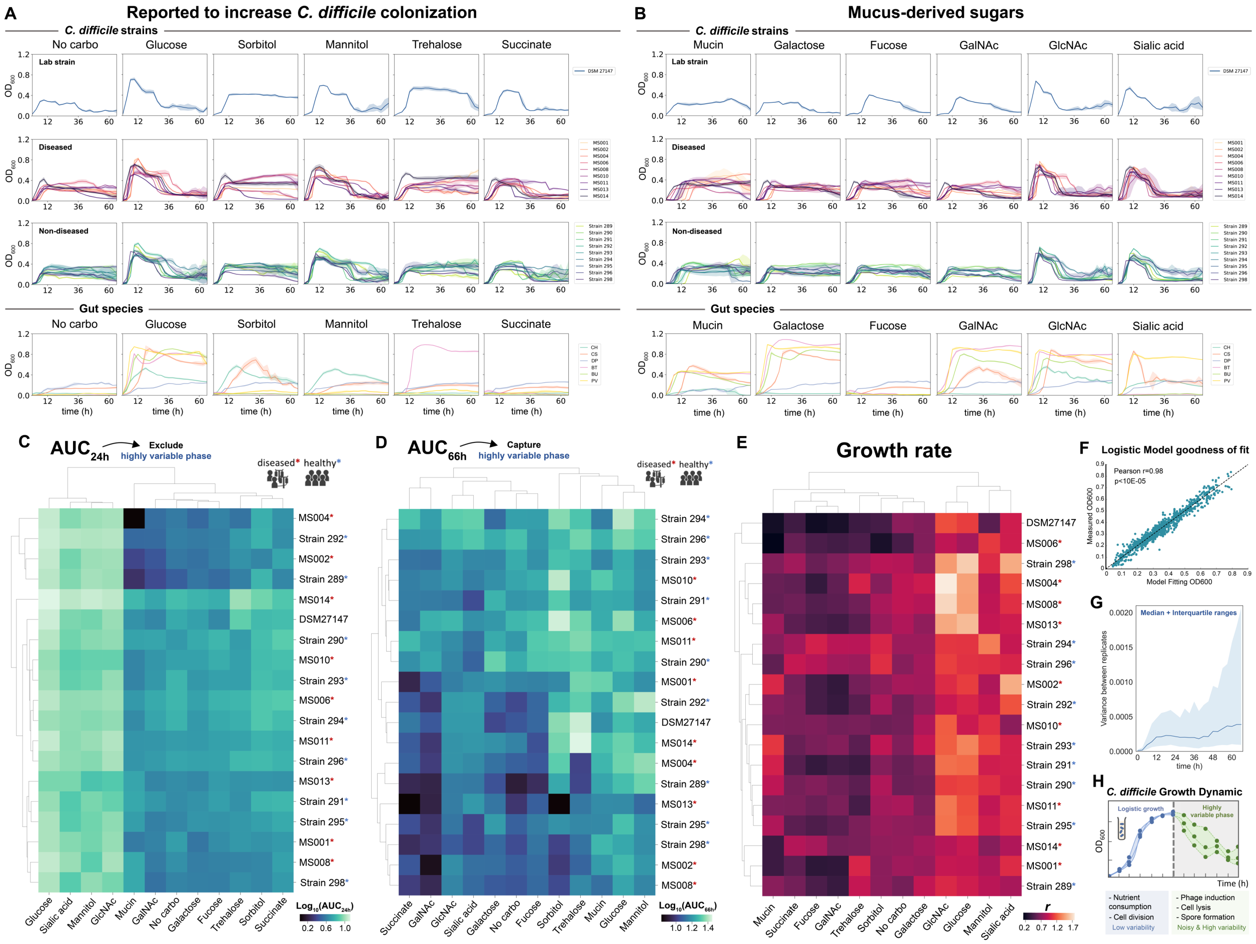


**Supplementary Figure 1**. ***C. difficile* monoculture growth profile. a-b**, $OD_{600}$ measurement of *C. difficile* strains and gut bacteria grown in DM29 defined media (See **Methods**) supplemented with different carbohydrates that were previously reported to increase *C. difficile* colonization (**a**) and mucus-derived sugars (**b**). Data were shown as mean and 95% c.i. (shading), n = 3 biological replicates. **c-d**, Biclustering heatmap of the average integral $OD_{600}$ or the Area Under the Curve (AUC) extracted from monoculture growth data of *C. difficile* in media supplemented with different carbohydrate sources after 24 h of growth ($AUC_{24h}$) (**c**) and 66 h of growth ($AUC_{66h}$) (**d**). Strains marked with red asterisks were isolated from diseased patients whereas the ones marked with blue asterisks were isolated from healthy individuals. **e**, Biclustering heatmap of the maximum growth rate of the *C. difficile* isolates. The experimental data were fitted into the logistic growth model as shown in **Fig. 1a. f**, Goodness of fit of the monoculture logistic growth model. The x-axis shows the $OD_{600}$ from model fitting, whereas y-axis shows the measured $OD_{600}$ from all time points across all 12 media conditions. Pearson's correlation coefficient (*r*) and *p*-values are shown, which were computed using the pearsonr from the scipy package in Python. Dashed line indicates y=x. **g**, Variance between biological replicates of all *C. difficile* strains over time. The blue line and shaded area are the median and interquartile ranges (n=228 per timepoint, i.e. 19 strains grown in 12 media). **h**, Schematic of *C. difficile* growth dynamic. The first stage marked the logistic bacterial growth with low variability between replicates. The second stage is associated with the massive $OD_{600}$ reduction and high variability between replicates due to unpredictable events such as phage induction, cell lysis, and spore formation.

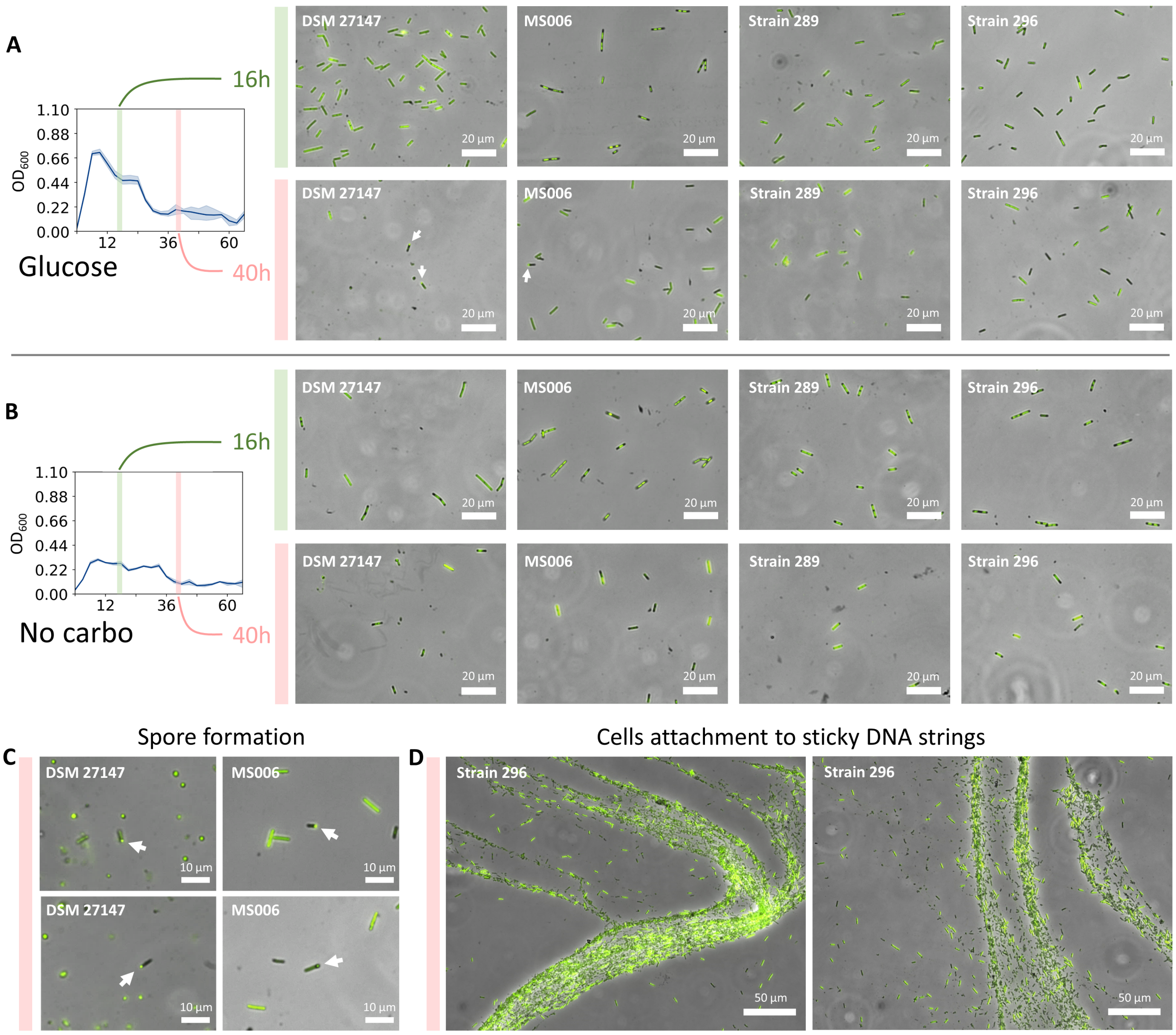


**Supplementary Figure 2**. **Fluorescence microscopy of *C. difficile* strains in different growth phases. a-b**, Four different *C. difficile* strains (DSM 27147, MS006, Strain 289, Strain 296) were either grown anaerobically in media with glucose as a carbohydrate source (**a**) or media without carbohydrate (**b**), sampled at two time points (16 h and 40 h), and stained with SYBR Green dye. White arrows marked spore formation events. **c-d**, Representative images showing the formation of spores as indicated by the white arrows (**c**), and cells attached to DNA strings (**d**), which were only observed in t = 40 h.

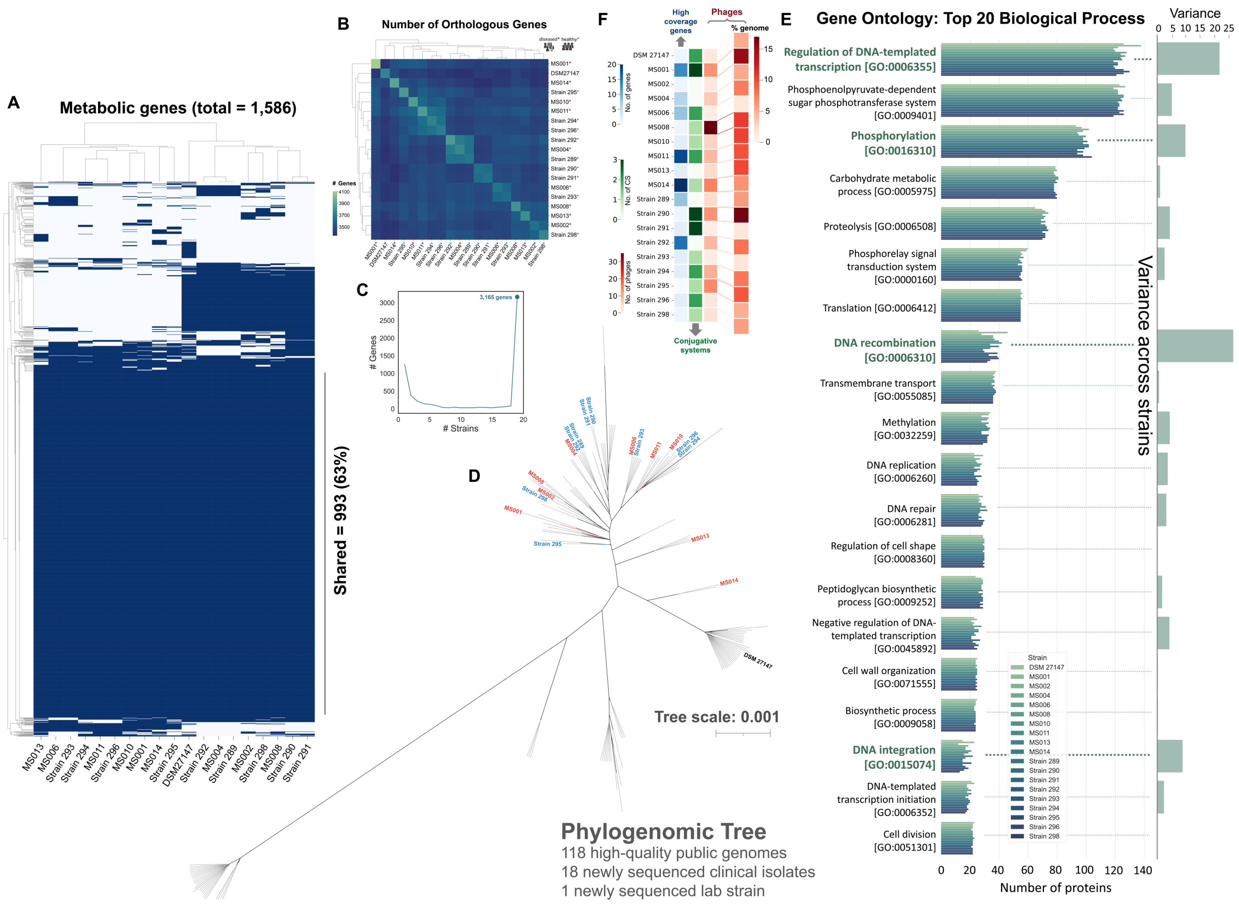


**Supplementary Figure 3**. **Genotypic characterization of the *C. difficile* isolates**. **a**, Biclustering heatmap showing the presence and absence of metabolic genes identified across the 19 *C. difficile* strains. The metabolic genes were determined by constructing strain-specific genome scale models for the 19 *C. difficile* strains (See **Methods**) and each reaction in the model is regarded as one gene. The rows indicate the metabolic genes, and the columns indicate the *C. difficile* strains. Blue means gene present and white means gene absent. 63% of the whole metabolic genes are shared among the 19 strains. **b**, Biclustering heatmap of the number of orthologous genes between isolate pairs. The horizontal boxes indicated the total number of genes in a specific isolate. Strains marked with red asterisks were isolated from diseased patients whereas the ones marked with blue asterisks were isolated from healthy individuals. **c**, Gene conservation across isolates demonstrates functional diversity within *C. difficile*, with a core genome of 3,165 genes. **d**, Phylogenomic tree constructed using 118 high-quality public *C. difficile* genomes (**Table S4**) and our newly sequenced *C. difficile* isolates. Black, blue, and red color labels correspond to the DSM27147 strain, isolates from healthy individuals, and isolates from diseased individuals, respectively. Each dashed line on the panel represents one strain. **e**, Top 20 biological processes observed across all isolates with the number of annotated proteins in each process. Bar plots on the right show the variance across isolates (n=19). Processes that have high variance across isolates were bolded with green color. **f**, Summary of the high coverage genes, conjugative systems (CS), and phages in specific *C. difficile* strains. The three heatmaps on the left show the number of high-coverage genes (blue), CS (green), and phages (red) found in the *C. difficile* genomes, whereas the larger heatmap on the right corresponds to the percentage of the genome that the phages occupy. Details are available in **Table S5-7** and **Fig. S5**.

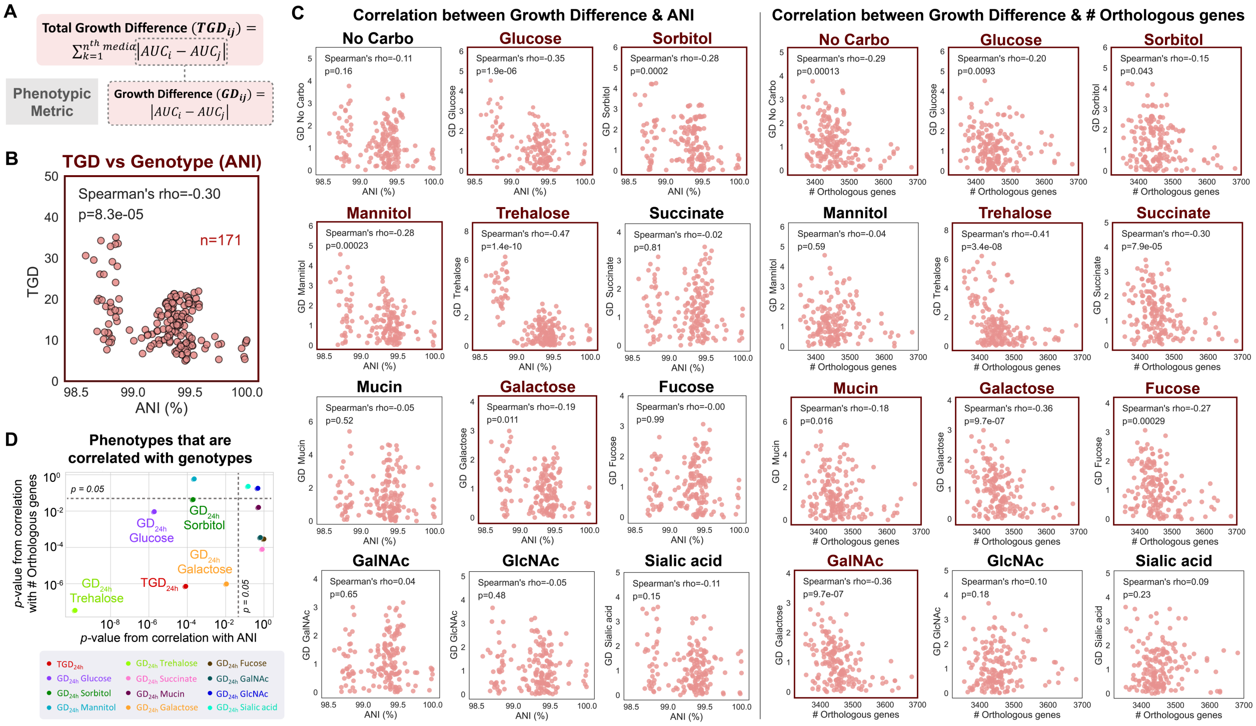


**Supplementary Figure 4. Correlation between growth and the genotypes of *C. difficile* isolates. a,** Mathematical formula to calculate phenotypic metrics: Growth Difference (GD) in specific media and the Total Growth Difference (TGD). The GD between two isolates ($GD_{ij}$) is the difference in the AUC from 24 h of growth in specific media, whereas the TGD between two isolates ($TGD_{ij}$) is the sum of all AUC differences from 24 h of growth in the twelve media. **b**, Scatter plot of TGD between isolate pairs and ANI between isolate pairs. **c**, Scatter plot of Growth Difference (GD) between isolate pairs calculated using AUC from 24 hours of growth and ANI (left) or the number of orthologous genes (right). Spearman's rho and *p*-value are shown (n=171 data points for each plot), which were computed using the spearmanr from the scipy package in Python. Plots outlined with red color marked statistically significant correlations. **d,** Scatter plot of the Spearman's *p*-values from correlation with ANI and Spearman's *p*-values from correlation with the number of orthologous genes for each phenotypic metric (denoted with different colors). The dotted lines marked the *p*=0.05 threshold.

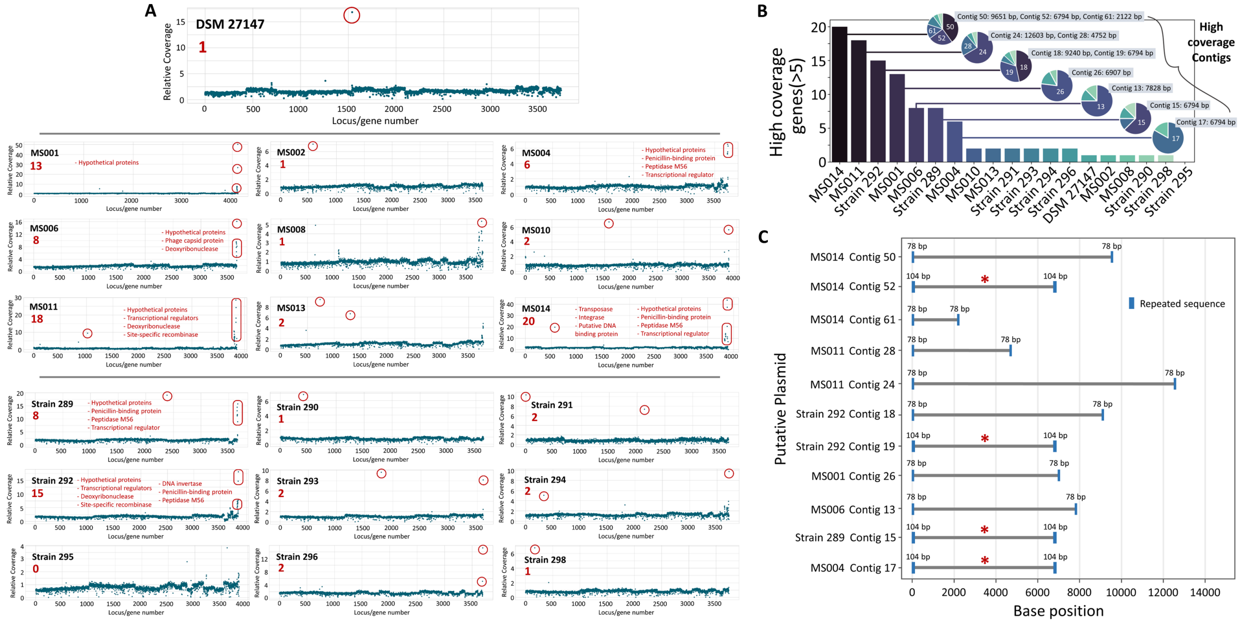

**Supplementary Figure 5. Putative plasmids on the *C. difficile* strains. a,** Scatter plot of the relative coverage of all genes in the *C. difficile* isolates. The x-axis indicates specific genes, whereas the y-axis is the relative coverage calculated by the total number of aligned sequencing raw reads divided by gene length. Data points with high relative coverage (>5) are highlighted in red, and the gene annotations were shown. **b**, Bar plot of the number of genes with high coverage (>5) across different *C. difficile* isolates. The pie charts on the right side show the contigs where the high coverage genes were detected. For instance, most of the high coverage genes from MS014 were detected in Contig 50, 52, and 61, with Contig lengths of 9651, 6794, and 2122 base pairs respectively. **c**, 11 Putative plasmids (>2kbp, >5-fold increase in coverage versus the median of the genome) from 7 isolates were detected with the presence of repeated termini. Red asterisks marked the same plasmid that is present in four different genetically distant *C. difficile* isolates from different patients. A homology search of the shared plasmid by BLAST showed a partial identity match (~50%) to another plasmid that was engineered as an *E. coli* – *C. difficile* shuttle vector [1-4] and a 100% identity match to a plasmid found in *C. difficile* clinical isolates from two other distant regions, Perth (Australia) and Pennsylvania (United States). This implies that horizontal gene transfer yielded distinct strains harboring the same plasmid or that strains harboring the plasmid evolved from a common ancestor.

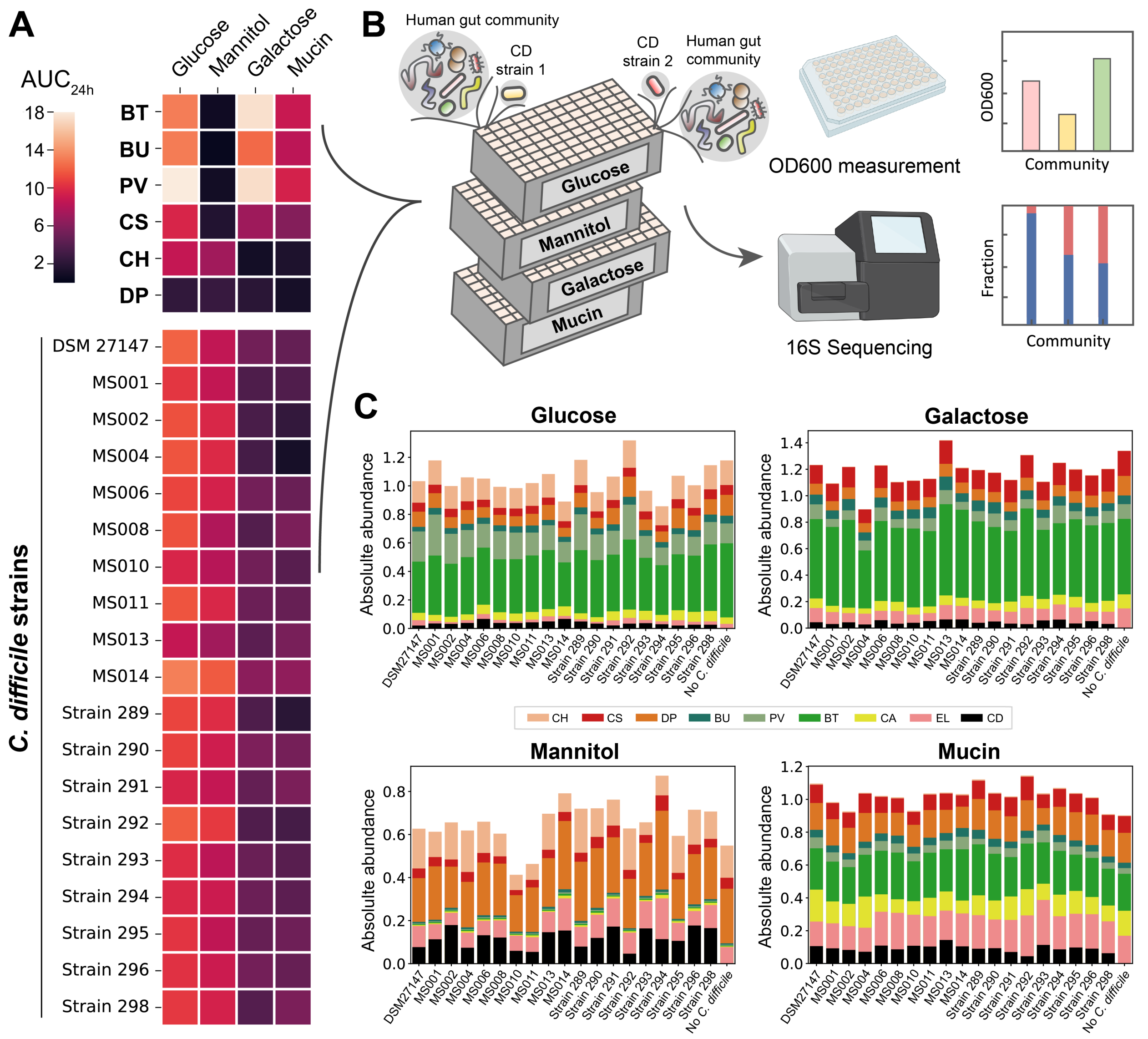


**Supplementary Figure 6. Growth of *C. difficile* strains with the human gut community in media supplemented with single carbohydrate sources. a**, Heatmap of the Area Under the Curve (AUC) extracted from monoculture growth data of 6 gut bacteria and 19 *C. difficile* strains in media supplemented with four different carbohydrate sources after 24 h of growth (See **Fig. S1a-b**). **b**, Different *C. difficile* isolates and the 8-member human gut community were grown in media containing four different carbohydrate sources; Glucose, galactose, mannitol, and mucin, representing different growth scenarios based on monoculture growth data. For instance, media containing glucose favors the growth of *C. difficile* and all other gut species, whereas media containing mannitol only favors the growth of *C. difficile* but not the other gut species. Synthetic communities are cultured in microtiter plates at an equal absolute abundance ratio in anaerobic conditions and incubated at 37°C for 24 h. The absolute abundance of each species is determined by measuring cell density at 600 nm ($OD_{600}$) and community composition using multiplexed 16S rRNA sequencing. **c**, Stacked bar plot of the composition of the 8-member community and *C. difficile* grown in media containing different carbohydrates as indicated. The composition of an 8-member community that was grown without *C. difficile* was shown as a control. Each bar represents the average absolute abundance of each species (n=3). Parts of the figure was generated using Biorender.

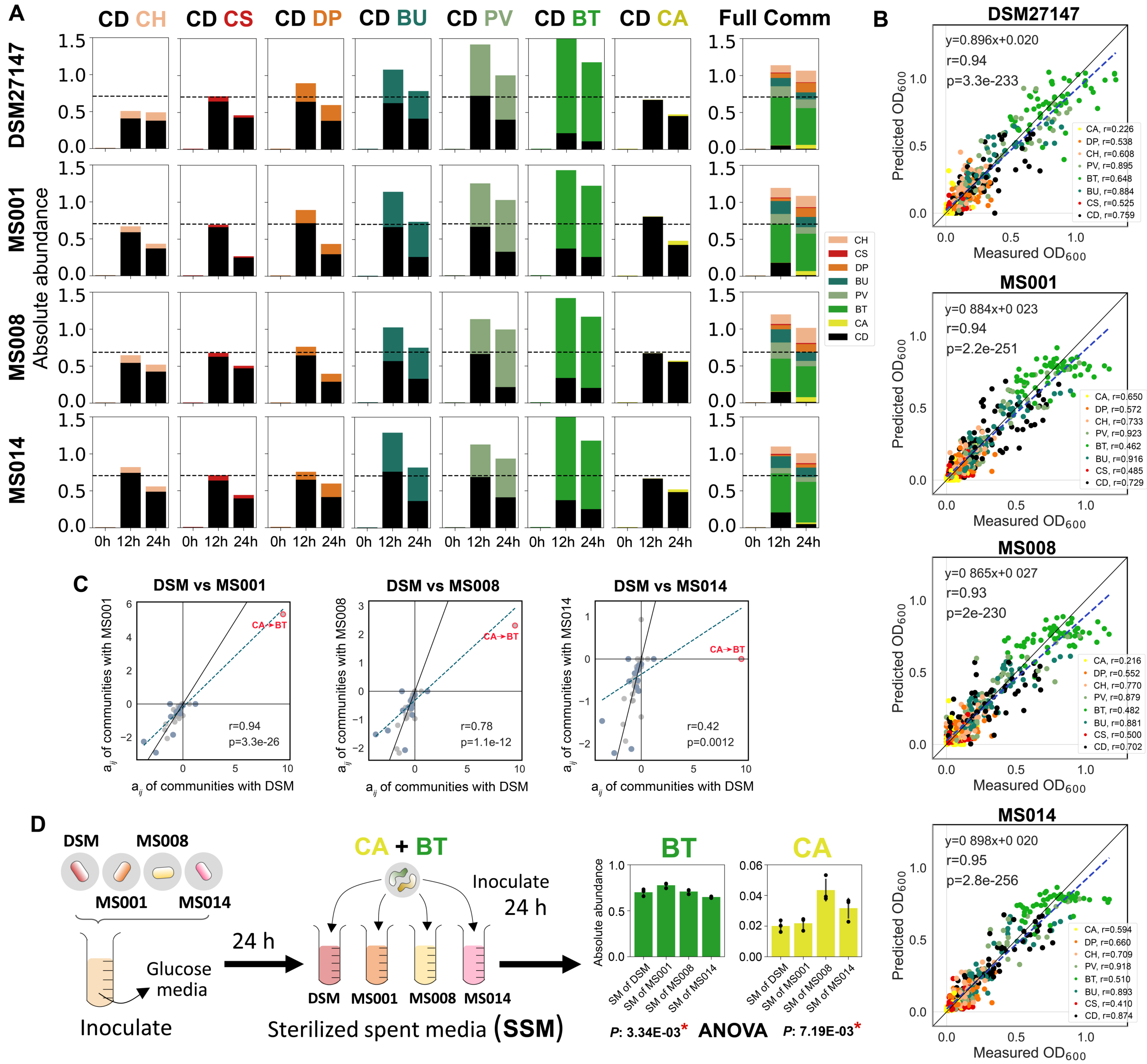


**Supplementary Figure 7. Growth of four representative *C. difficile* strains with the human gut community in the glucose media. a**, Absolute abundance (calculated $OD_{600}$) of pairwise communities containing different representative *C. difficile* strains over time when grown in the glucose media. Absolute abundance is the product of 16S relative abundance and community $OD_{600}$, and each bar represents the average absolute abundance of each species (n=3). The plots on the right side are the stacked bar plots of the full 8-member community. Horizontal dashed line indicates the $OD_{600}$ of C. difficile monoculture. **b**, Scatter plot of measured $OD_{600}$ versus predicted $OD_{600}$ for 2-8 species communities. The data used for parameter estimation is listed in **Table S8** (DATASET001). Colors indicate the species in the community that was measured and predicted. Blue dashed line indicates the linear regression between the mean measured $OD_{600}$ and the predicted $OD_{600}$ (See **Methods** for details). Pearson's correlation coefficient (*r*) and *p*-values are shown, which were computed using the pearsonr from the scipy package in Python. **c**, Scatter plots of the interspecies interaction coefficients ($a_{ij}$) between communities containing different *C. difficile* strains in the glucose media. Grey data points are interaction coefficients between two gut species, whereas blue data points are interaction coefficients between *C. difficile* and a gut species. Blue dashed line indicates the linear regression between the coefficients of the two communities. Pearson's correlation coefficient (*r*) and *p*-values are shown, which were computed using the pearsonr from the scipy package in Python. **d**, Growth of *C. aerofaciens* and *B. thetaiotaomicron* pairwise co-culture in *C. difficile* supernatants. Different *C. difficile* strains were grown in the glucose media for 24 h. The cultures were centrifuged, and the supernatant was filter sterilized and mixed with fresh media at an equal ratio to replenish the nutrients. CA-BT pair were grown in the four sterilized spent media from different *C. difficile* strains at an equal initial absolute abundance for 24 h. Each bar represents the average absolute abundance of each species, and the error bars represent s.d. (n=3). *P*-values from one-way ANOVA statistical test of the absolute abundance values of each species across pairwise communities grown in the spent media from different *C. difficile* strains were shown. Red asterisks indicate statistically significant p-values.

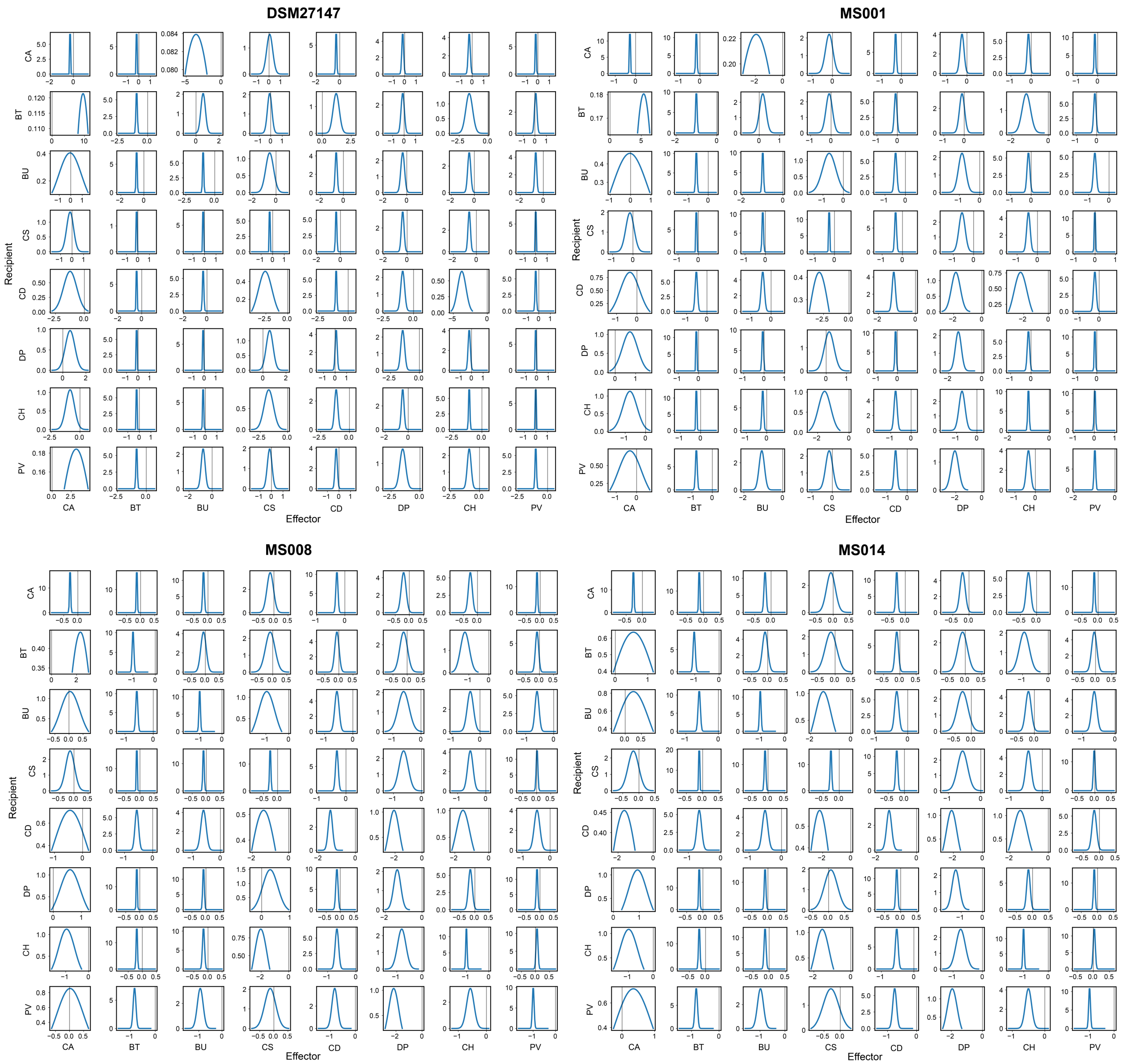


**Supplementary Figure 8. Parameter estimates for the gLV model in the glucose media.** The plots show the mean and variance of each gLV parameter ($a_{ij}$ values) for communities containing the four *C. difficile* strains (details can be found in the **Supplementary text**). Only parameters whose absolute values were significantly constrained to be non-zero based on the Wald test are shown in the interaction networks.

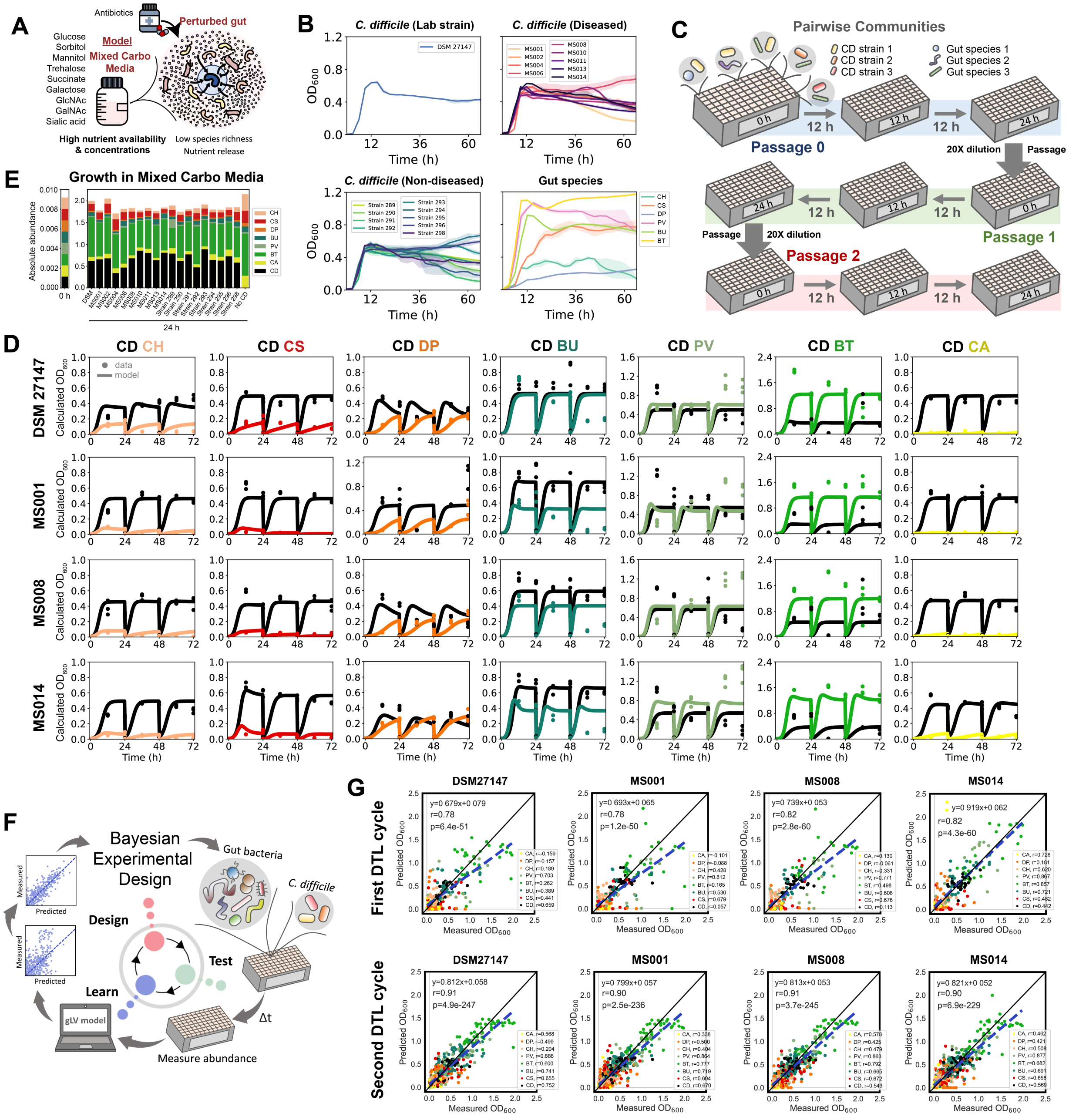


**Supplementary Figure 9. Growth of different *C. difficile* strains in monocultures and with human gut communities in mixed carbohydrates media. a**, Schematic of the mixed carbohydrates media that represents post-antibiotic perturbation where species richness is low and there are plenty of available nutrients to consume. **b**, $OD_{600}$ measurement of *C. difficile* strains and gut bacteria in the mixed carbohydrates media. Data were shown as mean and 95% c.i. (shading) (n=3). **c**, Schematic of the workflow for time-resolved abundance measurement of *C. difficile* and gut bacteria in pairwise communities. *C. difficile* strains were cultured at an equal initial absolute abundance ratio in the mixed carbohydrates media in microtiter plates under anaerobic conditions and incubated at 37°C. Every 12 h of growth, aliquots of the culture were taken for multiplexed 16S rRNA sequencing to determine community composition. The communities were passaged using a 1:20 dilution at 24 and 48 h to observe community assembly over three batch culture growth cycles and capture the longer-term behavior of the consortia. **d**,

Absolute abundance (calculated $OD_{600}$) of pairwise communities containing *C. difficile* over time for three growth cycles in the mixed carbohydrates media (n=3). Datapoints indicate experimental data replicates and lines indicate simulations using the gLV model (trained on DATASET002 in **Table S8**). Calculated $OD_{600}$ is the product of 16S relative abundance and community $OD_{600}$. **e**, Stacked bar plot of the absolute abundance of the 8-member human gut community and *C. difficile* grown in the mixed carbohydrates media at t=0 h and t=24 h. The *C. difficile* abundance at t=0 h is the average value across 19 strains. For communities after 24 h of growth, each bar represents the average absolute abundance of each species (n=3). **f**, Design–Test–Learn (DTL) cycle for model development in the mixed carbohydrates media. **g,** Model performance from the first DTL cycle using monoculture, pairwise, leave-one-out, and full communities' data to fit the gLV model (left), and second DTL cycle using additional data from the EDA (right). The plots are measured $OD_{600}$ versus predicted $OD_{600}$ of each species in the community containing different *C. difficile* strains. The data used for parameter estimation are listed in **Table S8** (DATASET002 for the first DTL cycle and DATASET003 for the second DTL cycle). Colors indicate the species in the community. Blue dashed line indicates the linear regression between the mean measured $OD_{600}$ and the predicted $OD_{600}$. Pearson's correlation coefficient (*r*) and *p*-values are shown, which were computed using the pearsonr from the scipy package in Python.

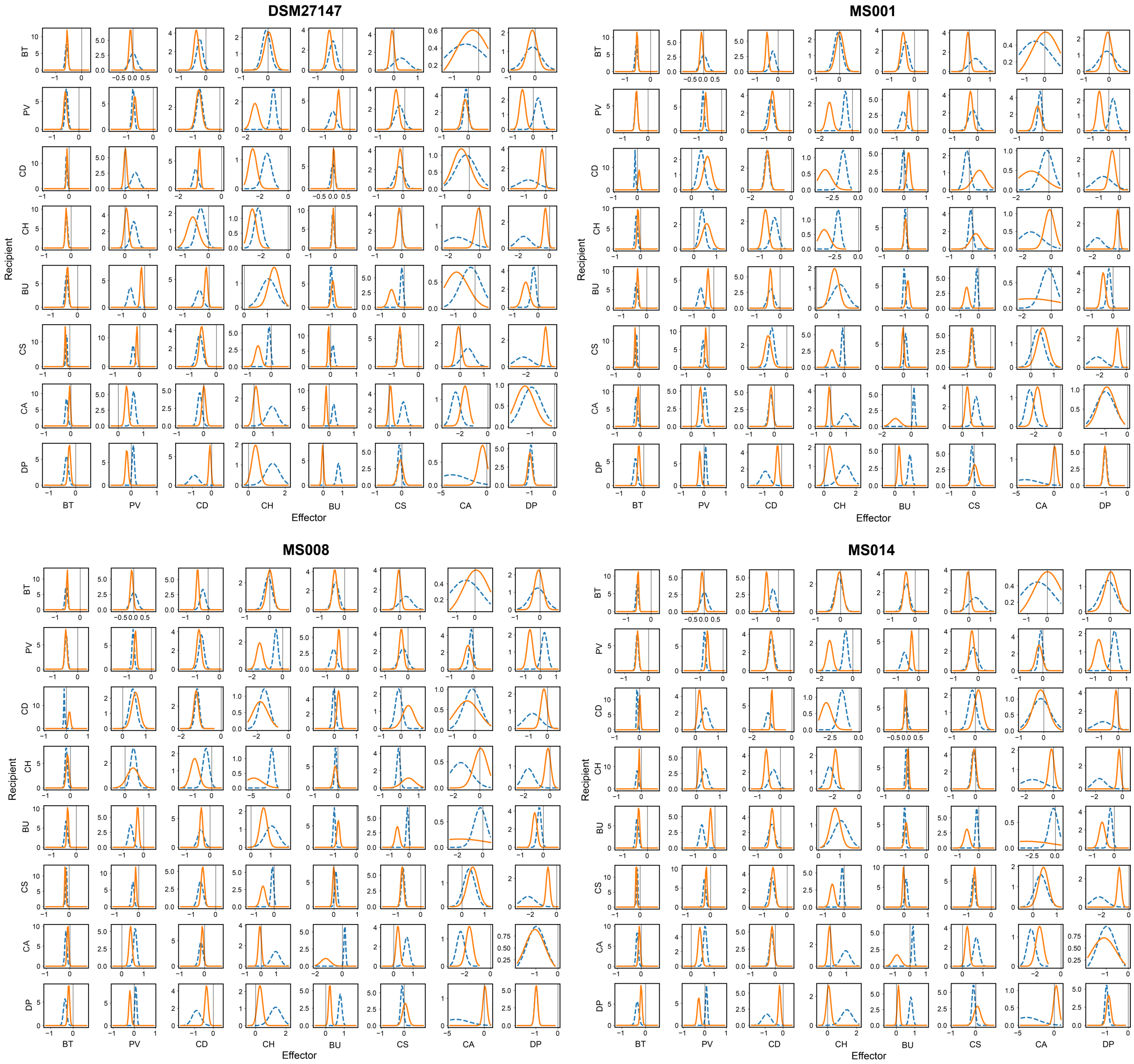


**Supplementary Figure 10. Parameter estimates for the gLV model in the mixed carbohydrates media.** The plots show the mean and variance of each gLV parameter ($a_{ij}$ values) for communities containing the four *C. difficile* strains (details can be found in the **Supplementary text**). Dashed blue lines indicate the parameter estimates from the first DTL cycle (**DATASET002, Table S8**) whereas the solid orange lines indicate the parameter estimates from the second DTL cycle (**DATASET003, Table S8**). Only parameters whose absolute values were significantly constrained to be non-zero based on the Wald test are shown in the interaction networks.

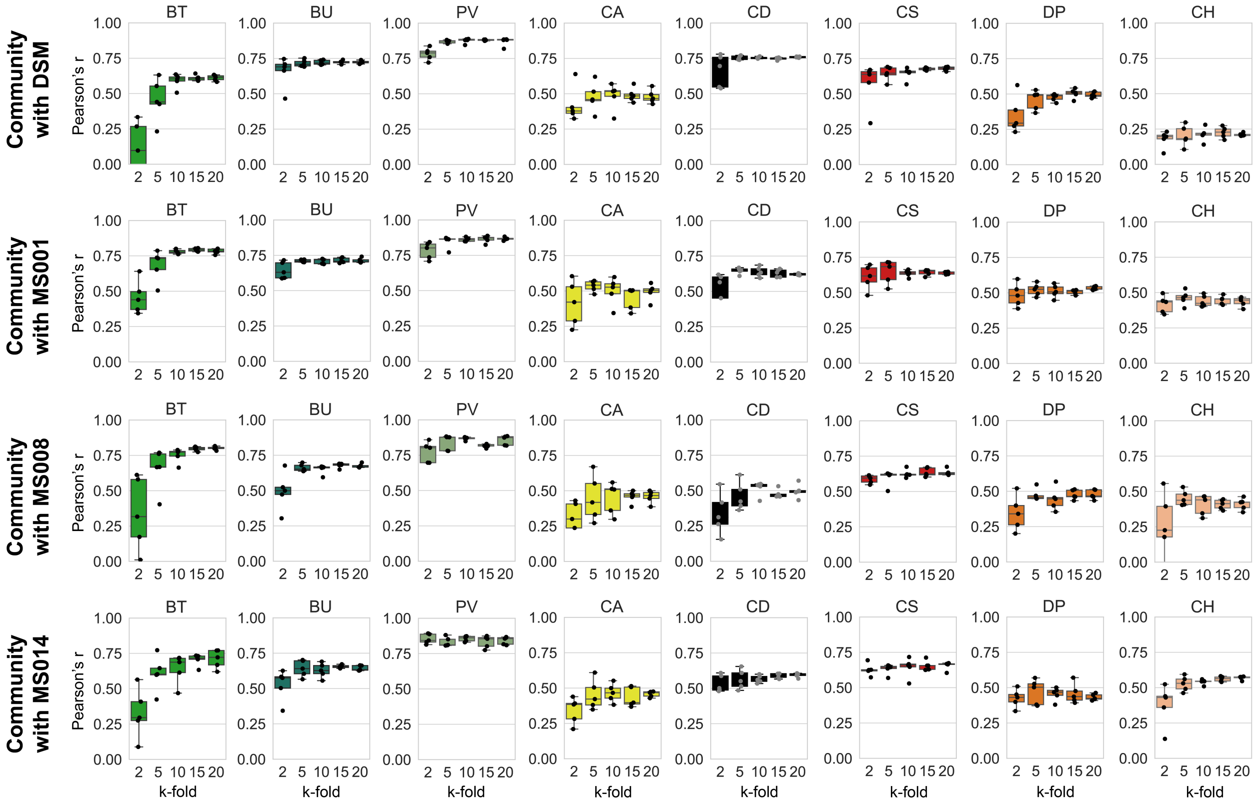


**Supplementary Figure 11. Sensitivity analysis of the gLV model trained on experimental data of communities grown in the mixed carbohydrates media.** The box plots showed the k-fold prediction performance of each species in the communities containing different *C. difficile* strains. The x-axis indicates the number of train/test splits ("k" in k-fold), and the y-axis is Pearson's correlation coefficient (*r*). Five repeated trials were performed for each k-fold prediction where the train/test splits were randomized.

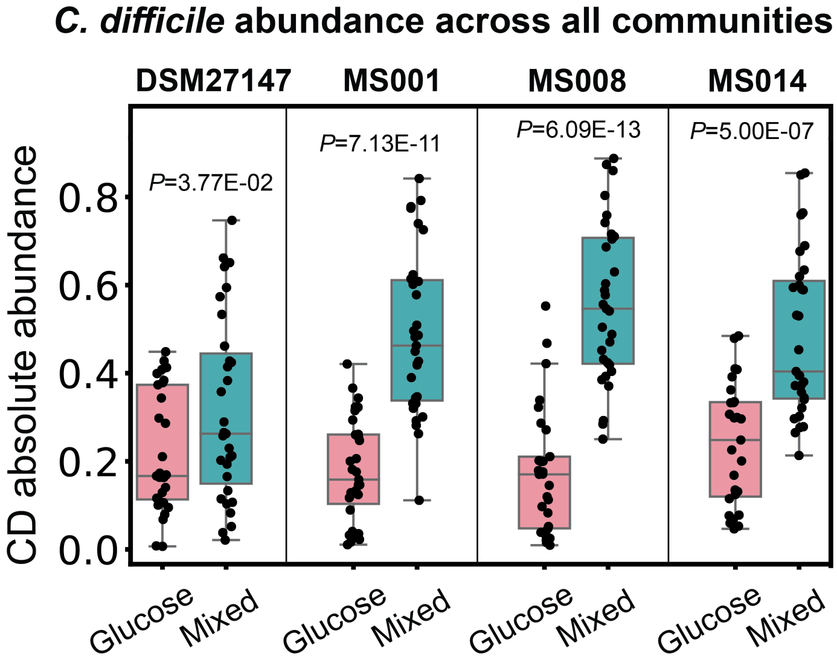


**Supplementary Figure 12. Comparison of *C. difficile* abundance when grown with the human gut communities in the mixed carbohydrates media and glucose media.** Box plot of *C. difficile* absolute abundance values at 24 h from all community growth data that were used to fit the gLV models in the glucose media (pink) and the mixed carbohydrates media (green). *P*-values from unpaired *t*-test of *C. difficile* abundance in glucose media vs. mixed carbohydrates media were shown.

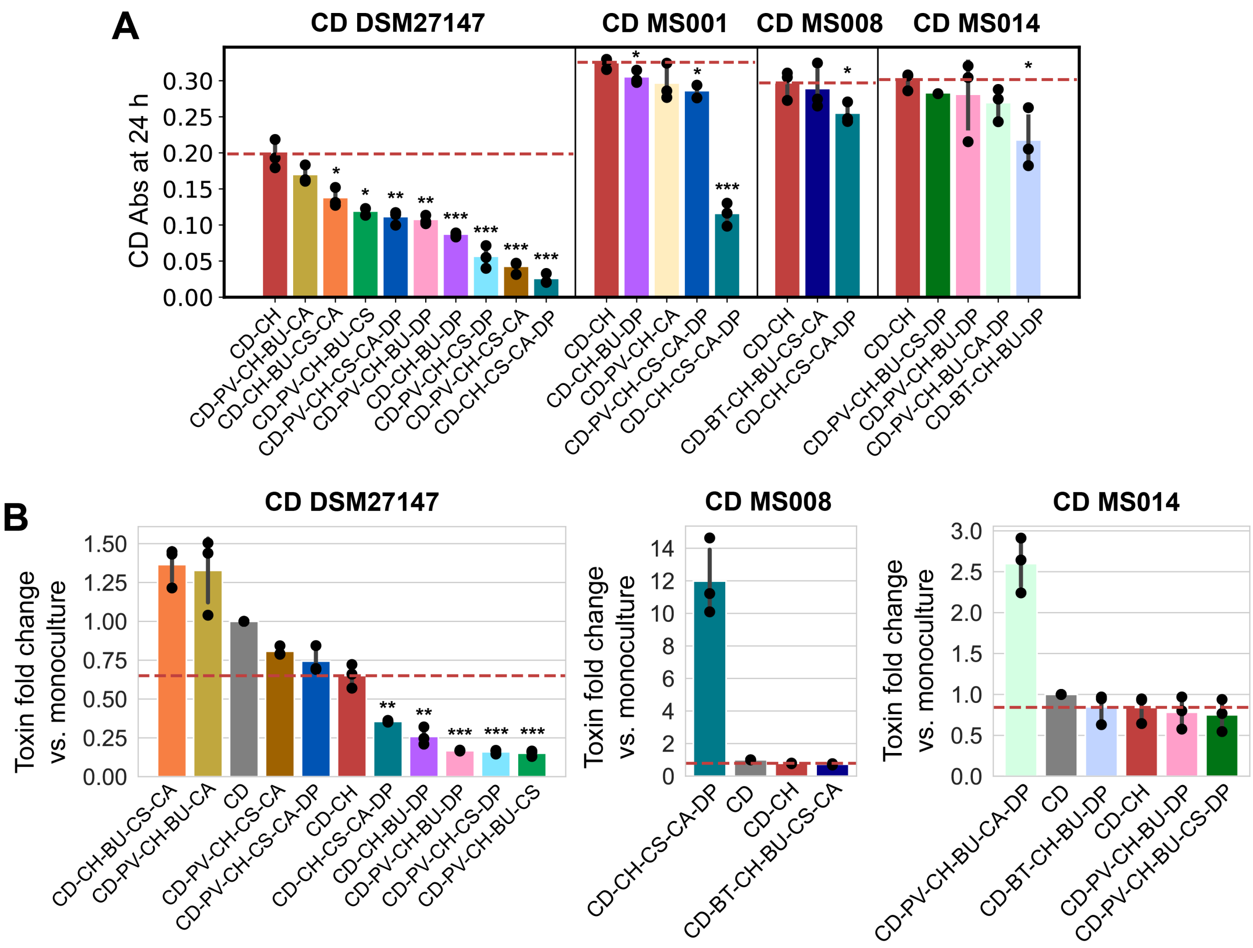


**Supplementary Figure 13. Middle-richness communities containing *C. hiranonis* with similar or better inhibitory activity against *C. difficile* compared to *C. hiranonis* alone**. **a-b**, *C. difficile* absolute abundance at 24 h (**a**) and toxin fold change compared to monoculture as quantified by ELISA (**b**) in different community combinations when grown in the mixed carbohydrates media. Different colors indicate different communities. Horizontal dashed lines indicate *C. difficile* absolute abundance in the CD-CH pair (**panel a**) or toxin fold change in the CD-CH pair compared to monoculture (**panel b**). Asterisks above the bars indicate the *p*-value from unpaired *t*-test of *C. difficile* absolute abundance or toxin fold change in specific middle-richness community and *C. difficile* absolute abundance or toxin fold change in CD-CH pair: * indicates p<0.05, ** indicates p<0.01, *** indicates p<0.001.

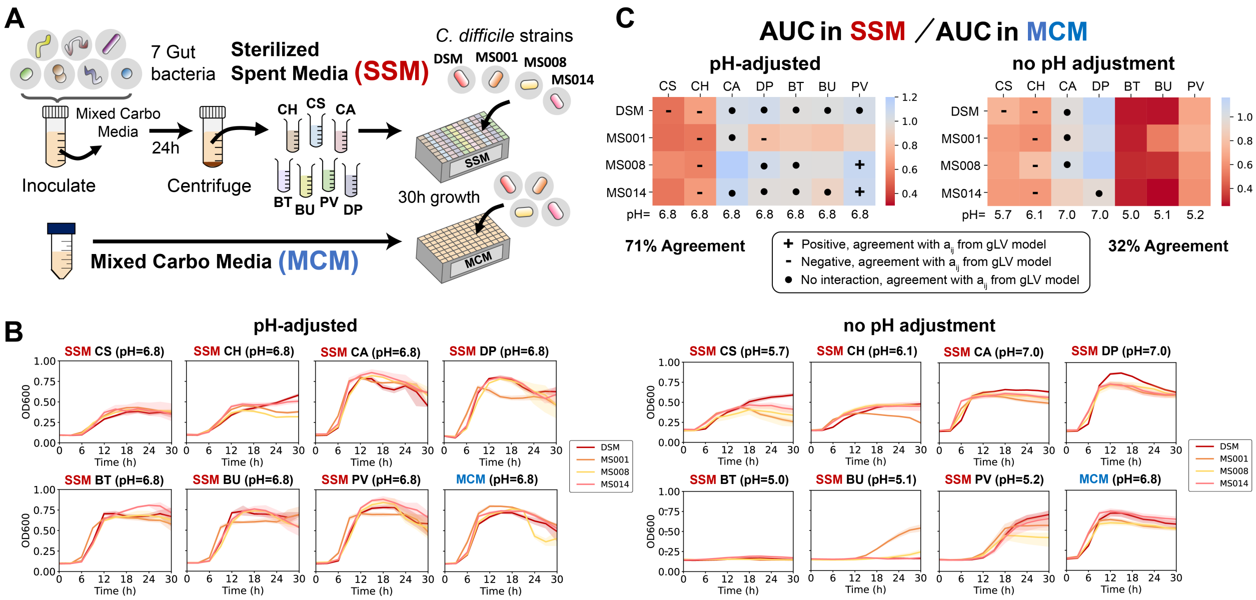

**Supplementary Figure 14. Growth of different *C. difficile* strains in the sterilized spent media of the human gut bacteria. a**, Schematic of the experimental workflow for studying the growth of *C. difficile* strains in the spent media of the gut bacteria. Each of the gut species was grown anaerobically in mixed carbohydrates media for 24 h. The cultures were centrifuged, and the supernatants were filter sterilized. The *C. difficile* strains were grown anaerobically in each of the sterilized spent media (SSM), and the mixed carbohydrates media (MCM) as a control. **b**, $OD_{600}$ measurement of four *C. difficile* strains in the SSM of each gut bacteria or MCM for 30 hours. Data were shown as mean and 95% c.i. (shading), n = 3 biological replicates. pH values for each of the SM and CM were shown in the titles of the growth curves. The figures on the left show growth on the pH-adjusted SSM and MCM, whereas the figures on the right show growth on the SSM and MCM without pH adjustment. **c**, Heatmap of the Area Under the Curve (AUC) upon growth in specific SSM divided by AUC upon growth in MCM, extracted from 30 h of monoculture growth data. The AUC is calculated from the mean of 3 biological replicates. The left heatmap is AUC from growth in SSM with pH adjusted to the pH of MCM whereas the right heatmap is AUC from growth in SSM without pH adjustment. The symbols indicate agreement with the interaction parameters (aij) from the gLV model. The results were in agreement if the AUC in SSM per AUC in MCM > 1 and $\alpha_{ij} > 0.1$ (positive interaction), or AUC in SSM per AUC in MCM < 1 and $\alpha_{ij} < -0.1$ (negative interaction). or 0.9 < AUC in SSM per AUC in MCM < 1.1 and $|\alpha_{ij}| = 0.1$ (no interaction).

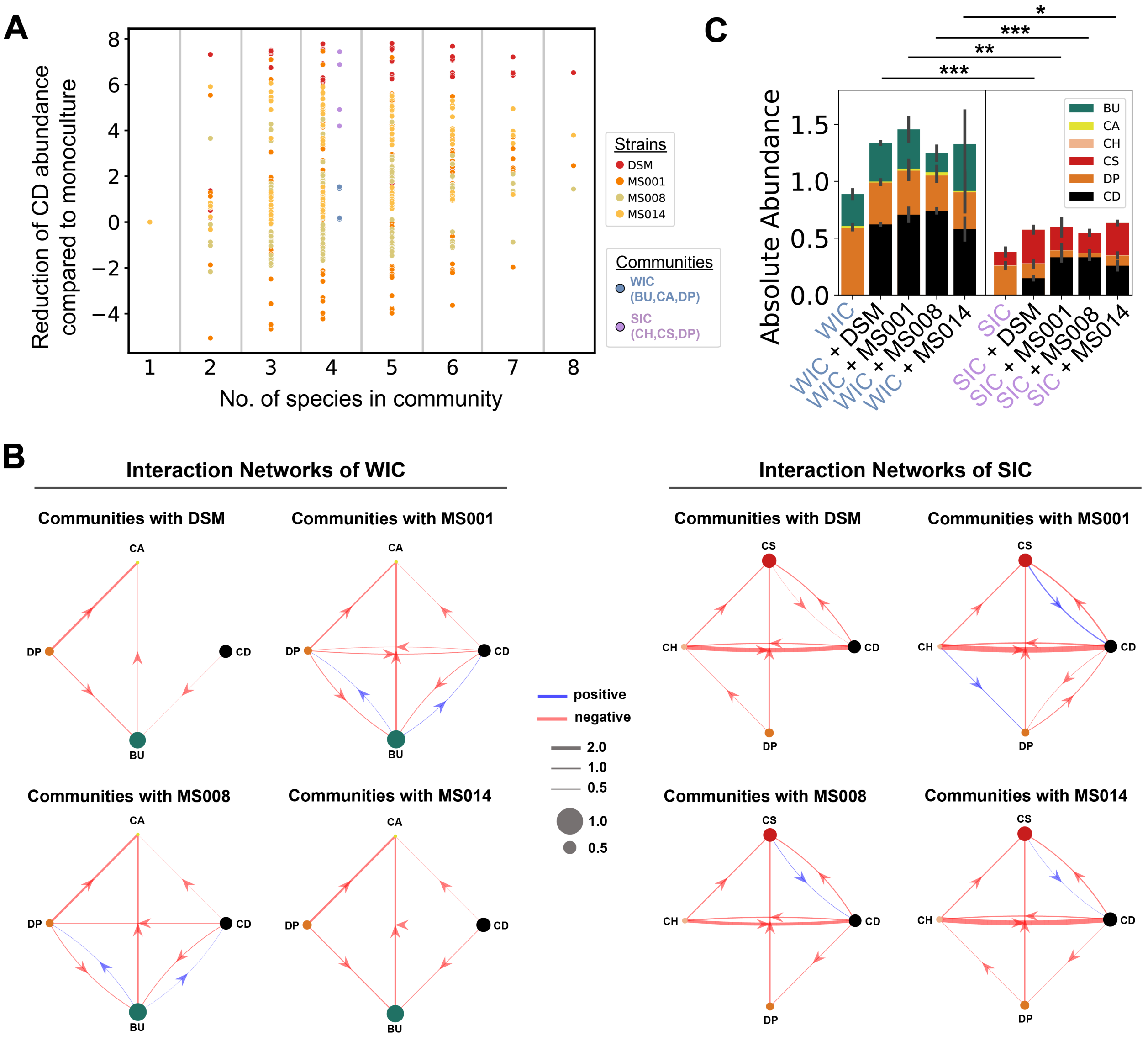


**Supplementary Figure 15. Model could distinguish strong inhibitory community (SIC) and weak inhibitory community (WIC) against *C. difficile*. a**, Extent of *C. difficile* inhibition by all possible 2–8-member community combinations as predicted by the gLV model trained on the mixed carbohydrates media. Two communities were selected for follow-up analyses: Weak Inhibitory Community (WIC) which consists of BU, CA, and DP, and Strong Inhibitory Community (SIC) which consists of CH, CS, and DP. **b**, Inferred interspecies interaction networks between the gut species in the WIC (left) or SIC (right) and each of the representative *C. difficile* strains in the mixed carbohydrates media. Node size represents species carrying capacity in monoculture (mean of all biological replicates) and edge width represents the magnitude of the interspecies interaction coefficient ($a_{ij}$). Edges shows parameters whose absolute values were significantly constrained to be non-zero based on the Wald test. **c**, Species absolute abundance experimentally measured in the absence and presence of different *C. difficile* strains in the mixed carbohydrates media after growth for 24 h. Each bar represents the average absolute abundance of each species (n=3). Error bars represent the standard deviation of 3 biological replicates.

Asterisks above the bars indicate the *p*-value from unpaired *t*-test of the *C. difficile* absolute abundance: * indicates $p<0.05$, ** indicates $p<0.01$, *** indicates $p<0.001$.

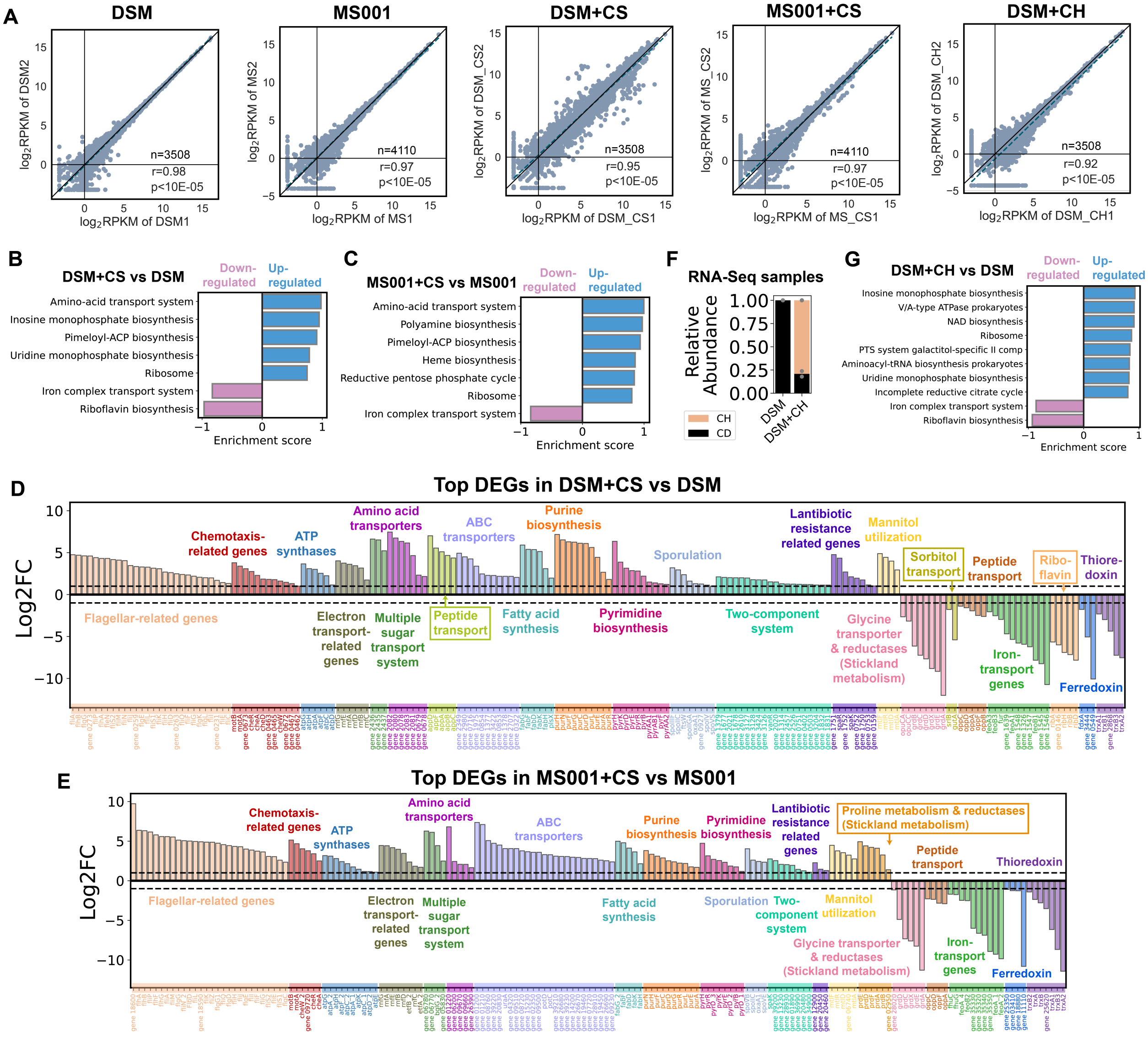


**Supplementary Figure 16. Transcriptomics analysis on *C. difficile* DSM and MS001 in monocultures and cocultures with *C. scindens* or *C. hiranonis*. a**, Scatter plot showing the correlation between the Reads Per Kilobase Million (RPKM) of the two biological replicates in each sample. Blue dashed line indicates the linear regression between the coefficients of the two communities. Pearson's correlation coefficient (*r*) and *p*-values, together with the number of data points (n) are shown. Pearson's *r* and *p*-values were computed using the pearsonr from the scipy package in Python. **b-c**, Enriched gene sets in *C. difficile* DSM27147 (**b**) and MS001 (**c**) grown in the presence of *C. scindens* compared to *C. difficile* MS001 grown in monoculture. All gene sets with significant enrichment scores from GSEA are shown. Gene sets are defined using KEGG modules. **d-e**, Top differentially expressed genes (DEGs) of two different *C. difficile* strains in the presence of *C. scindens*. Bar plot of the log-transformed fold changes of selected highly differentially expressed genes or operons of *C. difficile* DSM27147 (**d**) and MS001 (**e**) in the presence of *C. scindens*. Horizontal dashed lines indicate a 2-fold change. Genes or operons where the text is boxed indicate those that were not highly differentially expressed in the other strain. **f**, Stacked bar plot of the composition of the *C. difficile* monoculture, and *C. difficile*-*C. hiranonis* coculture subjected to RNA-Seq as determined by 16S sequencing. **g**, Enriched gene sets in *C. difficile* DSM27147 grown in the presence of *C. hiranonis* compared to *C. difficile* DSM27147 grown in monoculture. All gene sets with significant enrichment scores from GSEA are shown. Gene sets are defined using KEGG modules.

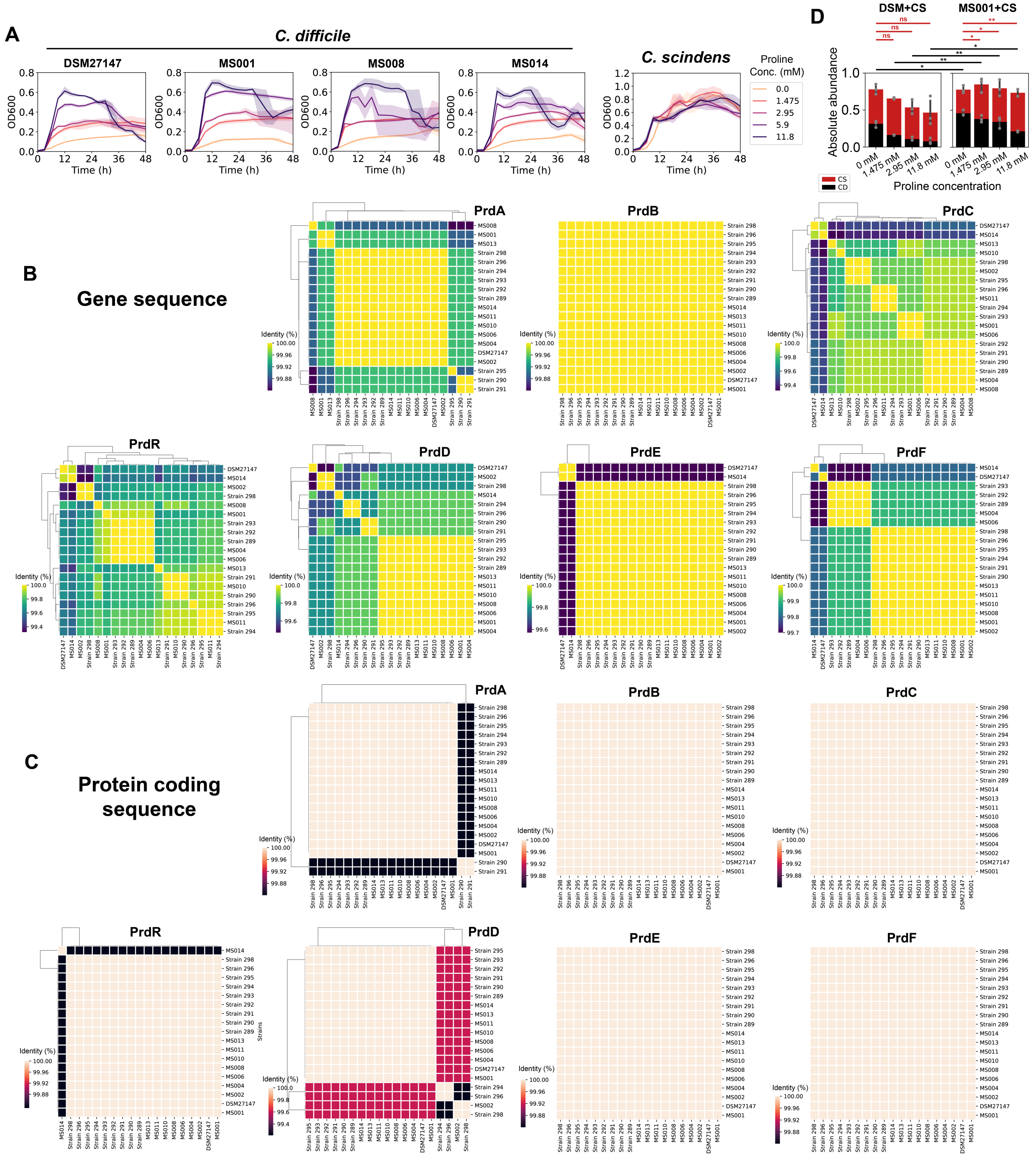


**Supplementary Figure 17. Proline utilization of *C. difficile* isolates. a**, Growth of *C. difficile* and *C. scindens* in monoculture and coculture in media supplemented with different proline concentrations. The plots show time-course $OD_{600}$ measurements of different *C. difficile* strains and *C. scindens* in the mixed carbohydrates media with varying concentrations of proline. Data were shown as mean and 95% c.i. (shading), n = 3 biological replicates. **b-c,** Biclustering heatmap of the percent identity of genes in the *prd* operon (**b**) and the protein coding sequences of the genes in the *prd* operon (**c**) between different *C. difficile* strains. **d**, Stacked bar plot of the absolute abundance of *C. difficile* DSM27147 or MS001 grown with CS in media supplemented with different proline concentrations. Each bar represents the average absolute abundance of each species, and the error bars represent s.d. (n=3). Black asterisks above the bars indicate the *p*-value from unpaired *t*-test of *C. difficile* absolute abundance between DSM27147 and MS001 strain grown in co-culture with *C. scindens* under the same proline concentration, whereas red asterisks indicate the *p*-value from unpaired *t*-test of *C. scindens* absolute abundance grown in co-culture with *C. difficile* under different proline concentration: * indicates $p<0.05$, ** indicates $p<0.01$, ns indicates not significant.

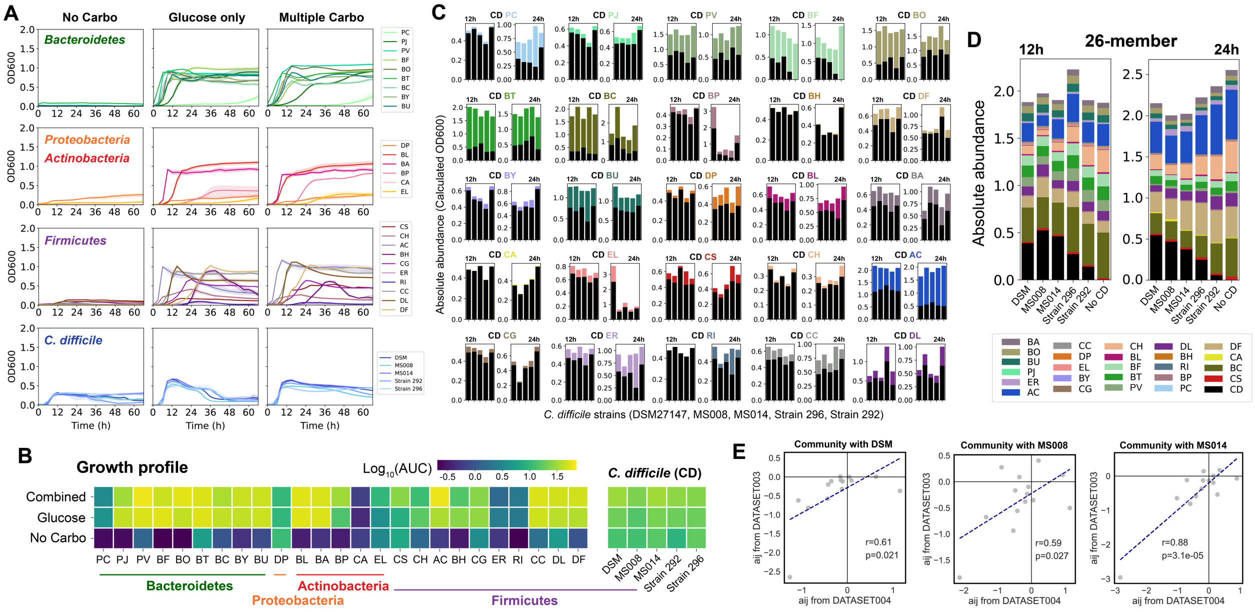


**Supplementary Figure 18. Monoculture growth of different *C. difficile* strains and 25 gut commensal bacteria. a,** OD$_{600}$ measurement of *C. difficile* strains and gut bacteria grown anaerobically in DM29 defined media without any carbohydrate source, supplemented with glucose (glucose media), and supplemented with multiple carbohydrate sources (mixed carbohydrates media). Data were shown as mean and 95% c.i. (shading), n = 3 biological replicates. **b**, Heatmap of the average integral OD$_{600}$ or the Area Under the Curve (AUC) extracted from monoculture growth data of different *C. difficile* strains and gut species after 66 h of growth. **c-d**, Stacked bar plot of the composition of pairwise communities (**c**) and full 26-member community (**d**) containing *C. difficile* and human gut bacteria grown in the mixed carbohydrates media. The *y*-axis represents the absolute abundance or the calculated OD$_{600}$, whereas the *x*-axis represents communities with different *C. difficile* strains (left to right: DSM27147, MS008, MS014, Strain 296, Strain 292). Two plots were shown for each community, where the left plot is the abundance at 12 h whereas the right plot is the abundance at 24 h of growth. Each bar represents the average absolute abundance of each species (n=3). **e,** Correlations of the growth interaction parameters ($a_{ij}$) between *C. difficile* (Strain DSM, MS008, and MS014) and gut bacteria based on gLV model fit from DATASET003 and DATASET004 (**Table S8**). Pearson's correlation coefficient (*r*) and *p*-values are shown, which were computed using the pearsonr from the scipy package in Python.

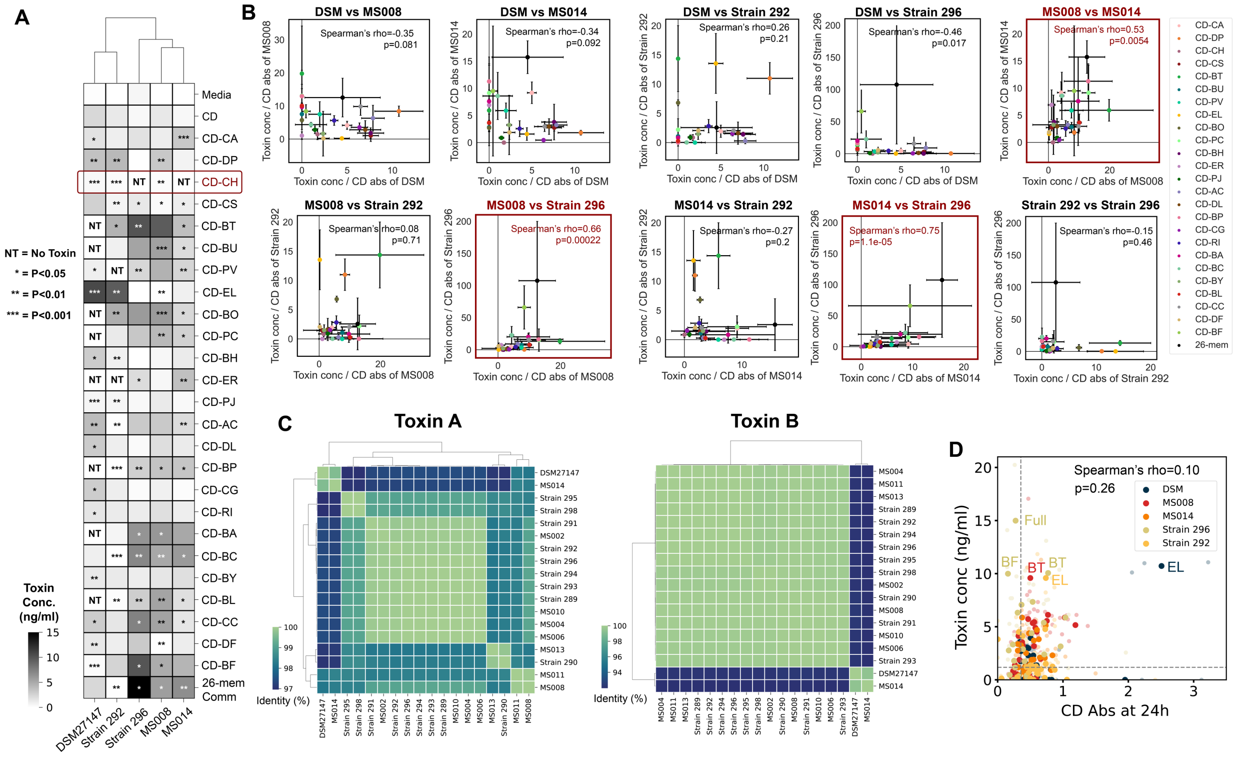

**Supplementary Figure 19. Variation in toxin production between *C. difficile* strains in communities. a,** Heatmap of toxin production of different *C. difficile* strains when grown in pairwise and 26-member communities with human gut bacteria in the mixed carbohydrates media (n=3). Asterisks on the heatmap indicate the *p*-value from unpaired *t*-test of the toxin production in cocultures compared to *C. difficile* monocultures: * indicates p<0.05, ** indicates p<0.01, *** indicates p<0.001, NT indicates No Toxin (toxin concentration = 0 ng/ml). **b,** Correlations of toxin yield between different *C. difficile* strains in communities. Each data point is the toxin yield of *C. difficile* (either strain shown in the x-axis or strain shown in the y-axis) when grown in a pairwise or full 26-member community (indicated with different colors). Error bars indicate standard deviation (n=3). Spearman's rho and *p*-values are shown, which were computed using the spearmanr from the scipy package in Python. Plots outlined with red color marked statistically significant positive correlations. **c**, Biclustering heatmap of the percent identity of toxin A and toxin B amino acid sequence between different *C. difficile* strains. **d**, Scatter plot of *C. difficile* absolute abundance and toxin concentration at 24 h. Horizontal and vertical dashed lines indicate the average toxin concentration and absolute abundance of *C. difficile* in monocultures, respectively. Solid data points indicate the mean of the biological replicates (transparent data points). The colors indicate which *C. difficile* strain was present in the community. Spearman's rho and *p*-value are shown.

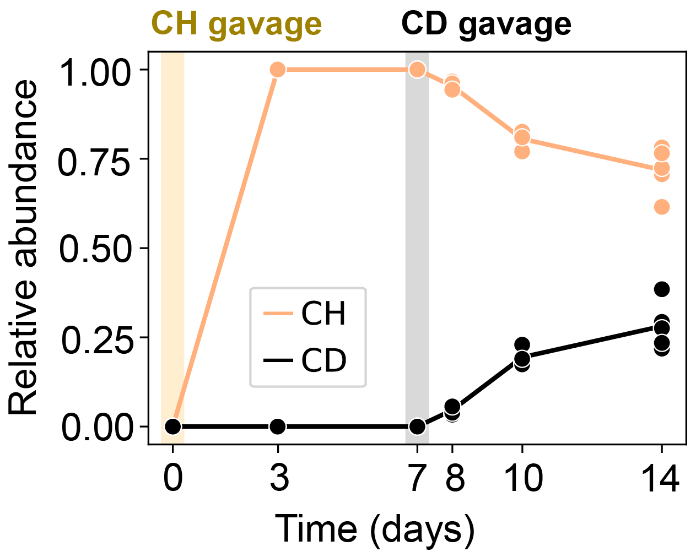


**Supplementary Figure 20. Composition of the fecal (day 0 to 10) and cecal (day 14) content over time**. Relative abundance is determined through 16S sequencing. Datapoints represent individual mice, and the line represents the average of all mice in the group.
